# Design of allosteric modulators that change GPCR G protein subtype selectivity

**DOI:** 10.1101/2024.11.20.624209

**Authors:** Madelyn N. Moore, Kelsey L. Person, Abigail Alwin, Campbell Krusemark, Noah Foster, Caroline Ray, Asuka Inoue, Michael R. Jackson, Michael J. Sheedlo, Lawrence S. Barak, Ezequiel Marron Fernandez de Velasco, Steven H. Olson, Lauren M. Slosky

## Abstract

G protein-coupled receptors (GPCRs), the largest family of drug targets, can signal through 16 subtypes of Gα proteins. Biased compounds that selectively activate therapy-relevant pathways promise to be safer, more effective medications. The determinants of bias are poorly understood, however, and rationally-designed, G protein-subtype-selective compounds are lacking. Here, using the prototypical class A GPCR neurotensin receptor 1 (NTSR1), we find that small molecules binding the intracellular GPCR-transducer interface change G protein coupling by subtype-specific and predictable mechanisms, enabling rational drug design. We demonstrate that the compound SBI-553 switches NTSR1 G protein preference by acting both as a molecular bumper and a molecular glue. Structurally, SBI-553 occludes G protein binding determinants on NTSR1, promoting association with select G protein subtypes for which an alternative, shallow-binding conformation is energetically favorable. Minor modifications to the SBI-553 scaffold produce allosteric modulators with distinct G protein subtype selectivity profiles. Selectivity profiles are probe-independent, conserved across species, and translate to differences in *in vivo* activity. These studies demonstrate that G protein selectivity can be tailored with small changes to a single chemical scaffold targeting the receptor-transducer interface and, as this pocket is broadly conserved, present a strategy for pathway-selective drug discovery applicable to the diverse GPCR superfamily.

## INTRODUCTION

G protein-coupled receptors (GPCRs) are the largest family of transmembrane receptors in the human genome and the largest group of targets for clinically tested drugs^1^. GPCRs convert extracellular signals into intracellular responses by activating transducer proteins, including 16 subtypes of Gα G proteins from four families (i.e., Gi/o, Gs, Gq, G12/13) and two β-arrestin proteins. These transducers mediate distinct and, in some cases, opposite effects. A growing body of evidence suggests that different ligands of the same receptor can induce distinct transducer coupling and physiological responses, a phenomenon known as functional selectivity or biased signaling^2,3^. With greater signaling pathway selectivity, biased ligands may serve as a basis for safer, more effective medications^4^.

GPCRs govern most physiological functions through G protein signaling. The discovery of compounds that shunt receptor signaling away from one G protein subtype and toward another will allow not only for the separation of on-target therapeutic and side effects, but also for the redirecting of signaling by receptors responsible for disease^4^. For example, the potential creation of neurokinin receptor ligands that inhibit rather than produce pain, chemokine receptor ligands that prevent rather than promote cancer metastasis, and metabotropic glutamate receptor ligands that suppress rather than induce seizure activity. The molecular basis of biased agonism, however, is not well understood. In some instances, there is a clear connection between the changes in the receptor conformation induced by the binding of a biased ligand and the type of signaling bias observed^5,6^. In other cases, the connection is not obvious^7^. As such, the structure-based and *de novo* molecular design of biased compounds has been met with limited success.

We propose that small molecules that bind the intracellular GPCR-transducer interface can change G protein subtype selectivity by specific and predictable mechanisms. In the intracellular pocket, compounds are positioned to interact directly with transducers. Here, they may serve simultaneously as molecular bumpers and molecular glues, sterically excluding some transducers while promoting association with others. Given the sequence diversity of the portion of the G protein that interacts with the GPCR core, it should be possible for a core-binding ligand to change receptor G protein affinity in a subtype-specific manner, switching that receptor’s preferred G protein, resulting signaling, and physiological effect. As the receptor-transducer interface is both highly receptor-specific in size and character and broadly conserved across the diverse GPCR superfamily, the ability to design compounds that bind this site and confer signaling bias opens a myriad of new avenues for GPCR drug discovery.

As proof-of-concept, we examined the prototypical class A GPCR NTSR1, a receptor for which pathway-specific physiology has been uncovered and an intracellular small molecule allosteric modulator has been identified: SBI-553^8-10^. NTSR1 ligands have been sought-after for the treatment of psychiatric diseases, including schizophrenia and substance use disorders, but development of balanced agonists that activate many transducers is precluded by on-target side effects, including hypothermia^11^. Biased NTSR1 agonists that preferentially activate β-arrestins over G proteins attenuate addiction-associated behaviors without the side effects characteristic of balanced receptor activation^9,12^. Here, we find that SBI-553, in addition to conferring β-arrestin bias, switches NTSR1 G protein subtype preference. We describe the mechanism by which this G protein subtype selectivity is achieved and use this mechanism to design new compounds with distinct selectivity profiles. We find that, at the receptor-transducer interface, SBI-553 promotes association with G protein subtypes that can adopt an alternative, shallow-binding conformation and excludes those G protein subtypes for which this conformation is energetically unfavorable. Leveraging predictive structural models and high-dimensional structure-activity relationship (SAR) studies, we demonstrate that small modifications to the SBI-553 scaffold produce new allosteric modulators that differentially change NTSR1 G protein coupling. These modulators exhibit exquisite selectivity, discriminating both between and within G protein families. G protein selectivity profiles are independent of the activating orthosteric ligand, conserved across receptor species, and translate to differences in efficacy in a rodent model of NTSR1 agonist-induced hypothermia. These findings indicate that G protein-subtype-selective biased allosteric modulators (BAMs) can be rationally designed and provide a roadmap for the discovery of these BAMs for the wide array of GPCR therapeutic targets.

## RESULTS

### Characterization of ligand-directed G protein and β-arrestin activation by the NTSR1

There are four families of Gα proteins, Gi/o, Gq, Gs, and G12/13, which mediate distinct cellular effects. Most GPCRs couple with G proteins from more than one family^13,14^. While NTSR1 most potently activates Gq-family G proteins, it is promiscuous, activating at least 12 G proteins across three families^15^. SBI-553 does not activate Gq and blocks NT-induced Gq activation^9^, but a comprehensive assessment of its effects on other G proteins has not been undertaken. We began by characterizing human NTSR1 signaling following stimulation by: 1) the endogenous ligand NT, 2) the reduced amide NT active fragment mimetic PD149153^16,17^, 3) the intracellular β-arrestin BAM SBI-553^8,9^, and 4) the competitive NTSR1 antagonist/inverse agonist SR142948A^18,19^. Using the TRUPATH BRET2 sensors^15^ in HEK293T cells, we assessed ligand-induced activation of 14 Gα proteins (**Fig 1A, B**). In these assays, as reported^15^, NT exhibited extreme promiscuity. The only G protein family NT failed to activate in this platform was Gs (**Fig 1 C,D**). NT potency was highest at the Gq/11 family G proteins (EC_50_ 1.5-3 nM), but NT was also a potent Gi/o and G12/13 family activator, producing sub 30 nM EC_50_ for all family members.

**Figure 1.**
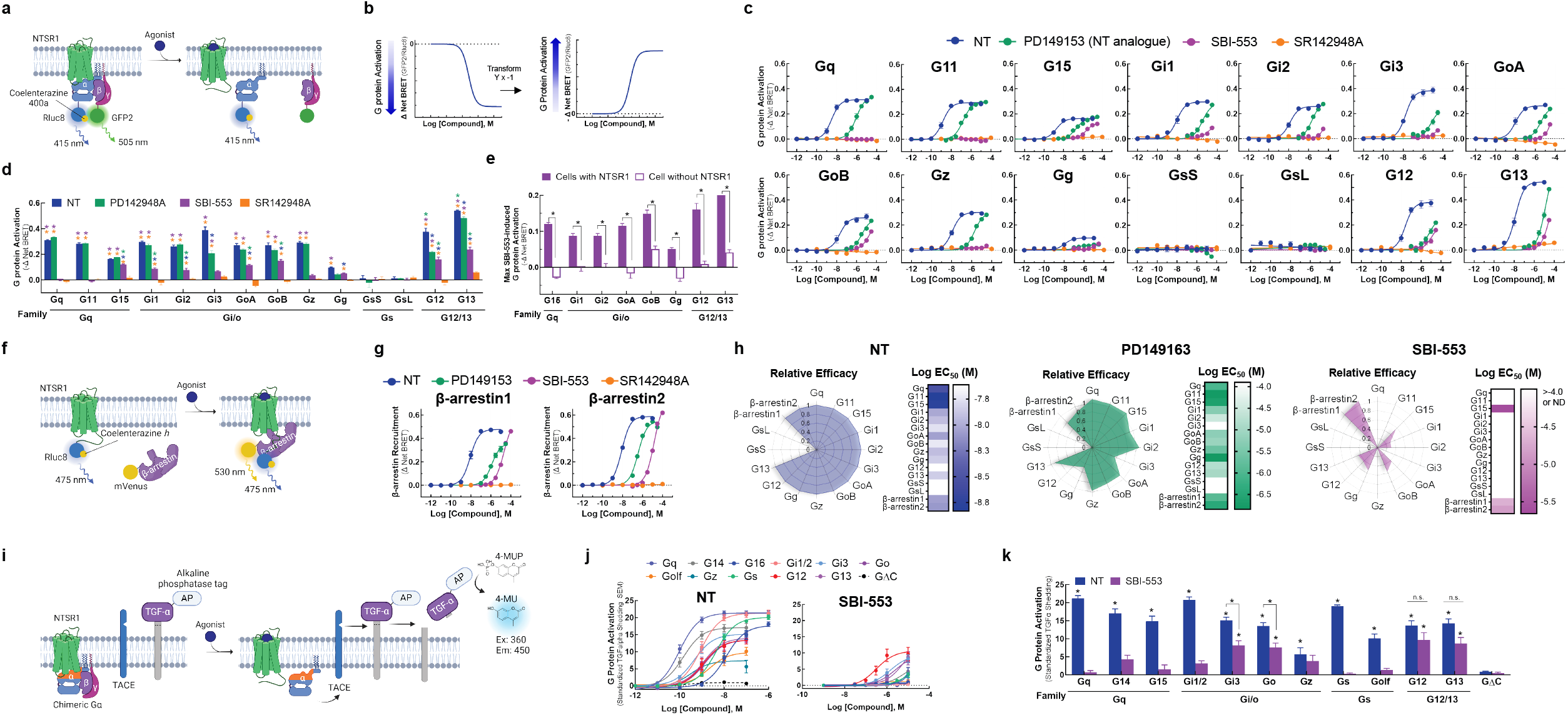
The NTSR1 allosteric modulator SBI-553 exhibits transducer-specific efficacy. Ligand-directed NTSR1 signaling was assessed in HEK293T cells transiently expressing NTSR1 and G protein or β-arrestin activation sensors. **(A-E)** *NTSR1 ligand-induced G protein activation by TRUPATH*. **(A)** Illustration of BRET2-based TRUPATH assay of G protein activation. **(B)** Depiction of TRUPATH data transformation. G protein activation results in reduced BRET as a BRET donor tagged-Gα and a BRET acceptor tagged-Gγ subunit dissociate. Curves were inverted such that G protein activation resulted in upward sloping curves. **(C)** G protein activation was assessed following treatment with the endogenous agonist NT, the NT peptide analog PD149163, the β-arrestin-biased ligand SBI-553, and the orthosteric antagonist SR142948A. **(D)** Maximal ligand-induced G protein activation. Colored asterisks over each bar indicate the treatments from which that compound significantly differed. Treatment vs NT (*), SR142948A (*), PD149163 (*), SBI-553(*). **(E)** Maximal SBI-553-induced G protein activation in cells transiently expressing NTSR1 or an empty control vector. **(F-G)** *NTSR1 ligand-induced β-arrestin recruitment by BRET*. **(F)** Illustration of BRET1-based assay for β-arrestin recruitment. **(G)** β-arrestin1/2 recruitment was assessed following treatment with NT, PD149163, SBI-553, or SR142948A. **(H)** *Summary of compound efficacy and potency*. Radar plots depict the extent of transducer activation relative to NT (i.e., fold change NT E_max_ for each transducer). Heat maps depict ligand potency. **(I-K)** *NTSR1 ligand-induced G protein activation by TGFα shedding*. **(I)** Illustration of TGFα shedding assay for G protein activation. **(J)** Ligand-induced G protein activation. **(K)** Maximal G protein activation following treatment with NT or SBI-553. Asterisks (*) indicate a change from the GΔC negative control, unless otherwise indicated. For curve parameters, sample size, and statistical comparisons, see **Table S1**.

Relative to NT, PD149163 was less potent and exhibited G protein-specific efficacy (**Fig 1C**). While PD142948A fully agonized all Gq/11 family G proteins and some Gi/o family G proteins, it was a partial agonist of Gi3, G12, and G13 (**Fig 1D**). SR142948A neither stimulated NTSR1 G protein activation nor exhibited inverse agonist properties (**Fig 1C**). In line with prior reports^8,9^, SBI-553 did not activate Gq or G11 (**Fig 1C**). SBI-553 did, however, act as a weak partial agonist of other G proteins. While SBI-553 did not show saturating responses within the evaluated concentration range, precluding EC_50_ determination, it stimulated some degree of activation of G15, Gi1, Gi2, GoA, GoB, Gg, G12 and G13 (**Fig 1D**). SBI-553-induced G protein activation was NTSR1-mediated (**Fig 1E**).

We used a BRET1-based assay to characterize ligand-induced recruitment of human β-arrestin1 and 2 to the NTSR1^9^ (**Fig 1F**). NT induced concentration-dependent recruitment of human β-arrestin1 and 2. PD149163 partially agonized β-arrestin1 and fully agonized β-arrestin2. SBI-553 fully agonized both β-arrestin1 and 2, and SR142948A was without effect (**Fig. 1G**). G protein and β-arrestin agonism data were standardized to maximum NT levels and summarized in radar plots (**Fig 1H**). These plots highlight that SBI-553 acts as a weak partial agonist of a subset of G proteins, but not NTSR1’s cognate Gq/11.

To validate SBI-553’s G protein-subtype-specific effects, we used a complementary transforming growth factor-α (TGFα) shedding-based assay of G protein activation^20^ (**Fig 1I**). In this assay, all G protein sensors are based on the same Gq backbone and G protein-subtype specificity is conferred by substitution of the 6 C-terminal amino acids. As in the TRUPATH assay, NT-stimulated NTSR1 was promiscuous; NT-induced NTSR1 to activate all 11 of the G protein sensors, but not a control G protein construct lacking the C-terminus (i.e., GΔC) (**Fig 1J**). SBI-553 acted as a weak but full agonist of G12/13 and a weak, partial agonist of some Gi/o family members (e.g., Go, Gi3) (**Fig 1J, K**). This finding corroborates and extends the previous results, demonstrating that SBI-553 has previously unappreciated G protein agonist activity and that substituting the 6 C-terminal amino acids of Gq is sufficient to bestow this activity.

### Uniform antagonism of NT-induced NTSR1 activation by a competitive antagonist

As SBI-553 and NT can occupy NTSR1 simultaneously, we turned our attention to how SBI-553 modulates NT-induced NTSR1 transducer activation. To verify our ability to generate and identify dose-response curve (DRC) families characteristic of competitive antagonism (**Fig 2A,B**), we assessed the effect of SR142948A pretreatment on NT-induced transducer activation. SR142958A pretreatment produced uniform, concentration-dependent blockade of NT-induced β-arrestin recruitment (**Fig 2C**) and G protein activation, regardless of Gα subtype (**Fig 2D**). In line with a competitive mechanism, SR142948A increased NT’s apparent EC_50_ in a concentration-dependent manner for all transducers (**Fig 2E**). These data showcase the effects of a competitive antagonist that indiscriminately blocks NT-induced transducer activation.

**Figure 2.**
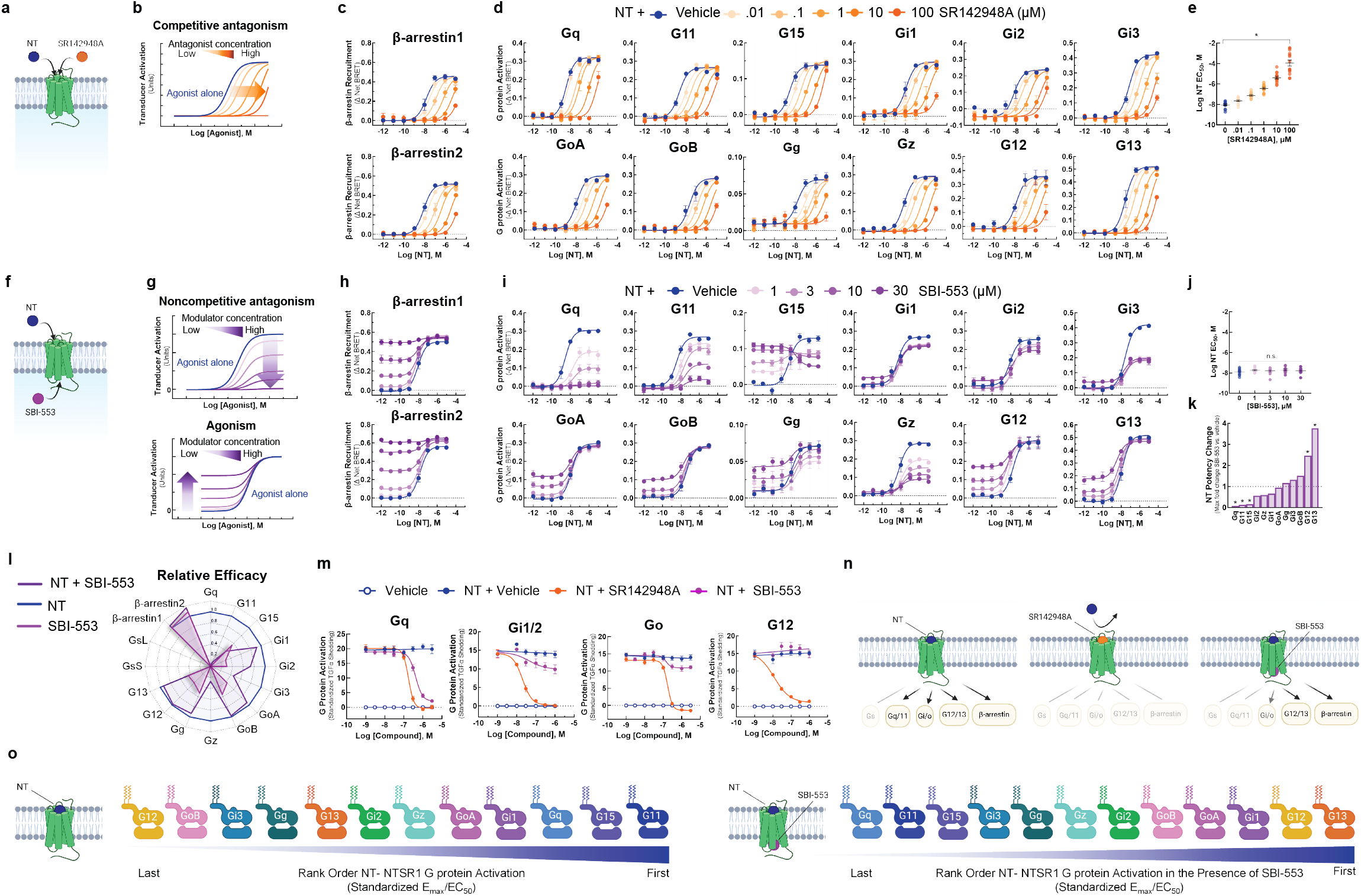
Unlike competitive antagonist SR142948A, SBI-553 biases NT-NTSR1 signaling away from Gq/11 and toward both β-arrestin recruitment and alternative G protein activation. **(A-E)** *SR142948A uniformly competitively antagonizes NT-induced NTSR1 transducer activation*. **(A)** Illustration of NT and SR142948 competing to occupy the same binding site on NTSR1. **(B)** Characteristic agonist dose-response curve (DRC) shifts in the presence of increasing concentrations of a competitive antagonist. **(C)** NT-induced β-arrestin1/2 recruitment to the NTSR1 was assessed by BRET in the absence and presence of SR142948A. **(D)** NT-induced G protein activation was assessed by TRUPATH in the absence and presence of SR142948A. **(E)** NT DRC EC_50_ values for the 12 G proteins evaluated in D in the presence of increasing concentrations of SR142948A. **(F-L)** *SBI-553 exerts transducer-specific effects on NT-induced NTSR1 transducer activation*. **(F)** Illustration of NT and SBI-553 binding distinct sites on the NTSR1. **(G)** (Top) Characteristic agonist DRC shifts in the presence of a noncompetitive antagonist. (Bottom) Characteristic agonist DRC shifts in the presence of an allosteric agonist. **(H)** NT-induced human β-arrestin1/2 recruitment to the NTSR1 in the absence and presence of SBI-553. **(I)** NT-induced G protein activation was assessed by TRUPATH in the absence and presence of SBI-553. **(J)** NT DRC EC_50_ values for the 12 G proteins evaluated in H in the presence of increasing concentrations of SBI-553. **(K)** NT DRC EC_50_ values for each G protein at the maximal SBI-553 concentration for which a sigmoidal curve could be fit. Data are represented as fold-change from vehicle. **(L)** Radar plots depict the extent of activation of each transducer in the presence NT or SBI-553 alone and both NT and SBI-553 in combination. Fold-change values are standardized to NT E_max_ for each transducer. **(M)** *Effect of SR142948A and SBI-553 on NT-induced G protein activation, second assay validation*. NT-induced G protein activation was assessed by TGFα shedding. NT-induced activation of Gq, Gi1/2, Go, and G12 sensors in the presence of SR142948A, SBI-553, or vehicle. **(N)** Illustration of NTSR1 G protein activation following application of NT alone and in the presence of SR149163 or SBI-553. **(O)** Rank order of NT-NTSR1 G protein preference in the absence and presence of SBI-553. For curve parameters, N, and statistical comparisons, see **Table S2**.

### Biasing of NT-induced NTSR1 signaling by the intracellular allosteric modulator SBI-553

In contrast to the action of SR142948A, SBI-553 exhibited allosteric effects that were transducer dependent. From its allosteric location (**Fig 2F**) SBI-553 produced two different activities depending on the transducer evaluated: 1) non-competitive antagonism, as evidenced by a reduction in the height of the NT DRC, and 2) agonism, as evidenced by a rising NT DRC baseline (**Fig 2G**). SBI-553 stimulated β-arrestin recruitment to NTSR1 alone and permitted NT-induced β-arrestin recruitment (**Fig 2H**). In combination with NT, in the TRUPATH assay, SBI-553 fully antagonized some G proteins (i.e., Gq, G11), partially antagonized others (i.e., Gi1, Gi2, Gi3, Gz), and was permissive of NT-induced activation of others (i.e., GoA, GoB, G12, G13) (**Fig 2I**). Unlike a competitive antagonist, SBI-553 did not exert consistent effects on NT EC_50_ values across transducers (**Fig 2J**). SBI-553 increased NT potency at G12 and G13, decreased NT potency at Gq, 11, and 15, and did not significantly change NT potency at other G proteins (**Fig 2K**). Notably, the G proteins for which SBI-553 continued to permit NT-induced signaling were the same as those at which SBI-553 acted as a weak agonist (**Fig 2L**).

To determine whether SBI-553’s G protein-specific effects were consistent across assays, we assessed the ability of SBI-553 to antagonize NT-induced activation of select G proteins in the TGFα shedding-based G protein activation assay. In line with the TRUPATH results, SBI-553 fully antagonized Gq, partially antagonized Gi1/2, and was permissive of NT-induced Go and G12 activation (**Fig 2M**). Together, these data suggest that SBI-553 antagonizes NT-induced activation of some G proteins but not others. By doing so, SBI-553 biases NTSR1 not only toward β-arrestin recruitment, but also toward noncanonical NTSR1 G protein signaling (**Fig 2N**), changing NTSR1’s most preferred G proteins from those of the Gq/11 family to the G12/13 family (**Fig 2O**).

### SBI-553 prevents NTSR1 coupling to select G proteins via a β-arrestin-independent mechanism

SBI-553 antagonism of NT-NTSR1 G protein activation may arise either from a reduction in the formation of NTSR1-G protein complexes or from the stabilization of signaling incompetent NTSR1-G protein complexes, as proposed^10,21^. To determine whether SBI-553 permits or blocks NTSR1 G protein coupling, we turned to a BRET1-based assay of mini-G protein recruitment (**Fig 3A**). Mini-G proteins are modified G proteins that form more stable GPCR complexes, permitting monitoring of complex association by BRET^22^. NT stimulated recruitment of mini-Gq, Gi1, Gs, Go and G12 to NTSR1 (**Fig S1**). In line with SBI-553 inhibiting NTSR1-Gq coupling, SBI-553 fully antagonized NT-induced mini-Gq recruitment (**Fig 3B**). SBI-553 partially antagonized NT-induced mini-Gi1 and Gs recruitment, had little effect on Go recruitment, and increased G12 recruitment, a finding that may reflect an increase in the number of NTSR1-G12 complexes or a conformational change that brings BRET sensors into closer proximity. These data suggest that SBI-553 selectively fully antagonizes NTSR1 Gq activation by preventing NTSR1-Gq coupling. β-arrestins desensitize GPCRs, in part, by occupying the receptor core and preventing G protein coupling^23^. SBI-553 antagonism of NTSR1-Gq coupling may occur because SBI-553 binding is incompatible with Gq binding or because SBI-553 stimulates β-arrestin recruitment to NTSR1, and β-arrestin, in turn, prevents Gq binding. To address this question, we evaluated the ability of SBI-553 to antagonize NT-induced NTSR1-Gq complex formation in β-arrestin1/2-null HEK293 cells^24^. SBI-553 was equally potent and fully blocked NTSR1-Gq recruitment in both β-arrestin1/2-null cells and their parental control line (**Fig 3C**), suggesting that β-arrestins are not necessary for SBI-553’s blockade of NTSR1 Gq coupling.

**Figure 3.**
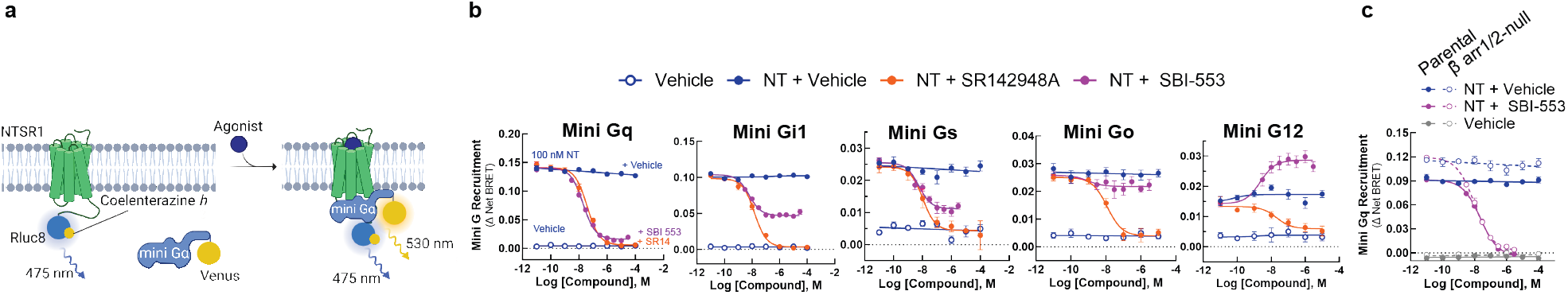
SBI-553 blocks NTSR1 coupling with a subset of G proteins by a β-arrestin-independent mechanism. **(A)** Illustration of BRET1-based assay for mini-G recruitment. **(B)** NT (100nM)-induced Mini-Gq, Gi1, Gs, Go, or G12 recruitment to the NTSR1 was assessed in the presence of Vehicle (HBSS with 0.5%HP-β-cyclodextrin), SBI-553 or SR142948A. Controls received Vehicle alone. **(C)** NT induced Mini-Gq recruitment in the presence of SBI-553 was assessed in β-arrestin1/2-null HEK293 cells and their parental control line. Cells were pretreated with either SBI-553 or its vehicle prior to application of 100 nM NT or its vehicle. For curve parameters, N, and statistical comparisons, see **Table S3**. For supporting data, see **Figure S1**.

### Identification of the molecular determinants of SBI-553’s G protein selectivity

Rather than being a consequence of β-arrestin recruitment, SBI-553’s antagonism of NTSR1-Gq/11 activation may result from direct or allosteric occlusion of Gq/11’s binding determinants on NTSR1’s intracellular surface. Gα’s associate with the core of GPCRs through their highly variable C-termini. To test whether sensitivity to SBI-553’s antagonism was secondary to the primary structure of the G protein C-terminus, we created chimeric G proteins based on the TRUPATH Gα proteins in which we swapped the 5 C-terminal residues between GoA and Gq (**Fig. 4A**). While SBI-553 was fully permissive of NT-induced GoA activation, SBI-553 antagonized the GoA-Gq chimera (GoA^Gq C-term^, **Fig 4B**). Conversely, SBI-553 acted as a partial rather than a full antagonist of the Gq-GoA chimera (Gq^GoA C-term^, **Fig 4C**). Because the Gq^GoA C-term^ chimera was not fully insensitive to SBI-553 antagonism, we created a larger C-terminal tail swap, removing the 13 C-terminal residues from Gq and replacing them with those from GoA (**Fig. 4D**). This Gq^Δ234-246^ was insensitive to antagonism by SBI-553 at all concentrations of SBI-553 except for the highest (**Fig. 4E**). Together, these data suggest that SBI-553’s G protein subtype selective agonism and antagonism is a result of the different primary structures of the Gα C-termini.

**Figure 4.**
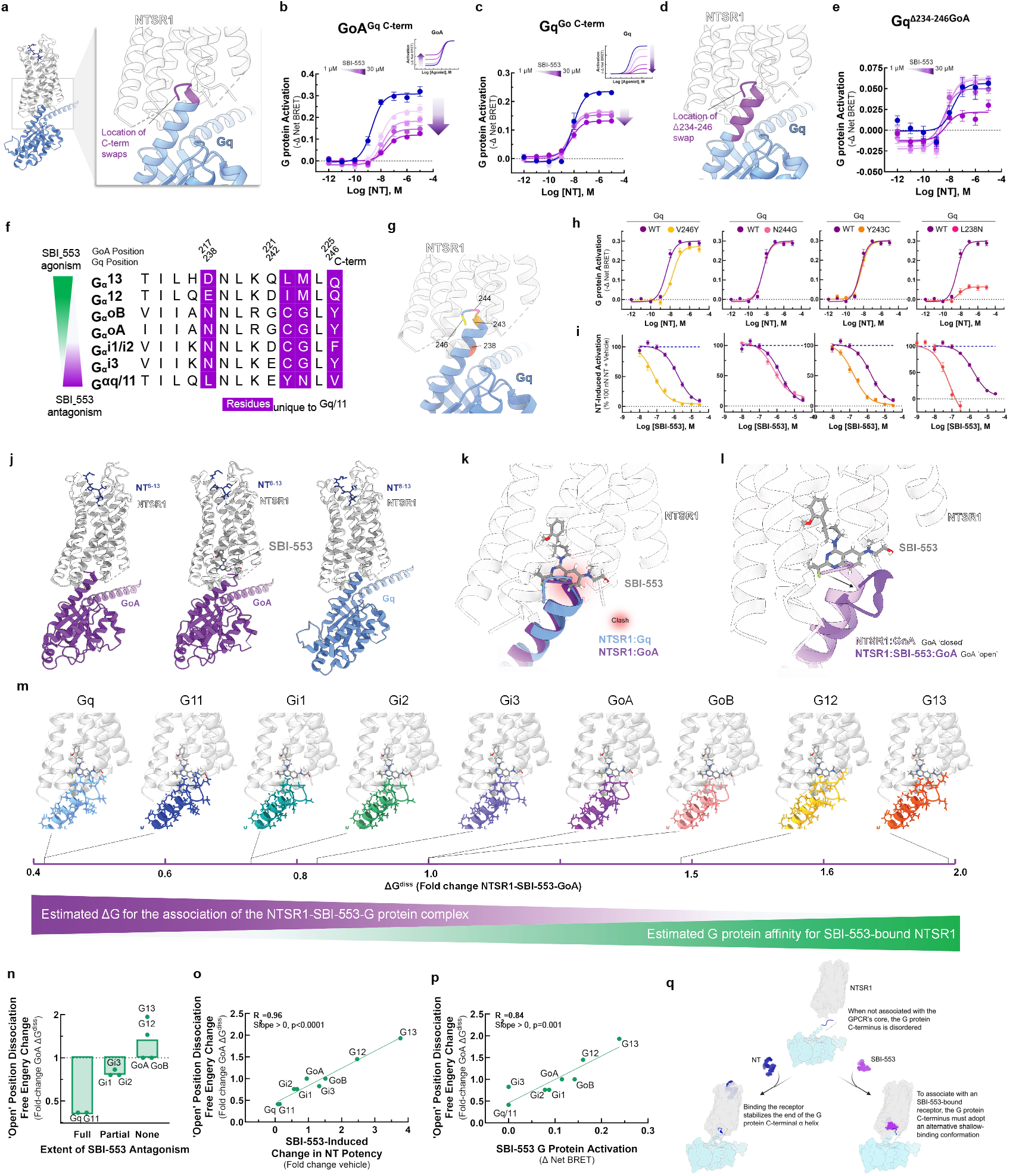
The sensitivity of a Gα protein to SBI-553 antagonism is determined by the ability of its C-terminus to adopt an alternative shallow-binding conformation. **(A-E)** *Sensitivity of G proteins to SBI-553 antagonism can be reversed by exchanging their C-termini*. **(A)** Location of the GoA/Gq 5 C-terminal amino acid residue swap. **(B)** Swapping GoA’s 5 C-terminal amino acids for those of Gq confers sensitivity to SBI-553 antagonism. Inset, effect of SBI-553 on NT-induced activation of WT GoA for reference. **(C)** Swapping Gq’s 5 C-terminal amino acids for those of GoA reduces the antagonist efficacy of SBI-553. Inset, effect of SBI-553 on NT-induced activation of WT Gq for reference. **(D)** Location of Gq/GoA 13 C-terminal amino acid residue swap, Gq^Δ234-246^. **(E)** Swapping Gq’s 13 C-terminal amino acids for those of GoA reduces the antagonist potency and efficacy of SBI-553. **(F)** Alignment of G protein C-termini. **(G-I)** *Single amino acid substitutions on Gq’s C-terminus are insufficient to render Gq permissive of SBI-553*. **(G)** Location of Gq point mutants on the NTSR1-Gq structure. **(H)** NT-induced activation of WT and mutant Gq constructs. **(I)** Effect of Gq mutagenesis on SBI-553’s ability to antagonize Gq activation by 100 nM NT. **(J-L)** *Identification of an SBI-553-induced shallow-binding ‘open’ GoA conformation*. **(A)** Cryo-EM structures showing NTSR1 bound by the NT active fragment and G protein in the absence and presence of SBI-553. (*Left*) NTSR1, NT, mini-GoA (PDB 8FN1). (*Middle*) NTSR1, NT, SBI-553, mini-GoA (PDB 8FN0). (*Right*) NTSR1, NT, mini-Gq (PDB 8FMZ). **(K)** Docking of SBI-553 in the 8FN1 and 8FMZ structures suggests that both GoA and Gq should clash with SBI-553. **(L)** Registration of the 8FN1 and 8FN0 structures illustrates that, in the presence of SBI-553, GoA adopts an alternative ‘open’ conformation to accommodate SBI-553 binding. **(M-P)** *Homology modeling indicates that some NTSR1-SBI-553-’open’ position G protein complexes are more energetically favorable than others*. In silico homology models were created based on the ‘open’ NTSR1-SBI-553-bound GoA conformation by changing the 13 C-terminal amino acids. Free energy of dissociation changes (ΔG^diss^) were calculated, and models are positioned from most stable (right) to least stable (left). **(N)** NTSR1-SBI-553-G protein ‘open’ conformation ΔG^diss^ presented for individual G proteins, categorized by extent of sensitivity to SBI-553 antagonism. **(O)** Correlation of NTSR1-SBI-553-G protein ‘open’ conformation ΔG^diss^ with experimental SBI-553-induced fold change in NT G protein activation potency, as presented in Fig 2K. **(P)** Correlation of NTSR1-SBI-553-G protein ‘open’ conformation ΔG^diss^ with experimental max SBI-553-induced G protein activation, as presented in Fig 2D. **(Q)** Model of SBI-555 G protein subtype selectivity. For curve parameters, sample size, and statistical comparisons, see **Table S4**. For supporting data, see **Figure S2, S3**.

To determine the residues responsible for this change in sensitivity, we aligned selected Gα C-termini, identifying 4 residues unique to Gq/11 (**Fig. 4F**). To evaluate the individual contributions of these residues to SBI-553 antagonism, we generated point mutants that replaced Gq residues in the TRUPATH construct with corresponding GoA residues: V246Y, N244G, Y243C, and L238N (**Fig. 4G**). NT continued to activate all point mutants (**Fig 4H**). SBI-553 IC_50_s were left-shifted for all point mutants relative to the wild-type control, and SBI-553 remained a full antagonist of all constructs (**Fig 4I**). This increase in SBI-553 potency was not explained by a reduction in sensor expression (**Fig S2**). These point mutations unmasked a potency of SBI-553 in the mid-nanomolar range, close to its 50-100 nM binding affinity^25^, possibly by reducing the affinity of the Gq constructs for inactivated NTSR1 and, consequently, binding competition. Critically, no point mutant reproduced the reversal in sensitivity to SBI-553 antagonism seen with the full C-terminal swap.

To understand why changes at single positions were insufficient to reduce Gq’s sensitivity to SBI-553 antagonism, we turned to structural data. SBI-553 binds NTSR1 at an intracellular site in the hydrophobic region of NTSR1’s core where it is positioned to interact with associating transducers ^10,26^. We previously generated structures of NTSR1-NT in complex with mini GoA in the absence (PDB: 8FN1) and presence (PDB: 8FN0) of SBI-553 and of NTSR1-NT in complex with mini Gq in the absence of SBI-553 (PDB: 8FMZ, **Fig 4J**). In the absence of SBI-553, the NTSR1-bound GoA and Gq C-termini exist in a ‘closed’ α-helical state. Superposition of these structures reveals a nearly complete overlap of the GoA and Gq C-termini in the receptor core. Docking SBI-553 at its observed binding site in both structures results in a steric clash (**Fig 4K**). From 8FN0^10^, we see that in the presence of SBI-553, the GoA C-terminus adopts an alternative, shallow-binding ‘open’ conformation. The Go helix tilts 14° toward TM1 and slightly unwinds over the last 5 residues, protruding 4 Å less deeply into the receptor core (**Fig 4L**). G221, at the top of the helix, changes conformation from a phi angle of −64° to +72°. In this ‘open’ orientation, GoA makes van der Waals contact with the hydroxyl side chain, quinazoline C10, and fluorine of SBI-553. NTSR1 undergoes a conformational change, positioning the R167 guanidine to make a strong electrostatic interaction with the quinazoline nitrogen of SBI-533, a nitrogen that SAR indicates is required for SBI-553 binding^8^.

Attempts to generate an NTSR1-SBI-553-Gq structure have not been successful^10^, potentially because NTSR1-SBI-553 binding is incompatible with Gq binding. To predict SBI-553 activity for a range of G proteins, we built *in silico* homology models for Gq, G11, GoB, Gi1, Gi2, Gi3, G12, and G13. In an optimized NT-NTSR1-SBI-553-GoA ‘open’ position structure, we replaced GoA’s 13 C-terminal residues with those of the other G proteins (**Fig S3**). These models reveal multiple structural differences that may increase the energy of this ‘open’ conformation for Gq/11 relative to GoA and provide a rational explanation for SBI-553’s G protein-specific effects. For example, forcing Gq/11’s E242 to adopt a positive phi angle, while allowed, is energetically unfavorable. For both G12 and G13, M223 must adopt a slightly higher energy conformation with gauche interactions, but this small penalty is compensated by close contact with the SBI-553 amino alcohol fragment. In addition to those contacts identified for GoA, G12’s I222 and G13’s L222 both make productive van der Waals contact with SBI-553, further increasing complex stability.

We estimated the free energy required to dissociate each of the NTSR1-SBI-553-’open’ position G protein complexes (ΔG^diss^) (**Fig 4M**). ΔG^diss^ is a measure of thermodynamic stability directly related to NTSR1-G protein affinity. These ‘open’ position ΔG^diss^ values were highest for those G proteins that SBI-553 agonized (i.e., G12, G13, GoA, GoB), were intermediate for those G proteins that SBI-553 partially antagonized (i.e., Gi1, Gi2, Gi3), and lowest for G proteins that SBI-553 fully antagonized (i.e., Gq, G11; **Fig 4N**). Notably, NTSR1-SBI-553-G protein ‘open’ position ΔG^diss^ values accurately predicted SBI-553’s G protein-specific effects on NT potency (R^2^=0.96, **Fig 4O**) and were directly proportional to the extent of SBI-553 agonism (R^2^=0.84, **Fig 4P**). These data suggest that SBI-553 promotes association with Gα proteins that can adopt an alternative shallow-binding conformation and prevents association with subtypes for which this conformation is energetically unfavorable (**Fig 4Q**).

### Discovery of NTSR1 BAMs with distinct G protein selectivity profiles

Based on these models, we undertook SAR studies to determine whether SBI-553’s G protein selectivity profile could be changed by modifying its structure. We hypothesized that SBI-553 analogs would displace or stabilize G protein interactions depending upon the position, size, and direction of substitution. We focused on the fluorine geminal to SBI-553’s cyclopropyl ring and quinazoline C8, C9, and C10 as, according to the model, these regions of the are positioned to interact with G protein C-termini (**Fig 5A**).

**Figure 5.**
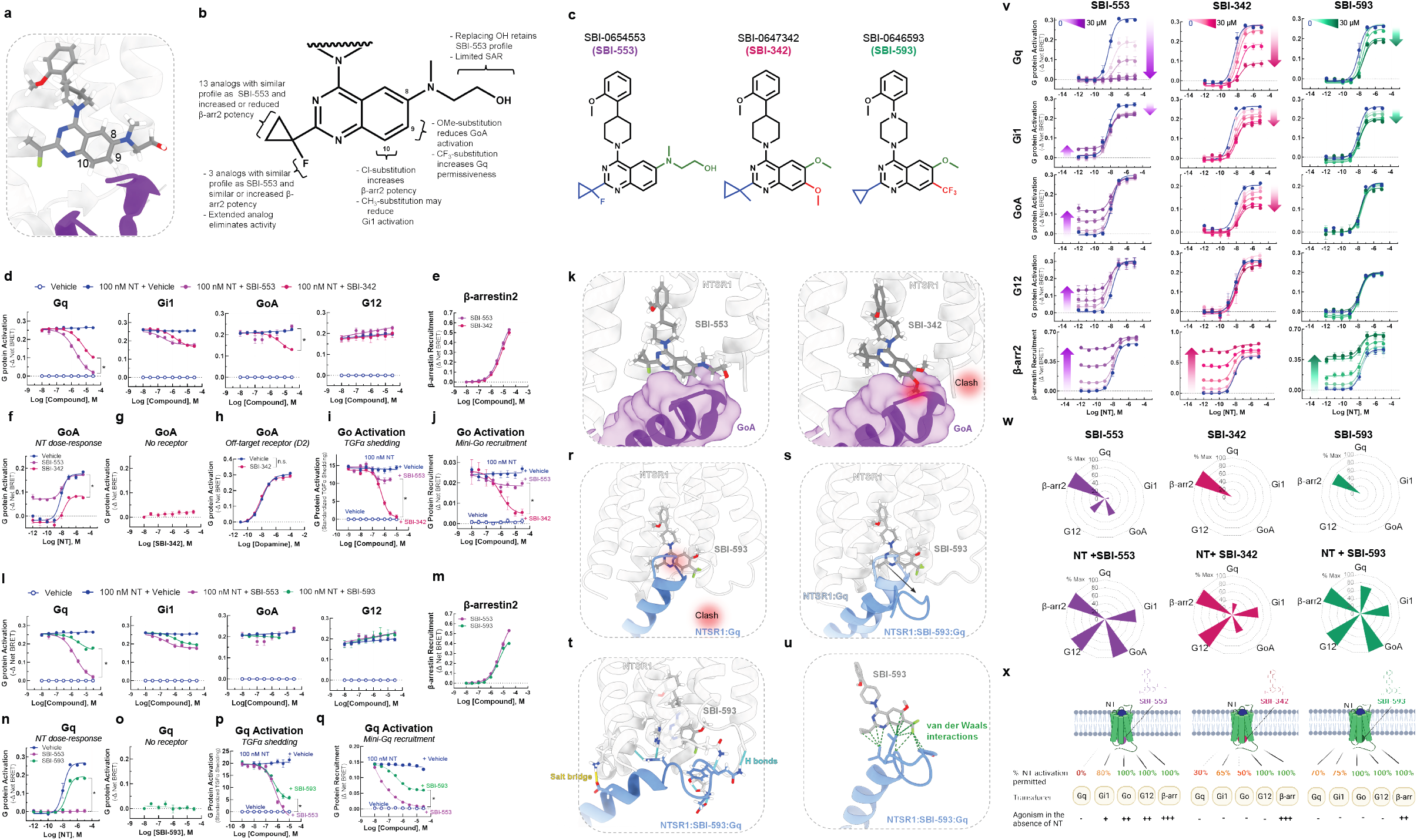
Discovery of SBI-553 analogs with distinct G protein selectivity profiles. **(A)** Position of SBI-553 in the NTSR1 intracellular core. Numbers mark quinazoline C8, C9, and C10. GoA shown in purple. **(B)** Summary of structure-activity relationship (SAR) study findings. **(C)** Structures of SBI-553 and analogs SBI-342 and SBI-593. **(D-K)** *Analog SBI-342 exhibits Go antagonism, not agonism*. **(D)** Screen evaluating SBI-342 antagonism of NT-induced Gq, Gi1, GoA, and G12 activation by TRUPATH. **(E)** Screen evaluating SBI-342 β-arrestin2 agonism by BRET. **(F)** Effect of 30 μM SBI-553 vs SBI-342 across the NT GoA DRC. **(G)** Assessment of SBI-342 on GoA TRUPATH activation sensor activity in HEK293T cells not expressing NTSR1. **(H)** Assessment of SBI-342 (30 μM) on GoA activation stimulated by the Gi/o-coupled dopamine receptor D2. **(I)** Comparison of SBI-553 and SBI-342 antagonism of NT-induced Go activation in the AP-TGFα shedding assay. **(J)** Comparison of SBI-553 and SBI-342 antagonism of NT-induced miniGoA recruitment to the NTSR1 by BRET. **(K)** SBI-553 can co-occupy NTSR1 with GoA’s C-terminus in its ‘open’ position, while the 9-methoxy of SBI-342 clashes with GoA. **(L-U)** *Analog SBI-593 exhibits partial rather than full Gq antagonism*. **(L)** Screen evaluating SBI-593 antagonism of NT-induced Gq, Gi1, GoA, and G12 activation by TRUPATH. **(M)** Screen evaluating SBI-593 β-arrestin2 agonism by BRET. **(N)** Effect of 30 μM SBI-553 vs SBI-593 across the NT Gq DRC. **(O)** Assessment of SBI-593 on Gq TRUPATH activation sensor activity in HEK293T cells not expressing NTSR1. **(P)** Comparison of SBI-553 and SBI-593 antagonism of NT-induced Gq activation in the AP-TGFα shedding assay. **(Q)** Comparison of SBI-553 and SBI-593 antagonism of NT-induced miniGoA recruitment to the NTSR1 by BRET. **(R)** Docking SBI-593 into the SBI-553 binding site in the NTSR1:Gq structure (PDB 8FMZ) indicates a clash between Gq and SBI-593, as for SBI-553. **(S)** Molecular dynamics simulations indicate a repositioning of the Gq C-terminus within the NTSR1 core. **(T)** Interactions between NTSR1 and Gq stabilizing this new position are shown. **(U)** Attractive van der Waals contacts between SBI-593 and the Gq C-terminus its new position are illustrated as dotted green lines. **(V-X)** *SBI-553 analogs have distinct G protein selectivity profiles*. **(V)** NT-induced G protein activation was assessed in the absence and presence of SBI-553, -342, and -593. **(W)** Radar plots depict extent of transducer activation induced by SBI-553, -342, and -593 alone (top) and in the presence of NT (bottom). Values reflect maximal % E_max_ relative to NT (G proteins) or SBI-553 (β-arrestin2). **(X)** Summary of NTSR1 G protein activation following application of NT in the presence of SBI-553, -342, and -593. For curve parameters, N, and statistical comparisons, see **Table S5**. For supporting data, see Figures **S3-9**.

We created a panel of 29 analogs of SBI-553 (**Fig S4**) and assessed them for their ability to antagonize NT-induced Gq, Gi, Go, and G12 activation and to agonize β-arrestin independently (**Fig S5**). A summary of the SAR is shown in **Fig 5B**. We identified compounds with 2-3-fold increased β-arrestin potency and comparable Gq inhibition potency relative to SBI-553, suggesting that these properties can vary independently (**Fig S6**). We also identified compounds that displayed neither G protein antagonism nor β-arrestin agonism, likely due to loss of NTSR1 affinity (**Fig S7**). Most compounds retained β-arrestin agonism. Remarkably, small changes to the SBI-553 scaffold induced pronounced differences in G protein selectivity.

Some of the G protein selectivity effects were predicted by the models, including substitution at quinazoline C8 and C9 leading to disrupted GoA signaling. SBI-0647342 (SBI-342) is an analog of SBI-553 with a dimethoxy-substituted quinazoline and a methyl-substituted cyclopropyl group (**Fig 5C**). In the initial screen, SBI-342 differed from SBI-553 in its ability to antagonize NT-induced GoA activation (**Fig 5D**). Its Gq antagonism was modestly reduced relative to SBI-553, but its β-arrestin agonism was preserved (**Fig 5E**). A comparison of SBI-553 and SBI-342’s effect on NT’s full DRC curve confirmed that SBI-342 lacked GoA agonism and reduced maximal NT GoA activation by ∼60% (**Fig 5F**). The action of SBI-342 was specific to NT-NTSR1, as it did not directly change GoA activation in cells without NTSR1 (**Fig 5G**) and did not impair the ability of other GPCRs to activate Go (**Fig 5H**). In both the TGFα shedding (**Fig 5I**) and Go recruitment (**Fig 5J**) assays, SBI-342 fully blocked NT-induced Go activation.

SAR and modeling data indicate that the change in GoA permissiveness is attributable to SBI-342’s methoxy groups rather than the methyl substitution on the cyclopropyl group, as the fluorine to methyl substitution alone is insufficient to change G protein selectivity (**Fig S8**). A comparison of SBI-553 vs. SBI-342 in the SBI-553 binding pocket of the NTSR1-SBI-553-GoA structure show a clash between the 9-methoxy on the quinazoline of SBI-342 and the C222 backbone carbonyl of GoA, which is absent with SBI-553 (**Fig 5K**). This example highlights the predictive and explanatory value of structure-based BAM design.

Other SBI-553 analogs evaluated in the screen produced unexpected results. For example, compound SBI-0646593 (SBI-593) was more permissive of NT-induced Gq activation than SBI-553. SBI-593 is an analog of SBI-553 with C8 methoxy and C9 trifluoromethyl quinazoline substitution and a cyclopropyl group lacking the fluorine substituent (**Fig 5C**). In the initial screen SBI-593 differed from SBI-553 in its reduced ability to antagonize NT-induced Gq activation (**Fig 5L**). It did not differ from SBI-553 in antagonistic efficacy at the other G proteins evaluated and retained β-arrestin agonism (**Fig 5M**). A comparison of SBI-553 and SBI-593’s effect on NT’s full DRC curve confirmed that SBI-593 only reduced NT Gq activation by 30% (**Fig 5N**), action that was specific to NT-NTSR1 (**Fig 5O**). Consistent with these results, SBI-593 produced partial rather than full inhibition of NT-induced Gq activation in the TGFα shedding (**Fig 5P**) and Gq recruitment (**Fig 5Q**) assays.

SAR and modeling data suggest that the change in Gq permissiveness is attributable to SBI-593’s trifluoromethyl quinazoline substituent rather than the lack of cyclopropyl substitution, as removing SBI-553’s fluorine had no effect on its G protein selectivity profile (**Fig S9**). Docking SBI-593 in the SBI-553 binding pocket of the NTSR1-Gq structure indicates that it should sterically exclude Gq (**Fig 5R**). The preserved Gq activity with SBI-593 indicates that Gq’s C-terminus can adopt multiple binding poses in the NTSR1 core. A series of molecular dynamics simulations performed on the NTSR1-Gq-SBI-593 complex predict that the Gq C-terminal helix tilts out of the way of SBI-593’s bulky trifluoromethyl group toward NTSR1’s transmembrane domain 4 and intracellular loop 1. In these simulations, NTSR1 maintained its tertiary structure, but Gq’s C-terminal helix tilted and unwound to maximize contacts with SBI-593 (**Fig 5S**). In the lowest energy model from these simulations, the interaction of NTSR1, SBI-593, and Gq is stabilized by multiple nonpolar and polar interactions (**Fig 5T**). There are van der Waals interactions between the trifluoromethyl of SBI-593 and Gq that are absent for SBI-553 (**Fig 5U**), which may explain why Gq does not adopt this posture in the presence of SBI-553.

We generated allosteric DRC families for NT with SBI-342, SBI-593, and SBI-553 at Gq, Gi1, GoA, G12, and β-arrestin2 (**Fig 5V**). The maximal extent of NT-induced activation in the presence of each of these BAMs is summarized in radar plots (**Fig 5W**). As is evident from these plots, SBI-553, -342 and -593 exhibit pathway-specific efficacy alone and exert differential effects on NT-induced transducer activation (**Fig 5X**). These studies demonstrate that allosteric modulator G protein selectivity can be changed by making minor modifications to a single scaffold.

### Changes in BAM G protein selectivity alter BAM *in vivo* activity

To determine whether BAMs with different G protein selectivity profiles differ in their *in vivo* activities, we turned to a model of NTSR1 agonist-induced hypothermia. PD149163 is brain penetrant, and its activation of central NTSR1 receptors reduces core body temperature in rats^27^ and mice^9,11,28^. At high doses, SBI-553 fully blocks PD149163-induced hypothermia^9^. As Gq/11 are the G proteins most closely associated with NTSR1’s physiological effects^29,30^ and the only G proteins that SBI-553 fully antagonizes (**Fig 2**), NTSR1-Gq/11 activation may mediate PD149163-induced hypothermia. As such, we hypothesized that SBI-593, which incompletely blocks NTSR1-Gq activation, would be inferior to SBI-553 in its ability to counter PD139163-induced hypothermia in mice.

To test this, we first verified that PD149163-induced hypothermia in mice is mediated by NTSR1. In line with this conclusion, PD149163 produced a dose-dependent reduction in core body temperature in wild-type (WT) mice but not in NTSR1 knockout (KO, NTSR1^-/-^) mice (**Fig 6A,B**). Next, we evaluated SBI-553 and -593 probe and species dependence. Consistent with NT findings, SBI-553 fully and SBI-593 partially antagonized PD149163-induced Gq recruitment to the human NTSR1 (**Fig 6C**). Mirroring data for the human receptor, SBI-553 fully and SBI-593 partially antagonized PD149163-induced Gq recruitment to the mouse NTSR1 (**Fig 6D**). To assess the ability of SBI-553 and SBI-593 to attenuate PD149163-induced hypothermia following systemic administration, mice received SBI-553 (12 mg/kg, i.p.), SBI-593 (12 mg/kg, i.p.), or their respective vehicles 45 min prior to PD149163 (0.15 mg/kg, i.p.) treatment (**Fig 6E**). While SBI-553 attenuated PD149163-induced hypothermia (**Fig 6F**), SBI-593 did not (**Fig 6G**).

**Figure 6.**
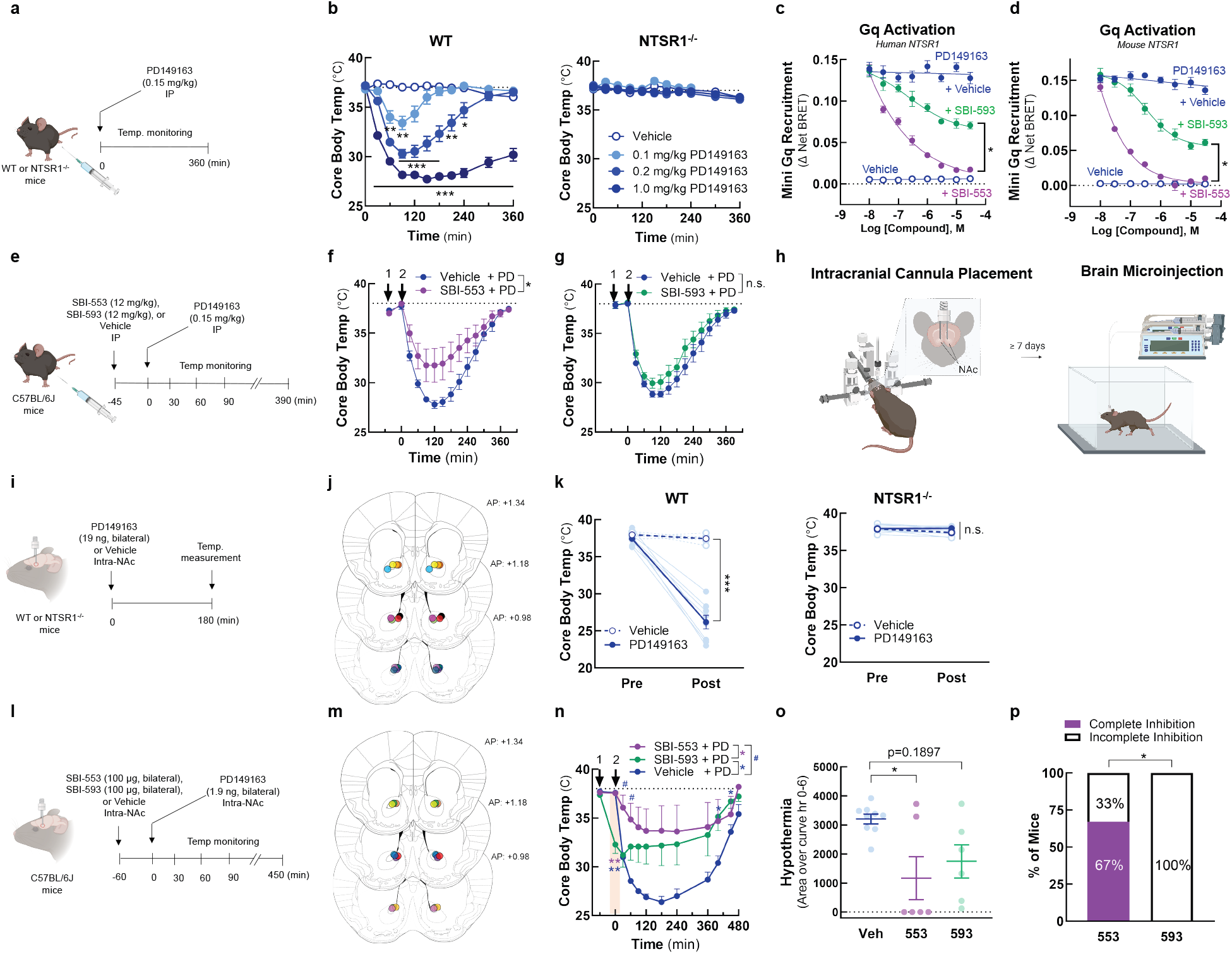
SBI-593 does not produce a complete block of NTSR1 agonist-induced, Gq-mediated hypothermia in mice. **(A-B)** *Systemic PD142948A induces an NTSR1-dependent hypothermia in mice*. **(A)** Experimental timeline. **(B)** Effect of PD149163 (0.1-1.0 mg/kg, i.p.) on core body temperature in WT and NTSR1^-/-^ mice. **(C-D)** *SBI-593 partially antagonizes PD149163-induced Gq activation by the NTSR1*. **(C)** Effect of SBI-553 and SBI-593 on PD149163-induced Gq activation by the **(C)** human and **(D)** mouse NTSR1. In HEK293T cells transiently expressing NTSR1, the NT-induced association between NTSR1 and miniGq was assessed by BRET. **(E-G)** *Systemic SBI-593 administration does not attenuate PD149163-induced hypothermia*. **(E)** Experimental timeline. **(F)** Effect of SBI-553 (12 mg/kg, i.p.) vs. Vehicle (5% HP-β-cyclodextrin) pretreatment on PD149163 (15 mg/kg,i.p.)-induced hypothermia. **(G)** Effect of SBI-593 (12 mg/kg, i.p.) vs. Vehicle (20% DMSO, 5% Tween 80) pretreatment on PD149163 (0.15 mg/kg,, i.p.)-induced hypothermia. **(H)** Overview of bilateral intracranial cannula placement in mice. Cannulas targeted the nucleus accumbens (NAc). **(I-K)** *Intra-NAc PD142948A induces an NTSR1-dependent hypothermia in mice*. **(I)** Experimental timeline. **(J)** Ink-verified cannula placements for mice in panel K. Numbers denote mm in front of bregma. AP, anterior-posterior. **(K)** Effect of microinjection of PD149163 (93 ng, bilateral) into the NAc on core body temperature in WT and NTSR1^-/-^ mice. **(L-P)** *Intra-NAc SBI-593 administration partially but does not fully attenuate PD149163-induced hypothermia*. **(L)** Experimental timeline. **(M)** Ink-verified cannula placements for mice in panel N. Numbers denote mm in front of bregma. AP, anterior-posterior. **(N)** Effect of intra-NAc SBI-553 (100 μg, bilateral) and SBI-593 (100 μg, bilateral) on PD149163 (9.3 ng, bilateral, intra-NAc)-induced hypothermia. **(O)** Area of the curve for data in panel N. Baseline set at 37°C. **(P)** Proportion of mice exhibiting complete vs. incomplete blockade of PD149163-induce hypothermia by treatment group. For information on N, statistical comparisons, and curve parameters, see **Table S6**.

Because the inability of systemic SBI-593 to block PD149163-induced hypothermia in this model could be a consequence of poor metabolic stability or brain penetrance, we evaluated the effects of these compounds following local delivery in the brain. NT produces hypothermia following microinjection into multiple brain regions^31^, including the nucleus accumbens (NAc) in mice^32^. In mice with bilateral cannula targeting the NAc core (**Fig 6H, I, J**), PD149163 microinjection reduced core body temperature via an NTSR1-dependent mechanism (**Fig 6K**). Mice received SBI-553 (100 μg, bilateral, intra-NAc), SBI-593 (100 μg, bilateral, intra-NAc), or their respective vehicles 1 hr prior to intra-NAc PD149163 treatment (**Fig 6L, M**). SBI-593 treated mice had reduced core body temperatures 1 hr after SBI-593 treatment and prior to PD149163 treatment, suggesting that SBI-593 alone produced hypothermia (**Fig 6N**). SBI-553 had no effect on core body temperature prior to PD149163 treatment and reduced PD149163 hypothermia (**Fig 6N, O**). SBI-593 reduced PD149163 hypothermia (**Fig 6N**), but the proportion of mice exhibiting a complete inhibition of hypothermia differed between the SBI-553 and SBI-593 groups, with 67% of the SBI-553-treated mice exhibiting complete inhibition compared to 0% of the SBI-593-treated mice (**Fig 6P**). This finding suggests that SBI-593 is less effective than SBI-553 at blocking NTSR1 agonist-induced, Gq/11-mediated hypothermia, indicating that BAMs with distinct G protein selectivity profiles can exhibit different *in vivo* activities.

## DISCUSSION

Although it was commonly believed that every GPCR couples to a specific Gα protein subfamily, two recent studies suggest that promiscuous G protein coupling is the norm, with 73% of the 124 GPCRs evaluated in one assessment^13^ and 71% of 100 GPCRs evaluated in another^14^ found to activate multiple G proteins across families. Compounds that shunt signaling toward some G protein pathways and away from others will enable the development of therapeutics that produce desired actions with reduced side effects and the re-targeting of established therapeutic targets for new indications. Here, we demonstrate that BAMs at the GPCR-transducer interface change G protein selectivity in specific and predictable ways, enabling rational design.

We assessed ligand-directed NTSR1 signaling using the most comprehensive transducer panel to date, a strategy made possible by the expanding toolkit for monitoring GPCR-transducer coupling in live cells in real-time^14,15,20,33^. We find that the intracellular NTSR1 BAM SBI-553 has previously unappreciated G protein activity. SBI-553 biased NT-NTSR1 not only away from Gq and toward β-arrestin recruitment, but also toward alternative G protein coupling (i.e., GoA/B, G12/13). Homology modeling indicated that SBI-553 agonized and was permissive of NT-induced activation of those G protein subtypes whose C-termini could adopt an alternative, shallow-binding conformation (e.g., GoA/B, G12/13). SBI-553 did not agonize and antagonized NT-induced activation of those G protein subtypes for which adoption of this shallow-binding formation was energetically disfavored (e.g., Gq/11). To test the hypothesis that changes to the SBI-553 scaffold could change permitted NTSR1 G protein coupling in a subtype-specific manner, we used structural models to design a panel of SBI-553 derivatives and undertook high-dimensional SAR studies. Out of this effort came molecules that differentially change the NTSR1-G protein signaling landscape. This finding demonstrates that while the surface area between the receptor and G protein is limited, this site is accessible to small molecules, and that while G protein subtype C-termini differ by only a handful of residues, subtype-specific regulation is possible.

Our screen produced both expected and unexpected results. This is due, in part, to the finding that, G protein C-termini appear capable of adopting multiple conformations. Unexpected results are anticipated, as our predictions are complicated by ligand-induced recruitment of GPCR kinases (GRKs), ligand-induced conformational changes in NTSR1, and ligand-specific changes to the conformation of associating G proteins. Extensive high-dimensional SAR with pairwise comparators, structures of NTSR1 bound to other G protein subtypes in the presence and absence of BAMs, and assessments of NTSR1-G protein dynamics will improve predictive models. Nonetheless, the distinct G protein selectivity fingerprints of SBI-553, SBI-342, and SBI-593 are proof-of-concept that one can switch GPCR G protein preference with minor chemical modifications to a single scaffold.

SBI-553 and derivatives display uniquely complex allosteric effects. Adding to this complexity, is the possibility that these ligands direct signaling differently at different timepoints post application and at different locations within the cell. GPCR G protein coupling preference is time- and location-dependent^13,34-36^, and both NT and SBI-553 stimulate NTSR1 trafficking^9^. The signaling profiles presented here capture the maximal transducer activation over a 45 min period, regardless of activation rate and subcellular location. The availability of transducers may vary by time and location, making the properties of these compounds sensitive to trafficking.

Critically, the use of intracellular modulators to drive changes in GPCR G protein selectivity is likely transferrable to other receptors, as allosteric modulators binding the intracellular domain within the seven transmembrane helical bundle have been described for five other Class A receptors (CCR2^37^, CCR9^38^, CCR7^39^, CXCR2^40^, β2AR^41^) and the Class B PTH1R receptor^42^. While these compounds have been classified as agonist or antagonists^43^, modifications to the portions of these molecules extending into the intracellular cavity may confer transducer selectivity.

Much remains unknown, including the relative expression levels and availability of transducers, how those levels change by cell-type and disease state, and the consequences of differential coupling to G proteins within the same family. Work is needed for the NTSR1, and GPCRs broadly, to determine which signaling pathways are desirable and which are undesirable in contexts of health and disease. As this information becomes available, however, we are now equipped with the tools to develop small molecule GPCR-targeting therapeutics with precise pathway selectivity.

## MATERIAL AND METHODS

### Cell lines

HEK293T/17 (Cat# CRL-11268, RRID:CVCL_1926) cells were obtained from the American Type Culture Collection (ATCC, Manassas, VA). G protein-deficient HEK293 cells (ΔGNAS, GNAL, GNAQ, GNA11, GNA12, and GNA13 HEK293 - Clone 38^44^) and β-arrestin1/2-deficient HEK293 cells (arrestin 2/3-deficient (ΔArrb1/2) HEK293 - Clone 4^24^) have been previously described. All cells were cultured in minimum essential medium (DMEM) with 10% fetal bovine serum (FBS) (Invitrogen, Cat# CX30346) and 1x antibiotic antimycotic solution (100 IU−1 penicillin, 100 μg/mL streptomycin and 250 ng/ml amphotericin B; Thermo Fisher Scientific, Cat# 15240062). Cells were grown exponentially in an incubator at 37°C under 5% CO_2_ and subcultured at ratios of 1:2-1:10 every 2 to 4 days using 0.05% trypsin-EDTA (Thermo Fisher Scientific, Cat# 25300120).

### Chemicals

All chemicals were obtained from MilliporeSigma (St. Louis, MO, USA) unless otherwise noted. SBI-0654553 HCl (abbreviated SBI-553) was synthesized by the Conrad Prebys Center for Chemical Genomics at the Sanford Burnham Prebys Medical Discovery Institute (La Jolla, CA, USA). Coelenterazine *h* and coelenterazine 400a were obtained from Cayman Chemical (Ann Arbor, MI, USA). For receptor signaling assessments, NT (Sigma, product #N6383) and PD149163 (Sigma, product #PZ0175) were maintained as 2 mM stock in 80% glycerol. SBI-553 and SR142948A (Sigma, product #SML0015) were maintained as 50 mM stocks in DMSO. SBI-0654553 analogs were synthesized using previously published methods^8^. Derivates were maintained as 50, 25 or 12.5 mM stocks in DMSO, as solubility permitted. For *in vivo* studies, SBI-553 and SBI-593 were freshly prepared from powdered stocks. When applicable, doses and concentrations were calculated based on the formula weight of the compound salts and adjusted for fractional anhydrous base weight.

### Recombinant DNA plasmids

The 3xHA-NTSR1 plasmid consists of N-terminal 3xHA-tagged wild-type human NTSR1 cloned into the pcDNA3.1(+) vector (Invitrogen, Thermo Fisher Scientific Corporation, Waltham, MA, USA) at KpnI (5’) and XbaI (3’). This construct was purchased from the University of Missouri (cDNA Bank cDNA.org, CAT# NTSR10TN00). TRUPATH was a gift from Bryan Roth (Addgene kit #1000000163). The pcDNA3.1Zeo(-) vector was acquired from Invitrogen. The human mVenus-β-arrestin1 and mVenus-β-arrestin2 plasmids consist of human β-arrestin1 (cDNA Bank cDNA.org, CAT #ARRB100002) or β-arrestin2 (cDNA Bank cDNA.org, CAT #ARRB200001) cloned in frame, through restriction cloning, into a pcDNA3.1 plasmid that contained the mVenus sequence in the N-terminal position. The Gq-Rluc8 single point mutants (corresponding Gq residues: V246^Y^, N244^G^, Y243^C^, and L238) were obtained by single nucleotide mutagenesis utilizing the QuickChange II XL (Agilent Technologies) and following the manufacturer specifications. The mVenus-tagged mini-Go was generated by G-block synthesis of the reported mini-Go sequence^45^ and in frame restriction cloning into the same backbone as the rest of the mini-G constructs. The 5 and 13 C-terminal substitutions in the TRUPATH Gα subunits were performed by oligonucleotide synthesis and annealing, followed by in frame restriction cloning. Cloning of β-arrestin and G protein sensors was performed by the University of Minnesota Viral Vector and Cloning Core and validated through Sanger sequencing.

### BRET2 G protein activation assays

In the BRET2-based TRUPATH platform^15^, G protein activation results in a decrease in BRET between an Rluc8-tagged Gα and a GFP2-tagged Gγ protein. For visualization purposes, we plotted transformed (-Δ Net BRET) data, such that G protein activation produces upward sloping curves. Curve height is a function of both the number of Gα-Gβγ complexes dissociating and the relative proximity of the Rluc8 and GFP2 tags. Because tag location differs among the Gα proteins and distinct γ subfamily members are used for each Gα protein, maximal changes in BRET are expected to differ among the Gα sensors. We retain the original BRET values on these curves to preserve information on sensor dynamic range. On day 1, HEK293T cells were plated in 6-well plates (750,000/well) in DMEM containing 10% FBS and 1% 1x antibiotic antimycotic solution. On day 2, cells were transiently transfected with NTSR1 (200 ng/well), Gβ1 or Gβ3 (100 ng/well), GFP2-tagged Gγ9, Gγ8, Gγ13, or Gγ1 (100 ng/well), and the Gα of interest using a standard calcium phosphate transfection protocol, and the pairings of Gα, Gβ, and Gγ subunits as described in Olsen et al. On day 3, cells were plated (25,000 cells/well) onto poly-D-lysine (PDK)-coated [100 ng/mL], clear bottom, white-walled 96-well plates in Opti-MEM containing 2% FBS and 1% 1x antibiotic antimycotic solution. On day 4, cells were incubated in 70 or 80 μl/well Hanks’ Balanced Salt solution (HBSS) containing calcium and magnesium and 20 mM HEPES for 5-6 hr prior to treatment. Cells receiving an SBI-553 or SR142948A pretreatment incubated in 70 μl/well of the HBSS containing 20 mM HEPES. Cells not receiving any pretreatment incubated in 80 μl/well of HBSS plus 20 mM HEPES. SR142948A was freshly prepared in HBSS from a 50 mM DMSO stock. 10x NT and PD149163 were freshly prepared in HBSS from 2 mM 80% glycerol (NTS, PD149163) stocks. 10x SBI-553 was freshly prepared in HBSS with 5% 2-Hydroxylpropyl-β-cyclodextrin (HP-β-CD, Tokyo Chemical Industry Co.) from a 50 mM DMSO stock. For structure-activity-relationship studies, 10x SBI-553 and all derivatives were freshly prepared in 30% cyclodextrin from 50, 25 or 12.5 mM DMSO stocks. A white vinyl sticker was placed on the bottom of the plate. All plates were allowed to cool for 10 minutes until they reached approximately 25°C. 10 μl of 10x SBI-553, SR142948A, or SBI-553 derivative pretreatments were added during the 10-minute cooling period, a total of 20 minutes prior to reading. 10 μl of 10x NTS, PD149163, SBI-553 or SR142948A treatments were added 10 minutes prior to reading. 10 μl/well of a 10x concentration of coelenterazine 400a (final concentration ∼7.5 μM Cayman Chemical Co., Ann Arbor, MI, USA) was added 5 minutes prior to reading. After treatment with coelenterazine 400a, plates were protected from light. Plates were read with a Tecan SparkCyto Microplate reader (Switzerland) at room temperature in ambient air every 5 minutes for 20 minutes. BRET2 ratios were computed as the ratio of GFP2 emission to RLuc8 emission. The Δ net BRET ratio was calculated by subtracting the stimulated GFP2/Rluc8 ratios from control GFP2/Rluc8 ratios for each read. The minimum Δ net BRET ratio over time was averaged within treatments and combined between experiments. Data are presented as negative mean Δ net BRET ratio ± SEM from at least three independent experiments.

### BRET1 β-arrestin recruitment assays

Recruitment of Venus-tagged β-arrestin2 to Renilla luciferase (Rluc8)-tagged NTSR1 was assessed in HEK293T cells using a bioluminescence resonance energy transfer 1 (BRET) assay, as described^9^. Previous studies evaluating NTSR1 β-arrestin agonism have used bovine or rodent β-arrestin constructs^9,10^. To assess recruitment of the human β-arrestins: On day 1, HEK293T cells were plated in 6-well plates (750,000/well) in growth media. On day 2, cells were transiently transfected with Rluc8-tagged NTSR1 (100 ng/well), Venus-tagged β-arrestin2 or Venus-tagged β-arrestin1 (1.4 μg/well), and pcDNA3.1 (1.5 μg/well) using a standard calcium phosphate transfection protocol. On day 3, cells were plated onto PDK-coated [100 ng/mL], clear bottom, white-walled 96-well plates (40,000 cells/well) in Opti-MEM containing 2% FBS and 1x antibiotic antimycotic solution. On day 4, cells were incubated in 70 or 80 μl/well HBSS containing calcium and magnesium and 20 mM HEPES for 5-6 hr prior to treatment. Cells receiving an SBI-553 or SR142948A pretreatment incubated in 70 μl/well of the HBSS containing 20 mM HEPES. Cells not receiving a pretreatment were incubated in 80 μl/well of HBSS plus 20 mM HEPES. 10x NT and PD149163 were freshly prepared in HBSS from 2 mM 80% glycerol (NT, PD149163) stocks. 10x SBI-553 was freshly prepared in HBSS with 5% HP-β-CD from a 50 mM DMSO stock. For structure-activity-relationship studies, 10x SBI-553 and all derivatives were freshly prepared in 30% HP-β-CD from 50, 25 or 12.5 mM DMSO stocks. For compound combinations studies, cells were pretreated with 10 μl of 10x SBI-553, SR142948A, or vehicle pretreatments 15 min prior to reading. 10 μl/well of a 10x concentration of coelenterazine *h* (final concentration 4.7 μM; Cayman Chemical Co., Ann Arbor, MI, USA) was added 10 minutes prior to reading, followed by treatment with 10 μl of 10x NT 5 minutes prior to reading. Cells receiving no pretreatment received 10 μl of 10x NT, PD149163, SBI-553, and SR142948A treatments 10 minutes prior to reading and 10X coelenterazine h 5 minutes prior to reading. A white vinyl sticker was placed on the bottom of each plate. Plates were read with a CLARIOstar Plus microplate reader (BMG Labtech, Ortenberg, Germany) set at 35°C 5-, 10-, 15-, and 30-min post-treatment. BRET1 ratios were computed as the ratio of Venus emission to RLuc8 emission. The Δ net BRET ratio was calculated by subtracting the stimulated Venus/Rluc8 ratios from control Venus/Rluc8 ratios for each read. The maximum Δ net BRET ratio over time was averaged within treatments and combined between experiments. Data are presented as mean Δ net BRET ratio ± SEM from at least three independent experiments.

### TGFα shedding G protein activation assays

TGFα shedding assays of G protein activation were performed as originally presented^20^, with previously described modifications^9,46^. These modifications included using G protein-deficient (ΔGNAS, GNAL, GNAQ, GNA11, GNA12, and GNA13) HEK293 cells^44^ and the fluorescent substrate 4-methylumbelliferyl phosphate (4-MUP), at a working concentration of 1 mM per well, instead of p-Nitrophenyl phosphate (p-NPP). On day 1, G protein-deficient HEK293 cells were plated in 6-well plates (750,000/well) in growth media. On day 2, expression vectors were transiently transfected in HEK293 cells using Lipofectamine 2000 (Invitrogen; 8 μL per well in a 6-well plate). Expression vectors included 1.875 μg AP-TGFα, 750 ng HA-tagged NTSR1, and 350 ng of Gα protein. On day 3, transfected cells were detached with a brief rinse (1 mL, ∼30 s) of PBS followed by 0.5 mL per well of 0.05% trypsin-EDTA (Gibco). The cell suspension was pelleted by centrifugation (200g, 5 min) followed by a resuspension in 3 mL HBSS containing 5 mM HEPES (pH 7.4) and a 10-minute incubation at room temperature. Cells were again centrifuged (200g, 5 min) and resuspended in 4 mL HBSS containing 5 mM HEPES (pH 7.4). The resuspended cells were plated in 80 μL per well in a 96-well plate and placed in an incubator at 37°C with 5% CO_2_. After a 30-min incubation, cells were treated with 10 μL of vehicle (HBSS), 10X final concentrations of SR142948A, or SBI compounds and incubated at 37°C for 20 min. For preparation of 10x stocks, SBI compounds and SR142938A were diluted directly from 1-10 mM DMSO stocks into HBSS. The final concentration of DMSO did not exceed 0.4%. After a 20-min incubation with SBI, cells were treated with 10 μL of 10X concentration of NT and incubated at 37°C for 1 hour. For inhibition studies, 10 nM NT was used for Gq, Gi1/2, and G12 constructs, and 100 nM was used for the Go construct, concentrations selected to elicit maximal NT-induced shedding. Plates were centrifuged (190 g, 2 min) and conditioned media (80 μL) was transferred into a new 96-well plate. After a 20-min incubation at room temperature, 2 mM 4-MUP-containing solution was added (80 μL per well) to both the conditioned media and the cell plate. Alkaline phosphatase activity was measured using a CLARIOstar Plus microplate reader (Germany) set to 25°C. Data were collected before and after a 1-hour incubation at 37°C. Excitation was set at 360 nm (±10 nm) and emmision at 450 nm (±15 nm). TGFα shedding activity was calculated by dividing the amount of phosphatase activity present in the conditioned media by the amount present on the cells plus the conditioned media. All values were standardized to background shedding activity.

### Mini-G Protein recruitment assays

Mini G proteins have a truncated N-terminus and contain a mutation that uncouples GPCR binding from nucleotide release, allowing them to form more stable associations with GPCRs than unmodified G proteins^22,45^; these associations are amenable to monitoring by BRET. The recruitment of Venus-tagged mini-G proteins^22^ to Rluc8-tagged NTSR1 was assessed by BRET. Note that while mini-Go and G12 are derived from their full-length versions, the mini Gq and mini Gi1 constructs were derived from the Gs backbone with substitution of the α5 helix^22,45^. To assess recruitment, on day 1, HEK293T cells were plated in 6-well plates (750,000/well) in growth media. On day 2, cells were transiently transfected with Rluc8-tagged NTSR1 (100 ng/well), a venus-tagged mini-G protein, and pcDNA3.1 (1.5 μg/well or 2.65 μg/well) using a standard calcium phosphate transfection protocol. Cells transfected with a venus-tagged mini Gq or mini Gi1 received 250 ng/well. Cells transfected with a venus-tagged mini G12, mini Gs, or mini-Go received 1.5 μg/well. On day 3, cells were plated onto PDK-coated [100 ng/mL], clear bottom, white-walled 96-well plates (40,000 cells/well) in Opti-MEM containing 2% FBS and 1x antibiotic antimycotic solution. On day 4, cells were incubated in 70 or 80 μL/well HBSS containing calcium and magnesium and 20 mM HEPES for 3-4 hr prior to treatment. Cells receiving an SBI-553, PD149163, or SR142948A pretreatment incubated in 70 μl/well of the HBSS containing 20 mM HEPES. Cells not receiving any pretreatment incubated in 80 μl/well of HBSS plus 20 mM HEPES. Cells were pretreated with treated with 10 μL of 10X concentration of SBI compounds, SR142948A, PD149163, or vehicle (5% or 30% HP-β-cyclodextrin) 20 minutes prior to reading. Cells were then treated with vehicle (HBSS) or 100 nM NT 10 min prior to reading. Finally, cells were treated with 10 μl/well of a 10x concentration of coelenterazine h (final concentration 4.7 μM) 5 min prior to reading. Plates were read on a CLARIOstar Plus microplate reader set at 25°C 5-, 10-, 15-, and 30-min post-treatment. Mini G protein recruitment was calculated by subtracting the stimulated Venus/Rluc8 ratios from control Venus/Rluc8 ratios for each read (Δ net BRET).

### Structural models. Optimization of the NT-NTSR1-SBI-553-GoA complex structure for modeling

The CryoEM structure of NTRS1 bound to SBI-553 and Go (PDB: 8NF0) was used at the starting point for modeling. Structure preparation was performed with Maestro (Schrödinger). VDW clashes were removed by constrained minimization using the OPLS4 force field. A full conformational analysis was performed with the conformational ensemble generated by MOE (Chemical Computing Group) and minimization using density functional theory calculations (Gaussian, ωb97xd/6-311+G** functional). The SBI-553 piperidine preferred a chair conformation over the twist boat by >3.1 kcal/mol. As a result, the SBI-553 conformation from PDB:8JPB was substituted into the model by rigid superposition of the two SBI-553 conformations followed by a second constrained minimization of the minor clashes resulting from this replacement. Molecular mechanics calculations (minimization and molecular dynamics) with multiple force fields orient the fluorine on SBI-553’s cyclopropyl group perpendicular to the quinazoline ring. To investigate this orientation, a rotational analysis was performed on SBI-553, SBI-342, and SBI-593 at 10° increments about the quinazoline-cyclopropane C-C bond with full minimization using ωb97xd/6-311+G**. For each molecule, there was a strong preference (2.5 to 3.2 kcal/mol) for the conformation reported in Krumm et al., in which the substituent on the cyclopropane (F for SBI-553, CH_3_ for SBI-342, and H for SBI-593) is in the plane of the quinazoline and pointed away from the appended piperidine. Subsequent protein minimizations and molecular dynamics simulations employed explicit constrains to maintain desired small molecule conformations.

### G protein homology modeling

Homology models of each G protein were built with MOE (v2022.02, Chemical Computing Group, Montreal, CA) with the conformation of each mutated side chain optimized individually using the Amber 10:EHT force field built into MOE. Models of Gq conformations required building the H1/H2 loop absent in 8FN0. This loop is resolved in the 6OS9 structure, so the missing residues were concatenated onto 8FN0 using the Homology Model Application in MOE. Minimization of the protein complexes containing SBI-342 and SBI-593 utilized the OPLS4 force field in Maestro (v14.1.138, Schrödinger, Inc., NY, NY).

### Molecular dynamics simulations

Molecular dynamics simulations with Gq and SBI-593 were conducted with Desmond using the Schrödinger suite (v13.7.125). Except for the aforementioned small molecule conformational constraint, default conditions were used including explicit water, an explicit membrane, and 0.15 M NaCl. Four discrete simulations were performed, each from a slightly different starting orientation of Gq with the time ranging between 200 ps and 300 ps. Once the complex stabilize to a consistent RMSD, 20-25 frames from each run were selected from periods of greatest RMSD stability. Each complex was fully minimized in Maestro, where conformations, orientations, and total energies were compared.

### Free energy calculations

The free energy required to dissociate the individual NT-NTSR1-SBI-553-’open’ position G protein complexes created by homology modeling was estimated using ‘Protein interfaces, surfaces and assemblies’ service PISA at the European Bioinformatics Institute (http://www.ebi.ac.uk/pdbe/prot_int/pistart.html)^47^. Individual NT-NTSR1-SBI-553-G protein complex structures were uploaded iteratively. ΔG^diss^ values in kcal/mol were recorded and reported as fold-change over ΔG^diss^ of the NT-NTSR1-SBI-553-GoA complex.

### Animals

All mouse studies were conducted in accordance with the National Institutes of Health Guidelines for Animal Care and Use of Laboratory Animals and with approved animal protocols from the University of Minnesota University Animal Care and Use Committee. The mice studied in the embodied work include: C57BL/6J mice (The Jackson Laboratory, Bar Harbor, ME, Cat# 000664), global NTSR1^-/-^ mice (B6.129P2-Ntsr1tm1Dgen/J, Deltagen, The Jackson Laboratory Strain #:005826), and their respective WT littermates. All mouse lines were backcrossed onto a C57BL/6J genetic background for ≥ 10 generations prior to use. At study initiation, all animals were adults. Mice were 8-20 weeks old, weighed 19–30 g, and were age-matched across experimental groups. Experiments included both male and female mice, and experimental groups were sex-matched. NTSR1^-/-^ mice for experimental use were exclusively generated by NTSR1^+/-^ X NSTR^+/-^ breeding, such that littermate WT mice could serve as controls. In NTSR1^-/-^ studies, littermates of the same sex and genotype were randomly assigned to experimental groups. Mice for systemic PD149163-induced hypothermia studies were group housed in conventional cages with Teklad irradiated corncob bedding (Inotiv) and Enviro-Dri nesting material (Fibercore, Cleveland, OH) and maintained on a 14:10-hour light/dark cycle. Mice for local NAc injection studies were singly housed following implantation of intracranial cannulas in conventional cages with Teklad irradiated corncob bedding (Inotiv) and Enviro-Dri nesting material (Fibercore, Cleveland, OH), and maintained on a 14:10-hour light/dark cycle. Experiments began at the beginning of the light cycle (i.e., within 3 hrs of light cycling beginning). Tap water and standard laboratory chow were supplied ad libitum, except during testing.

### Assessments of core body temperature

Core body temperatures were measured using a rectal probe thermometer for mice (Thermalert Model TH-8, Physitemp Instruments, Inc., Clifton, NJ), as described ^9^. Core body temperatures were recorded from age-matched male and female NTSR1^-/-^ and WT mice prior to treatment (Time 0) and 30-, 60-, 90-, 120-, and 300-min post-treatment. Mice were gently restrained during the procedure and acclimated to this process during baselining. For single compound dosing studies, baseline temperatures were recorded, and animals received either PD149163 (0.03-1 mg/kg) or PD149163’s vehicle (physiological saline). All treatments were administered i.p. in a volume of 10 ml/kg. For systemic multiple compound dosing studies, animals received SBI-553 (12 mg/kg, i.p.) or its vehicle (5% cyclodextrin) or SBI-593 (12 mg/kg, i.p.) or its vehicle (20% DMSO, 5% Tween 80) prior to the start of the study (−45 min). Following temperature recording at Time 0, animals received PD149163 (0.15 mg/kg) in volumes of 10 μL/kg i.p.

### Intracranial cannulation and local NAc injections

Local NAc microinjections were accomplished following bilateral guide cannula placement in NTSR1^-/-^ mice, their WT littermates, C57BL/6J mice. Cannulas were acquired from Plastics One [bilateral guide: 2.0 mm spacing, 26G, 4 mm below pedestal; bilateral internal: 2.0 mm spacing, 33G, 0.5 mm projection; bilateral dummy: 2.0 mm spacing, 0.008”/0.2 mm, 0 mm projection]. Bilateral guide cannulas were inserted into the NAc at +1.3 mm AP with 2.0-mm spacing (±1.0 mm ML) and −4.5 mm DV and fixed to the skull with dental cement. Mice were single-housed post-surgery and allowed to recover for at least 7 days. After recovery, compounds were injected bilaterally using an automated syringe pump (Harvard Apparatus). SBI-553 (molecular weight 450 g/mol) was dissolved in 1% (vol/vol) DMSO and 20% (vol/vol) HP-β-cyclodextrin at 74.07 mM. 100 μg SBI-553 in 3 μL was injected per side at a rate of 0.2 μL/min. SBI-593 (molecular weight 458 g/mol) was dissolved in 80% (vol/vol) N,N-dimethylacetamide, 10% (vol/vol) Tween 80, and 10% (vol/vol) UltraPure distilled water at 72.7 mM. 100 μg SBI-593 in 3 μL was injected per side at a rate of 0.2 μL/min. After local injection of SBI compounds, mice were placed in their home cages for 1 hour prior to intra-NAc PD149163 administration. PD149163 (molecular weight 943.91 g/mol) was diluted from a stock concentration of 2 mM in 80% glycerol to 10 or 100 μM in separate SBI compound or vehicle preparations. 18.9 or 1.89 ng PD149163 in 0.2 μL was injected per side at a rate of 0.2 uL/min. The PD149163 dose used in the WT vs. NTSR1^-/-^ was 18.9 ng. The PD149163 dose used in the SBI pretreatment experiments was 1.89 ng. After local injection of PD149163, body temperature was monitored every 30-60 min for 8 hours. For all experiments mice were randomly assigned to treatment groups for no more than two experiments, separated by a minimum of seven days. Cannula placements were verified via ink injection. Following the experiment, 0.5 μL India ink (1:20 (vol/vol) saline) was injected at a flow rate of 0.5 μL/min to visualize the observed injection site. Mice were anesthetized with isoflurane and brains were harvested 1 hour after ink microinjection. Brains were sliced in 250 μm sections using a vibratome (VT1000S, Leica, Deer Park, IL).

### Statistical analysis

All data are represented as mean ± SEM, unless otherwise indicated. Data were analyzed and plotted using the software GraphPad Prism version 10.1.2. Information on curve fitting, statistical tests, and N are provided in supporting information Tables 1-6. In supporting information Tables 1-6, N represents the number of biological replicates. All data represent the average of at least 3 biological replicates. Individual cell-based experiments included 2 or 3 technical replicates for every condition. A p-value of < 0.05 was accepted as statistically significant.

### Figure illustration

Method and concept figure illustrations were created using BioRender (Toronto, ON). Images of NTSR1 and G protein structures were created in ChimeraX (UCSF, v1.6).

## Abbreviations

NTSR1: neurotensin receptor 1
GPCR: G protein-coupled receptor
BAM: biased allosteric modulator
DRC: dose-response curve
SAR: structure-activity relationship

## SUPPLEMENTAL INFORMATION

Attached as a separate file.

## DATA AVAILABILITY

The data that support the findings of this study are available from the corresponding author upon request.

## ACKNOWLEDGEMENTS

TRUPATH was a gift from Bryan Roth (Addgene kit #1000000163). We thank Orion Rainwater and Shannah Serres for the maintenance of the NTSR1^-/-^ and C57BL/6J mouse colonies and Nancy Xiong and Lavina Iskander for their technical assistance. This work was supported, in part, by NIH/NIDA R00 DA048970 (L.M.S.), NIH/NIDA UH3 DA050316 (M.R.J., L.S.B., L.M.S., S.H.O.), NIH/NIDA R01 DA061773 (L.M.S., S.H.O.), DOD W81XWH-22-1-0266-PR211151 (L.S.B., L.M.S.), and the University of Minnesota Foundation AP-0323-04 (L.M.S.). This work was also supported by the NIH/NIDA P30DA048742-sponsored Viral Innovation Core and Viral Vector and Cloning Core at the University of Minnesota (E.M.). A.I. was funded by KAKENHI JP21H05113 and JP24K21281 from the Japan Society for the Promotion of Science (JSPS), JP22ama121038 and JP22zf0127007 from the Japan Agency for Medical Research and Development (AMED), and JPMJFR215T and JPMJMS2023 from the Japan Science and Technology Agency (JST).

## AUTHORS CONTRIBUTIONS

Study design: LMS, SHO, MNM, KLP, AA, CK. Obtained funding: LMS, LSB, MRJ, SHO. Data collection: AA, MNM, KLP, CK, CR, NF. Data analysis: MNM, KLP, AA, SHO, LMS. Data interpretation: MNM, KLP, AA, CR, AI, LSB, EM, SHO, LMS. Drafted paper: LMS, SHO, MNM, KLP, AA. Revised paper critically for intellectual content: SHO, MNM, KLP, EM, AA, CK, NF, CR, AI, LSB, MRJ, LMS. Moore & Person et al. GPCR ALLOSTERIC MODULATORS CHANGE G PROTEIN SUBTYPE SELECTIVITY

