## Supplementary Material for "Design of allosteric modulators that change GPCR G protein subtype selectivity"

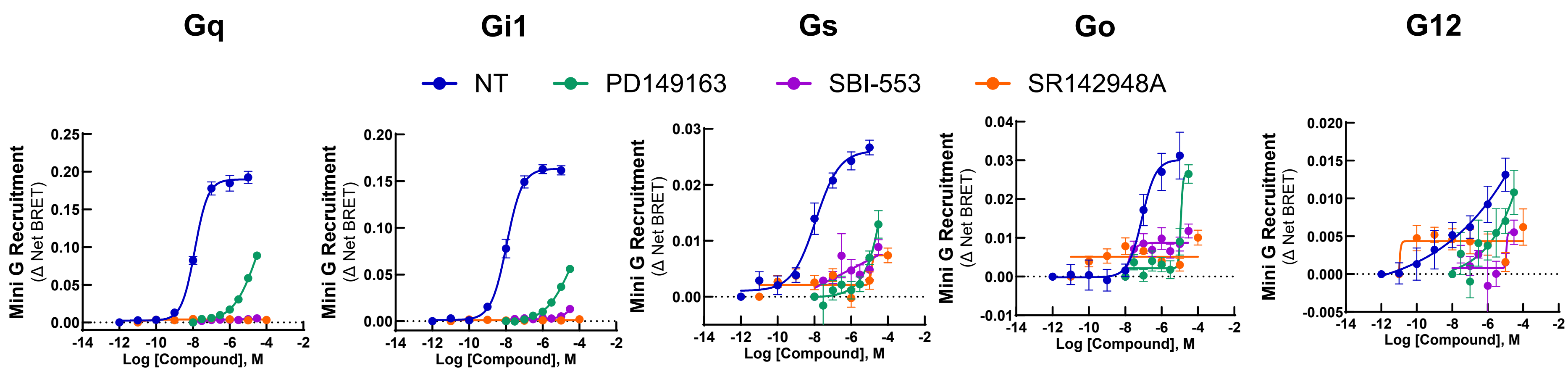

**Supplemental Figure 1. Ligand-induced mini G protein recruitment to the NTSR1.** In HEK293T cells transiently expressing NTSR1 and mini Gq, Gi1, Gs, Go, or G12, activation was assessed following treatment with the endogenous agonist NT, the modified NT peptide analogue PD149163, the β-arrestin-biased ligand SBI-553, and the orthosteric antagonist SR142948A. N=3.

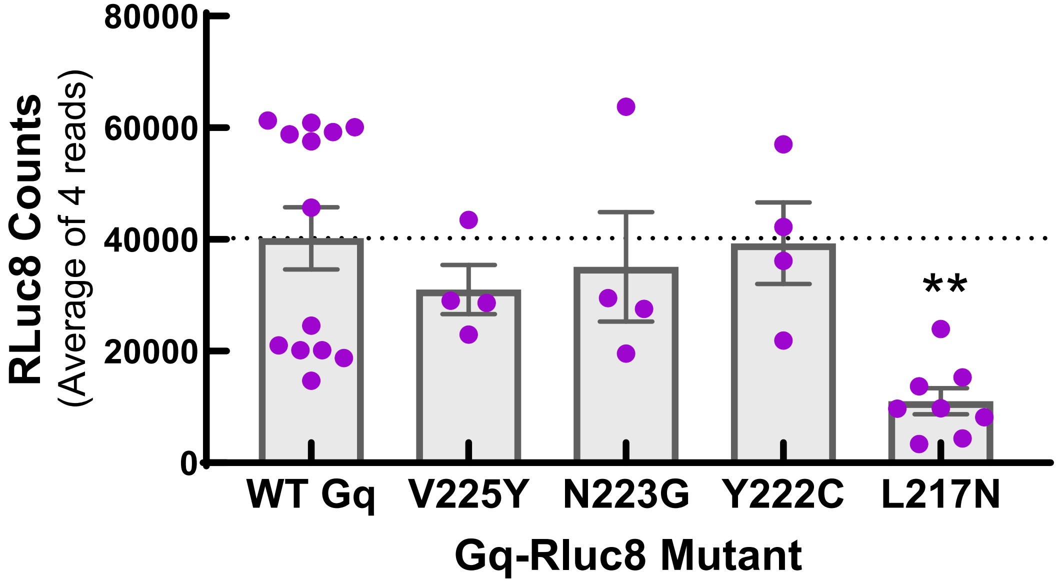

**Supplemental Figure 2. Relative expression of Gq-Rluc8 mutants as determined by total Rluc8 counts.** Rluc8 emission from experiments presented in Figure 4 M. Rluc8 counts per plate were averaged over 4 reads. One-way ANOVA followed by Tukey's multiple comparisons test. $F(4, 28) = 4.533$, $p=0.0060$. **$p<0.001$ L217N vs. WT Gq. N = 4-13.

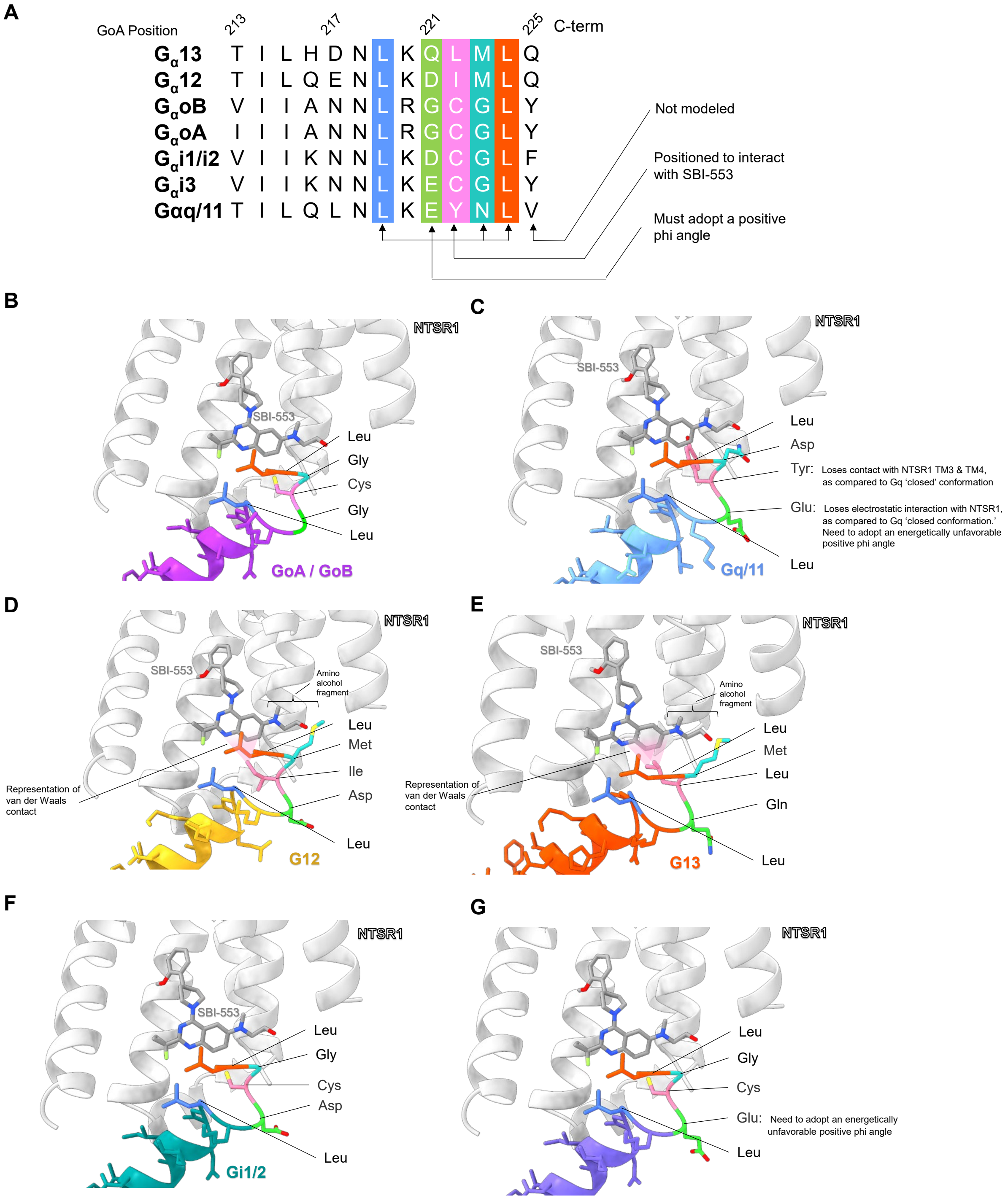

**Supplemental Figure 3. NTSR1:SBI-553:Gα homology models provide a rational basis for SBI-553's observed G protein/β-arrestin selectivity. (A)** G protein C-terminal alignments with residues positioned to interact with SBI-553 and determine stability of the NTSR1 interaction colored. Residues in proximity to SBI-553 (i.e., 219, 222, 223, 224), residues that adopt alternative positions relative to the 'closed' conformation, and residues that interact with NTRS1 are all predicted to have an impact on G protein selectivity. Other residues may also play a role but are difficult to predict with our current models. **(B)** We propose that GoB behaves similarly to GoA because their C-termini are identical until V213, which is not positioned to interact with either NTSR1 nor SBI-553 in the NTSR1:SBI-553:GoA open position model. **(C)** Three structural differences may increase the energy of this conformation for Gq, relative to GoA and relative to the 'closed' NTSR1:Gq conformation: (1) E242 of Gq (G221 in GoA) makes a dipolar interaction with the backbone NH of Q98 of NTRS1 in the 'closed' conformation that would be absent in the 'open' position, (2) forcing E242 to adopt a positive phi angle, while allowed, is energetically less favorable compared to G221, and (3) in the 'closed' conformation,

Y243 of Gq (C222 in GoA) extends between TM3 and TM4 and has a large contact energy, which would be absent in the ‘open’ conformation. Within their 13 C-terminal amino acids, **(D)** G12 and **(E)** G13 have 9 polymorphic differences with GoA. Of these, only 223 and 222 are positioned to interact with SBI-553. The final C-terminal residue at position 225 differs between G12/13 and GoA but is unresolved in our reference GoA structure PDB 8FN0 and difficult to incorporate in the model. Notably, PDB 8FMZ (Gq) indicates that it turns away from the SBI-553 binding pocket. The six remaining amino acid changes in G12 and G13 point away from NTRS1 in the open conformation. For both G12 and G13, M223 must adopt a slightly higher energy conformation with gauche interactions, but these are compensated by close contact with the SBI-553 amino alcohol fragment. G12’s I222 and G13’s L222 both make productive van der Waals contact with SBI-553. The G12 and G13 models predict that this complex if energically favorable, consistent with the observed activation of these G proteins by SBI-553. **(F)** Gi1 differs from GoA at four residues, none of which, however, are expected to make contact with SBI-553. **(G)** Gi3 has a sequence intermediate to GoA and Gq. Unlike Gq, Gi3 lacks the tyrosine at 222. As a result, relative to Gq, the models predict that Gi3 will more easily adopt the open conformation.

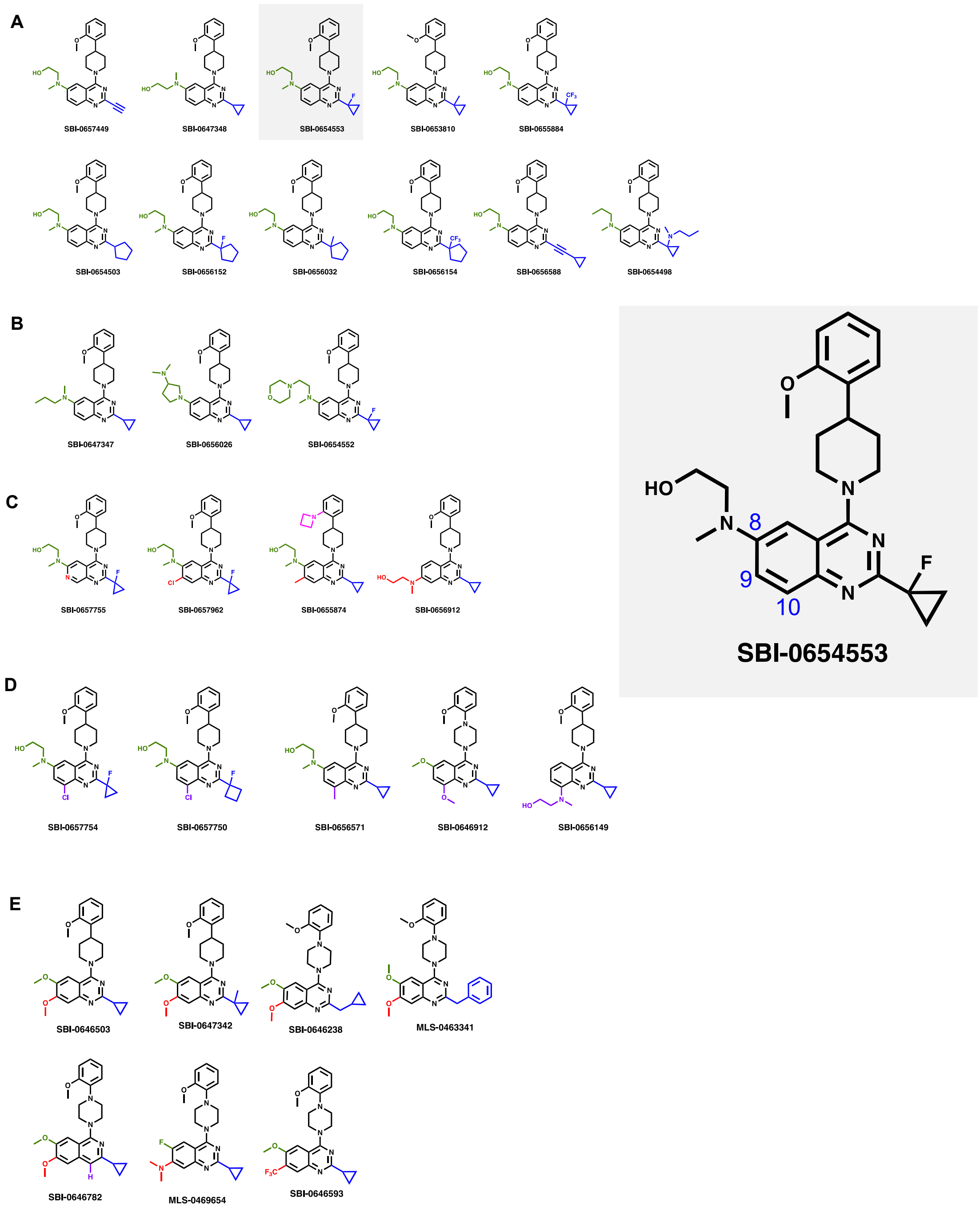

**Supplemental Figure 4. Structures of SBI-553 analogues screened. (A)** Changes to the cyclopropane (blue). Sorted left to right from small to large**. (B)** Changes to the quinazoline 8-position (green). Sorted left to right from small to large. Groups also have different rigidity / basicity / polarity. Note SBI-348 is the best comparator for compounds ‘347 and ‘026 and SBI-553 is the best comparator for compound ‘552. **(C)** Changes to the quinazoline 9-position (red). Sorted left to right from small to large. **(D)** Changes to the quinazoline 10-position (purple). Sorted left to right from small to large. **(E)** Dimethoxy (early hit) compounds and related compounds.

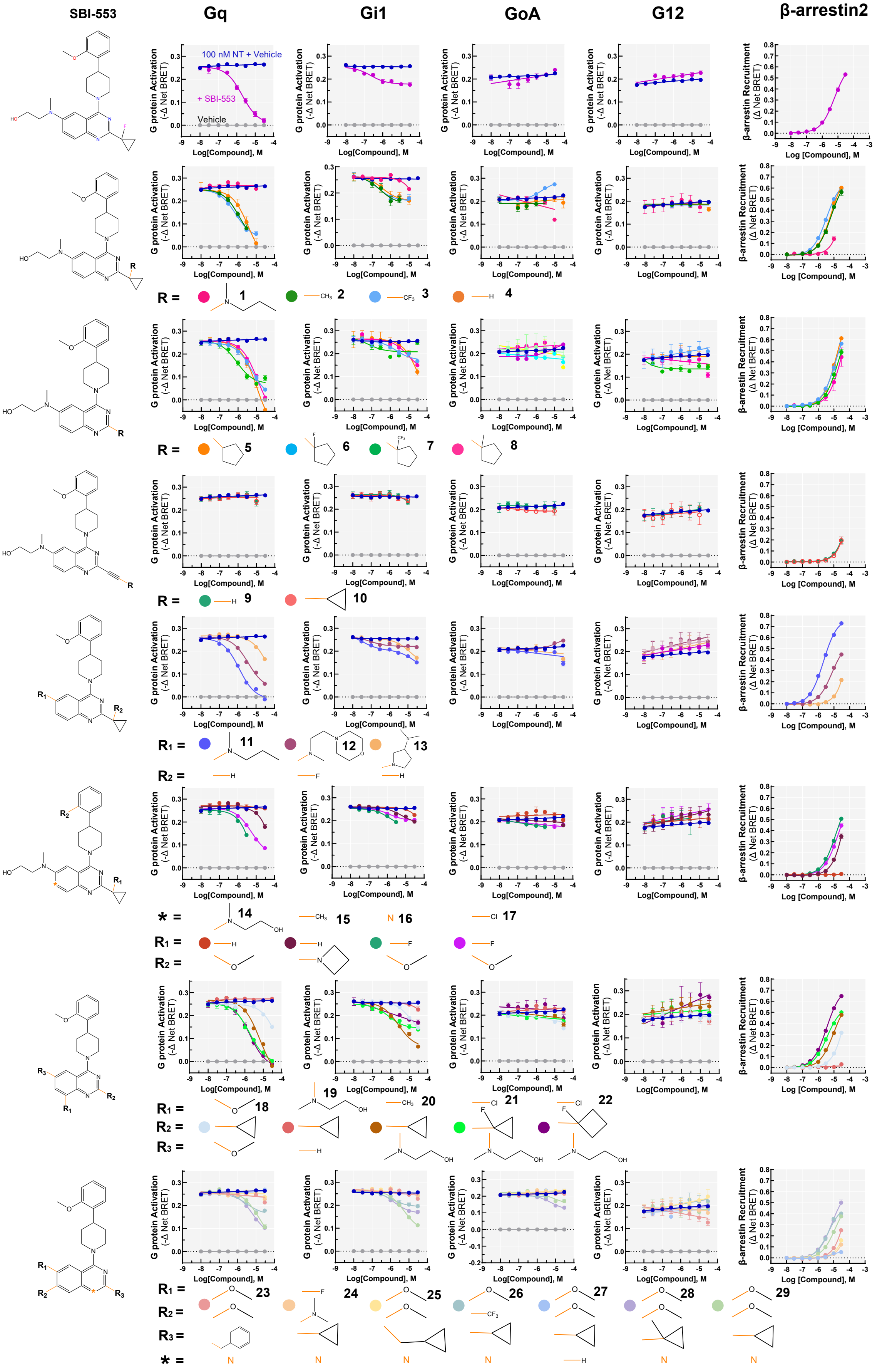

| # | SBP ID |
|---|---|
| 1 | SBI-0654498 |
| 2 | SBI-0653810 |
| 3 | SBI-0655884 |
| 4 | SBI-0647348 |
| 5 | SBI-0654503 |
| 6 | SBI-0656152 |
| 7 | SBI-0656154 |
| 8 | SBI-0656032 |
| 9 | SBI-0657449 |
| 10 | SBI-0656588 |
| 11 | SBI-0647347 |
| 12 | SBI-0654552 |
| 13 | SBI-0656026 |
| 14 | SBI-0656912 |
| 15 | SBI-0655874 |
| 16 | SBI-0657755 |
| 17 | SBI-0657962 |
| 18 | SBI-0646912 |
| 19 | SBI-0656149 |
| 20 | SBI-0656571 |
| 21 | SBI-0657754 |
| 22 | SBI-0657750 |
| 23 | MLS-0463341 |
| 24 | MLS-0469654 |
| 25 | SBI-0646238 |
| 26 | SBI-0646593 |
| 27 | SBI-0646782 |
| 28 | SBI-0647342 |
| 29 | SBI-0646503 |

**Supplemental Figure 5. Screen of SBI-553 derivatives in assessments of Gq, Gi1, Go, and G12 antagonism and β-arrestin agonism.** In HEK293T cells transiently expressing NTSR1, SBI-553, and 29 selected analogues (300 nM - 30 μM) were assessed for their ability to antagonize 100 nM NT-induced G protein activation by TRUPATH and stimulate β-arrestin2 recruitment independently by BRET. A table associating SBP compound ID numbers with screen IDs (1-29) is provided at right. Structures on the right of each row reflect the structural base for all compounds in that row. Some compounds selected for follow-up analyses produced non-NTSR1 mediated effects on G protein sensor activity. Concentrations producing these effects have been removed. Vehicle, 3% β-HB-cyclodextrin, <0.25% DMSO in HBSS. N = 3.

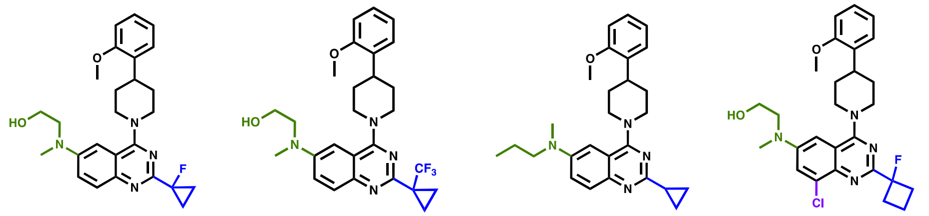

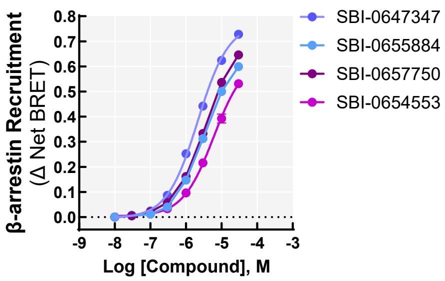

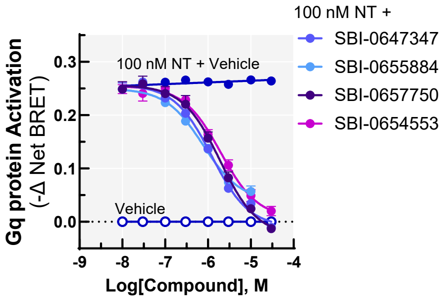

**D**

| SBP ID | Screen ID | β-arrestin2 $EC_{50}$ (μM) | β-arrestin2 $E_{max}$ (Δ BRET) | Gq $IC_{50}$ (μM) | Gq Bottom (Δ BRET) |
|---|---|---|---|---|---|
| SBI-0654553 | - | 6.0 (5.0-7.2) | 0.63 (0.60-0.68) | 1.9 (1.4-2.8) | 0.01 (-0.01-0.02) |
| SBI-0655884 | 3 | 3.4 (3.0-3.7)* | 0.67 (0.65-0.69) | 0.76 (0.50-1.2)* | 0.04 (0.02-0.06) |
| SBI-0647347 | 11 | 2.2 (2.0-2.4)* | 0.77 (0.78-0.79)* | 3.7 (2.5-5.5) | 0.03 (0.01-0.06) |
| SBI-0657750 | 22 | 2.2 (1.9-2.4)* | 0.77 (0.75-0.79)* | 1.9 (1.3-2.8) | -0.02 (-0.05-0.00) |

Data presented as mean (95% confidence interval). Asterisks (*) indicate confidence intervals that do not overlap with that of SBI-553.

**Supplemental Figure 6. SBI-553 analogues with increased β-arrestin2 agonist potency. (A)** Structures of SBI-553 and analogues with increased β-arrestin2 agonist potency. **(B)** Concentration-response curves for SBI-553 and selected analogues in a BRET-based assay of human β-arrestin2 recruitment to the human NTSR1. In HEK293T cells transiently expressing NTSR1, SBI-553, and analogues (300 nM - 30 μM) were assessed for their ability to stimulate β-arrestin2 recruitment independently by BRET. **(C)** Concentration-response curves for SBI-553 and selected analogues in a BRET-based assay of antagonism of NT-induced Gq activation. In HEK293T cells transiently expressing NTSR1, SBI-553, and analogues (300 nM - 30 μM) were assessed for their ability to NT-induced Gq activation by TRUPATH. **(D)** Table summarizing SBI compound agonist potency and efficacy at β-arrestin2 and antagonist potency and efficacy at Gq. Data are presented as mean (95% confidence interval). Asterisks (*) indicate confidence intervals that do not overlap with that of SBI-553. Note that changes in potency in β-arrestin do not always translate to changes in potency in Gq antagonism, suggesting that these properties can vary independently. SBI compounds were prepared in 3% β-HB-cyclodextrin, <0.25% DMSO in HBSS. NT vehicle, HBSS. N=3.

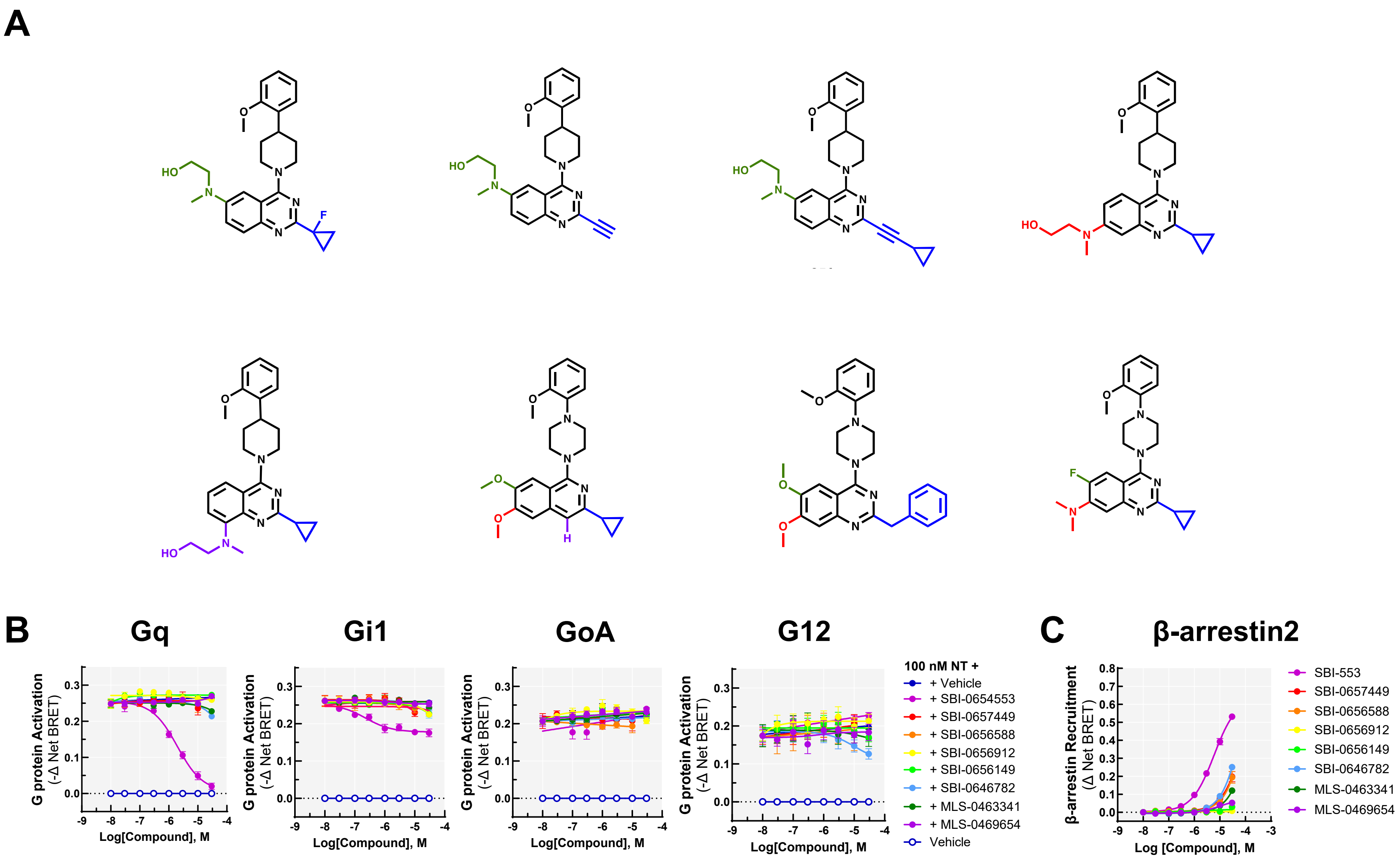

**Supplemental Figure 7. Compounds with little to no activity in functional assays.** These compounds may be silent allosteric ligands or may not bind NTSR1. **(A)** Structures of SBI-553 and analogues without activity in the evaluated assays. **(B)** Concentration-response curves for SBI-553 and analogues that lack activity in the evaluated concentration range in assessments of antagonism of NT-induced Gq activation by TRUPATH and β-arrestin agonism by BRET. In HEK293T cells transiently expressing NTSR1, SBI-553, and 29 selected analogues (300 nM - 30 μM) were assessed for their ability to antagonize 100 nM NT-induced Gq protein activation and **(C)** to promote agonism of β-arrestin2 independently. SBI compounds were prepared in 3% β-HB-cyclodextrin, <0.25% DMSO in HBSS. N = 3.

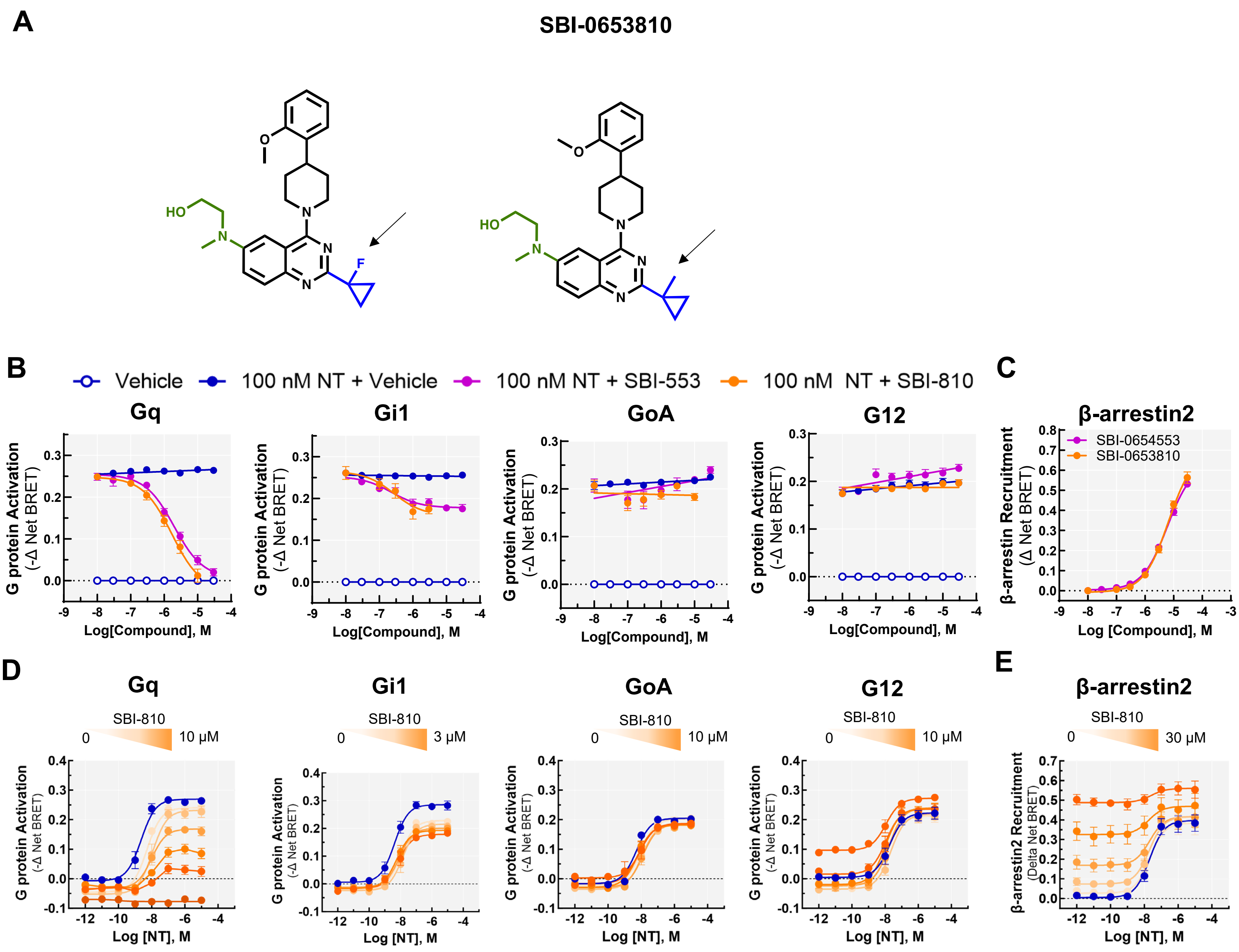

A
SBI-0653810
B
Vehicle
100 nM NT + Vehicle
100 nM NT + SBI-553
100 nM NT + SBI-810
Gq
Gi1
GoA
G12
G protein Activation (-Δ Net BRET)
Log[Compound], M
C
β-arrestin2
SBI-0654553
SBI-0653810
β-arrestin Recruitment (Δ Net BRET)
Log [Compound], M
D
Gq
Gi1
GoA
G12
SBI-810
0
10 μM
3 μM
G protein Activation (-Δ Net BRET)
Log [NT], M
E
β-arrestin2
SBI-810
0
30 μM
β-arrestin2 Recruitment (Delta Net BRET)
Log [NT], M

**Supplemental Figure 8. Substitution of SBI-553's fluorine for a methyl group does not reduce NT-induced Go activation. (A)** Structural comparison of SBI-553, SBI-810, and SBI-342. Arrows indicate the methyl substituent that differs between SBI-553 and both SBI-810 and SBI-342. **(B)** In HEK293T cells transiently expressing NTSR1, SBI-810 and SBI-553 were assessed for their ability to antagonize NT-induced G protein activation using the bioluminescence resonance energy transfer 2-based TRUPATH assay. SBI-810 produced non-NTSR1-dependent changes in some G protein sensors at high (i.e., 10 or 30 μM) concentrations. SBI-810 concentrations producing nonspecific effects on G protein sensors are not presented. SBI-553 and SBI-810 are comparable in these assays. **(C)** SBI-810 and SBI-553-induced recruitment of human β-arrestin2 to the human NTSR1 was assessed in HEK293T cells by BRET. Compounds are equally potent and efficacious in this assay. **(D)** G protein activation by the NTSR1 induced by co-treatment with NT and vehicle or 0.03, 0.3, 1, 3, or 10 μM SBI-810. The effect of SBI-810 on NT-induced G protein activation is similar to that of SBI-553, as presented in main text Figure 5. As the doses evaluated, unlike SBI-553, SBI-810 did not stimulate GoA activation in the absence of NT. **(E)** β-arrestin2 recruitment to NTSR1 induced by co-treatment with NT and vehicle or 0.03, 0.3, 1, 3, or 30 μM SBI-810. SBI-810 acted as a full β-arrestin2 agonist and permitted NT-induced β-arrestin2 recruitment. HEK293T cells, N=3-5.

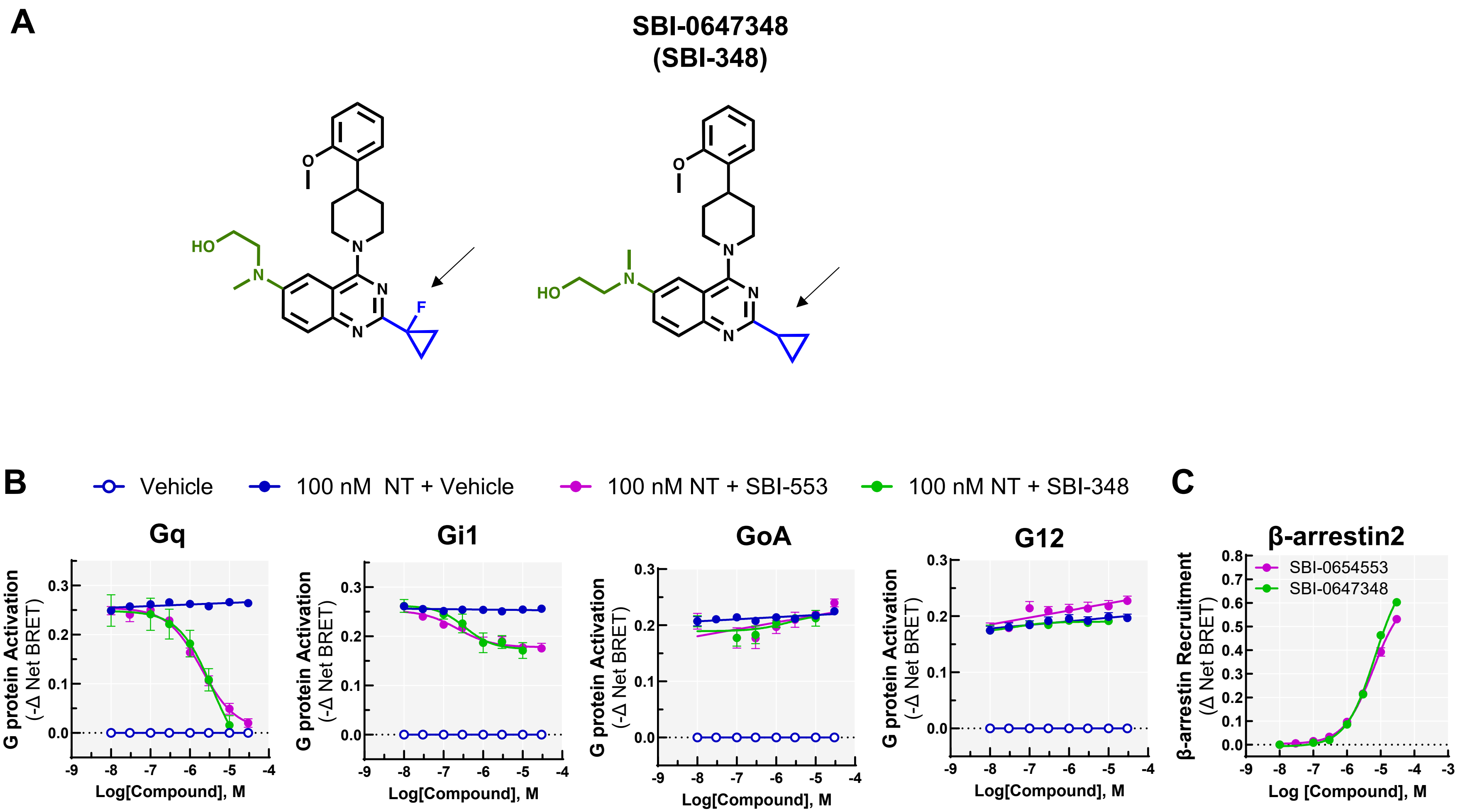

**Supplemental Figure 9. Removing SBI-553's F has no effect on G protein selectivity. (A)** Structural comparison of SBI-553 and SBI-348. Arrows indicate the fluorine that differs between SBI-553 and SBI-348. **(B)** In HEK293T cells transiently expressing NTSR1, SBI-348 (30 nM – 10 μM) and SBI-553 (30 nM – 30 μM) were assessed for their ability to antagonize NT-induced G protein activation using the bioluminescence resonance energy transfer 2-based TRUPATH assay. SBI-553 and SBI-348 are comparable in these assays. **(C)** SBI-348 and SBI-553 (30 nM – 30 μM)-induced recruitment of human β-arrestin2 to the human NTSR1 was assessed in HEK293T cells by BRET. Compounds are equally potent and efficacious in these assays. N = 3.

**Table S1. Curve parameters and statistical comparisons presented in Figure 1 – Supporting Figure 1.**

| Fig | Panel | Description | N | Condition | Parameter | Mean (95% CI) | Units |
|---|---|---|---|---|---|---|---|
| **1** | **A** | Illustration | | | | | |
| **1** | **B** | Illustration | | | | | |
| **1** | **C** | TRUPATH in HEK293T cells, NTSR1 ligand-induced activation of $G_{\alpha}q$ | 3 | NT | $EC_{50}$ | 3.009e-009 (2.186e-009 to 4.142e-009) | M |
| | | | | | Top | 0.3131 (0.3004 to 0.3258) | -Δ Net BRET |
| | | | | PD149163 | $EC_{50}$ | 9.301e-007 (6.730e-007 to 1.285e-006) | M |
| | | | | | Top | 0.3367 (0.3131 to 0.3602) | -Δ Net BRET |
| | | | | SBI-553 | Slope | -0.001859 (-0.003554 to -0.0001651) | M |
| | | | | | y-intercept | -0.01661 (-0.02803 to -0.005187) | -Δ Net BRET |
| | | | | SR142948A | Slope | -0.002316 (-0.003985 to -0.0006460) | M |
| | | | | | y-intercept | -0.02105 (-0.03445 to -0.007658) | -Δ Net BRET |
| | | TRUPATH in HEK293T cells, NTSR1 ligand-induced activation of $G_{\alpha}11$ | 3 | NT | $EC_{50}$ | 1.472e-009 (1.051e-009 to 2.063e-009) | M |
| | | | | | Top | 0.2933 (0.2807 to 0.3058) | -Δ Net BRET |
| | | | | PD149163 | $EC_{50}$ | 1.549e-007 (1.327e-007 to 1.807e-007) | M |
| | | | | | Top | 0.2934 (0.2850 to 0.3018) | -Δ Net BRET |
| | | | | SBI-553 | $EC_{50}$ | 7.648e-005 (5.192e-006 to 0.001127) | M |
| | | | | | Top | -0.06187 (-0.1445 to 0.02075) | -Δ Net BRET |
| | | | | SR142948A | Slope | 0.0009587(-0.0003953 to 0.002313) | M |
| | | | | | y-intercept | 0.01166 (0.0007933 to 0.02252) | -Δ Net BRET |
| | | TRUPATH in HEK293T cells, NTSR1 ligand-induced activation of $G_{\alpha}15$ | 3 | NT | $EC_{50}$ | 1.859e-009 (1.386e-009 to 2.494e-009) | M |
| | | | | | Top | 0.1630 (0.1564 to 0.1696) | -Δ Net BRET |
| | | | | PD149163 | $EC_{50}$ | 2.301e-007 (1.773e-007 to 2.986e-007) | M |
| | | | | | Top | 0.1734 (0.1651 to 0.1817) | -Δ Net BRET |
| | | | | SBI-553 | $EC_{50}$ | 1.506e-006 (5.688e-007 to 3.988e-006) | M |
| | | | | | Top | 0.08478 (0.06740 to 0.1022) | -Δ Net BRET |
| | | | | SR142948A | Slope | 0.002852 (-0.0002553 to 0.005960) | M |
| | | | | | y-intercept | 0.02919 (0.004257 to 0.05412) | -Δ Net BRET |
| | | TRUPATH in HEK293T cells, NTSR1 ligand-induced activation of $G_{\alpha}i1$ | 3 | NT | $EC_{50}$ | 8.383e-009 (6.000e-009 to 1.171e-008) | M |
| | | | | | Top | 0.2948 (0.2795 to 0.3101) | -Δ Net BRET |
| | | | | PD149163 | $EC_{50}$ | 3.177e-006 (2.056e-006 to 4.907e-006) | M |
| | | | | | Top | 0.2811 (0.2455 to 0.3166) | -Δ Net BRET |
| | | | | SBI-553 | $EC_{50}$ | 2.297e-005 (1.306e-005 to 4.041e-005) | M |
| | | | | | Top | 0.1583 (0.1262 to 0.1903) | -Δ Net BRET |
| | | | | SR142948A | Slope | -0.003421 (-0.006302 to -0.0005398) | M |
| | | | | | y-intercept | -0.02446 (-0.04758 to -0.001344) | -Δ Net BRET |
| | | TRUPATH in HEK293T cells, NTSR1 ligand-induced | 3 | NT | $EC_{50}$ | 1.341e-008 (9.697e-009 to 1.855e-008) | M |
| | | | | | Top | 0.2582 (0.2448 to 0.2716) | -Δ Net BRET |
| | | | | PD149163 | $EC_{50}$ | 2.243e-006 (1.775e-006 to 2.835e-006) | M |

| | | | | | | |
|---|---|---|---|---|---|---|
| | activation of $G_{\alpha}i2$ | | | Top | 0.2947 (0.2760 to 0.3134) | -Δ Net BRET |
| | | | SBI-553 | $EC_{50}$ | 5.167e-005 (2.664e-005 to 0.0001002) | M |
| | | | | Top | 0.2003 (0.1395 to 0.2611) | -Δ Net BRET |
| | | | SR142948A | Slope | -0.0005653 (-0.002768 to 0.001638) | M |
| | | | | y-intercept | 0.005272 (-0.01240 to 0.02295) | -Δ Net BRET |
| | TRUPATH in HEK293T cells, NTSR1 ligand-induced activation of $G_{\alpha}i3$ | 3 | NT | $EC_{50}$ | 1.753e-008 (1.245e-008 to 2.469e-008) | M |
| | | | | Top | 0.3837 (0.3626 to 0.4047) | -Δ Net BRET |
| | | | PD149163 | $EC_{50}$ | 5.864e-006 (3.152e-006 to 1.091e-005) | M |
| | | | | Top | 0.2392 (0.1891 to 0.2894) | -Δ Net BRET |
| | | | SBI-553 | $EC_{50}$ | 1.066e-005 (4.636e-006 to 2.449e-005) | M |
| | | | | Top | 0.08170 (0.06165 to 0.1017) | -Δ Net BRET |
| | | | SR142948A | Slope | 0.001546 (-0.001045 to 0.004137) | M |
| | | | | y-intercept | 0.03113 (0.01033 to 0.05192) | -Δ Net BRET |
| | TRUPATH in HEK293T cells, NTSR1 ligand-induced activation of $G_{\alpha}oA$ | 3 | NT | $EC_{50}$ | 1.000e-008 (6.526e-009 to 1.533e-008) | M |
| | | | | Top | 0.2628 (0.2450 to 0.2805) | -Δ Net BRET |
| | | | PD149163 | $EC_{50}$ | 3.372e-006 (2.770e-006 to 4.105e-006) | M |
| | | | | Top | 0.2578 (0.2429 to 0.2728) | -Δ Net BRET |
| | | | SBI-553 | $EC_{50}$ | 2.409e-005 (1.195e-005 to 4.855e-005) | M |
| | | | | Top | 0.1718 (0.1280 to 0.2156) | -Δ Net BRET |
| | | | SR142948A | Slope | -0.005954 (-0.008622 to -0.003285) | M |
| | | | | y-intercept | -0.06270 (-0.08411 to -0.04129) | -Δ Net BRET |
| | TRUPATH in HEK293T cells, NTSR1 ligand-induced activation of $G_{\alpha}oB$ | 4 | NT | $EC_{50}$ | 2.083e-008 (1.198e-008 to 3.623e-008) | M |
| | | | | Top | 0.2591 (0.2361 to 0.2821) | -Δ Net BRET |
| | | | PD149163 | $EC_{50}$ | 3.370e-006 (2.656e-006 to 4.276e-006) | M |
| | | | | Top | 0.2511 (0.2335 to 0.2687) | -Δ Net BRET |
| | | | SBI-553 | $EC_{50}$ | 9.556e-006 (6.521e-006 to 1.400e-005) | M |
| | | | | Top | 0.1714 (0.1525 to 0.1902) | -Δ Net BRET |
| | | | SR142948A | Slope | -0.002522 (-0.006699 to 0.001654) | M |
| | | | | y-intercept | -0.02835 (-0.06186 to 0.005159) | -Δ Net BRET |
| | TRUPATH in HEK293T cells, NTSR1 ligand-induced activation of $G_{\alpha}z$ | 3 | NT | $EC_{50}$ | 1.147e-008 (8.889e-009 to 1.479e-008) | M |
| | | | | Top | 0.3014 (0.2892 to 0.3137) | -Δ Net BRET |
| | | | PD149163 | $EC_{50}$ | 1.929e-006 (1.642e-006 to 2.267e-006) | M |
| | | | | Top | 0.2995 (0.2868 to 0.3122) | -Δ Net BRET |
| | | | SBI-553 | $EC_{50}$ | 2.489e-005 (7.106e-006 to 8.036e-005) | M |
| | | | | Top | 0.07472 (0.05158 to 0.1216) | -Δ Net BRET |
| | | | SR142948A | Slope | 0.0006605 (-0.0005994 to 0.001920) | M |
| | | | | y-intercept | 0.02037 (0.01026 to 0.03048) | -Δ Net BRET |
| | TRUPATH in HEK293T cells, NTSR1 ligand- | 4 | NT | $EC_{50}$ | 1.083e-008 (7.413e-009 to 1.581e-008) | M |
| | | | | Top | 0.09705 (0.09121 to 0.1029) | -Δ Net BRET |

| | | | | | | | |
|---|---|---|---|---|---|---|---|
| | | induced activation of $G_{\alpha}$gustducin | | PD149163 | $EC_{50}$ | 1.432e-007 (6.753e-008 to 3.035e-007) | M |
| | | | | | Top | 0.04144 (0.03621 to 0.04668) | -Δ Net BRET |
| | | | | SBI-553 | $EC_{50}$ | 3.591e-006 (1.586e-006 to 8.131e-006) | M |
| | | | | | Top | 0.03890 (0.03131 to 0.04649) | -Δ Net BRET |
| | | | | SR142948A | Slope | -0.0002241 (-0.001052 to 0.0006043) | M |
| | | | | | y-intercept | 0.003942 (-0.002704 to 0.01059) | -Δ Net BRET |
| | | TRUPATH in HEK293T cells, NTSR1 ligand-induced activation of $G_{\alpha}12$ | 3 | NT | $EC_{50}$ | 2.556e-008 (1.635e-008 to 3.996e-008) | M |
| | | | | | Top | 0.3809 (0.3534 to 0.4083) | -Δ Net BRET |
| | | | | PD149163 | $EC_{50}$ | 5.780e-006 (3.054e-006 to 1.094e-005) | M |
| | | | | | Top | 0.2549 (0.2002 to 0.3096) | -Δ Net BRET |
| | | | | SBI-553 | $EC_{50}$ | 1.742e-005 (8.940e-006 to 3.337e-005) | M |
| | | | | | Top | 0.2533 (0.2064 to 0.3172) | -Δ Net BRET |
| | | | | SR142948A | $EC_{50}$ | 3.860e-006 (1.247e-007 to 0.0001195) | M |
| | | | | | Top | -0.02221 (-0.04421 to -0.0002191) | -Δ Net BRET |
| | | TRUPATH in HEK293T cells, NTSR1 ligand-induced activation of $G_{\alpha}13$ | 3 | NT | $EC_{50}$ | 1.312e-008 (1.075e-008 to 1.602e-008) | M |
| | | | | | Top | 0.5418 (0.5246 to 0.5591) | -Δ Net BRET |
| | | | | PD149163 | $EC_{50}$ | 1.360e-005 (9.863e-006 to 1.876e-005) | M |
| | | | | | Top | 0.6957 (0.5979 to 0.7935) | -Δ Net BRET |
| | | | | SBI-553 | $EC_{50}$ | 2.810e-005 (2.135e-005 to 3.727e-005) | M |
| | | | | | Top | 0.4470 (0.4035 to 0.4991) | -Δ Net BRET |
| | | | | SR142948A | Slope | 0.006624 (0.004868 to 0.008379) | M |
| | | | | | y-intercept | 0.07884 (0.06475 to 0.09292) | -Δ Net BRET |
| | | TRUPATH in HEK293T cells, NTSR1 ligand-induced activation of $G_{\alpha}$sS | 4 | NT | Slope | 0.002161 (-0.003153 to 0.007476) | M |
| | | | | | y-intercept | 0.02633 (-0.02045 to 0.07312) | -Δ Net BRET |
| | | | | PD149163 | Slope | -0.003440 (-0.007835 to 0.0009548) | M |
| | | | | | y-intercept | -0.02678 (-0.05775 to 0.004186) | -Δ Net BRET |
| | | | | SBI-553 | Slope | -0.002718 (-0.006169 to 0.0007334) | M |
| | | | | | y-intercept | -0.01960 (-0.04286 to 0.003665) | -Δ Net BRET |
| | | | | SR142948A | Slope | 3.113e-005 (-0.003859 to 0.003922) | M |
| | | | | | y-intercept | 0.02107 (-0.009868 to 0.05201) | -Δ Net BRET |
| | | TRUPATH in HEK293T cells, NTSR1 ligand-induced activation of $G_{\alpha}$sL | 3 | NT | Slope | -0.003200 (-0.008678 to 0.002277) | M |
| | | | | | y-intercept | 0.002161 (-0.04606 to 0.05038) | -Δ Net BRET |
| | | | | PD149163 | Slope | 0.003582 (6.715e-005 to 0.007097) | M |
| | | | | | y-intercept | 0.05046 (0.02570 to 0.07523) | -Δ Net BRET |
| | | | | SBI-553 | Slope | -0.002897 (-0.007035 to 0.001242) | M |
| | | | | | y-intercept | -0.02070 (-0.04860 to 0.007195) | -Δ Net BRET |
| | | | | SR142948A | Slope | 0.001585 (-0.002138 to 0.005308) | M |
| | | | | | y-intercept | 0.03624 (0.006632 to 0.06585) | -Δ Net BRET |
| **1** | **D** | Maximal NTSR1 ligand-induced G protein activation by TRUPATH, two-way ANOVA followed by Tukey multiple comparisons tests. N = 3 | | | | | |

| | | | | | | | |
|---|---|---|---|---|---|---|---|
| | | $F_{G\ protein}$ (13, 490) = 140.1 P<0.0001<br>$F_{Ligand}$ (3, 490) = 1028 P<0.0001<br>$F_{Interaction}$ (39, 490) = 33.75 P<0.0001<br><br>Colored asterisks over each bar indicate the treatments from which that compound significantly differed at that G protein. *p<0.05, Treatment vs NT (*), SR142948A (*), PD149163 (*), SBI-553(*) | | | | | |
| **1** | **E** | Maximal SBI-553-induced G protein activation by TRUPATH in NTSR1 expressing and non-expressing cells, two-way ANOVA followed by Bonferroni multiple comparisons tests. N = 3<br><br>$F_{G\ protein}$ (7, 127) = 38.09 P<0.0001<br>$F_{Cell\text{-}type}$ (1, 127) = 625.2 P<0.0001<br>$F_{Interaction}$ (7, 127) = 9.236 P<0.0001<br><br>*p<0.0001 | | | | | |
| **1** | **F** | Illustration | | | | | |
| **1** | **G** | NTSR1 ligand-induced activation of β-arrestin1 | 3 | NT | $EC_{50}$ | 8.594e-009 (6.806e-009 to 1.085e-008) | M |
| | | | | | Top | 0.4769 (0.4596 to 0.4942) | Δ Net BRET |
| | | | | PD149163 | $EC_{50}$ | 1.247e-006 (9.151e-007 to 1.698e-006) | M |
| | | | | | Top | 0.3592 (0.3322 to 0.3861) | Δ Net BRET |
| | | | | SBI-553 | $EC_{50}$ | 2.043e-005 (1.716e-005 to 2.433e-005) | M |
| | | | | | Top | 0.5666 (0.5322 to 0.6010) | Δ Net BRET |
| | | | | SR142948A | Slope | 0.0003879 (0.0001093 to 0.0006665) | M |
| | | | | | y-intercept | 0.004380 (0.002145 to 0.006616) | Δ Net BRET |
| | | NTSR1 ligand-induced activation of β-arrestin2 | 3 | NT | $EC_{50}$ | 8.000e-009 (6.901e-009 to 9.275e-009) | M |
| | | | | | Top | 0.5878 (0.5744 to 0.6013) | Δ Net BRET |
| | | | | PD149163 | $EC_{50}$ | 3.616e-007 (2.925e-007 to 4.471e-007) | M |
| | | | | | Top | 0.5482 (0.5250 to 0.5714) | Δ Net BRET |
| | | | | SBI-553 | $EC_{50}$ | 1.423e-005 (1.199e-005 to 1.689e-005) | M |
| | | | | | Top | 0.7275 (0.6881 to 0.7669) | Δ Net BRET |
| | | | | SR142948A | Slope | 0.0009235 (-0.001006 to 0.002853) | M |
| | | | | | y-intercept | 0.01039 (-0.005099 to 0.02587) | Δ Net BRET |
| **1** | **H** | Radar plots | | | | | |
| **1** | **I** | Illustration | | | | | |
| **1** | **J, left** | NT-induced G protein activation by TGFα Shedding | 3 | $G_{\alpha}q$ | $EC_{50}$ | 9.592e-011 (7.268e-011 to 1.271e-010) | M |
| | | | | | Top | 21.38 (20.62 to 22.14) | % AP Shedding |
| | | | 3 | $G_{\alpha}14$ | $EC_{50}$ | 1.462e-010 (8.506e-011 to 2.594e-010) | M |
| | | | | | Top | 17.08 (15.80 to 18.40) | % AP Shedding |
| | | | 3 | $G_{\alpha}16$ | $EC_{50}$ | 1.241e-008 (8.276e-009 to 1.885e-008) | M |
| | | | | | Top | 17.74 (16.43 to 19.11) | % AP Shedding |
| | | | 3 | $G_{\alpha}i1/2$ | $EC_{50}$ | 8.692e-010 (6.207e-010 to 1.214e-009) | M |
| | | | | | Top | 21.31 (20.27 to 22.36) | % AP Shedding |
| | | | 3 | $G_{\alpha}o$ | $EC_{50}$ | 1.334e-009 (7.374e-010 to 2.481e-009) | M |
| | | | | | Top | 13.48 (12.07 to 14.98) | % AP Shedding |
| | | | 3 | $G_{\alpha}olf$ | $EC_{50}$ | 2.120e-009 (7.402e-010 to 7.127e-009) | M |
| | | | | | Top | 9.830 (8.177 to 11.91) | % AP Shedding |

| | | | | | | | |
|---|---|---|---|---|---|---|---|
| | | | 3 | $G_{\alpha}z$ | $EC_{50}$ | 3.083e-010 (9.698e-011 to 9.448e-010) | M |
| | | | | | Top | 7.338 (5.943 to 8.821) | % AP Shedding |
| | | | 3 | $G_{\alpha}s$ | $EC_{50}$ | 4.314e-009 (3.562e-009 to 5.211e-009) | M |
| | | | | | Top | -0.1547 (-0.6071 to 0.2952) | % AP Shedding |
| | | | 3 | $G_{\alpha}12$ | $EC_{50}$ | 1.165e-009 (5.450e-010 to 2.289e-009) | M |
| | | | | | Top | 13.48 (11.68 to 15.40) | % AP Shedding |
| | | | 3 | $G_{\alpha}13$ | $EC_{50}$ | 1.155e-009 (5.568e-010 to 2.475e-009) | M |
| | | | | | Top | 14.00 (12.27 to 15.84) | % AP Shedding |
| | | | 3 | $G_{\alpha}\Delta C$ | $EC_{50}$ | 1.288e-010 (1.778e-011 to 1.328e-009) | M |
| | | | | | Top | 0.9097 (0.5226 to 1.329) | % AP Shedding |
| **1** | **J, right** | SBI-553-induced G protein activation by TGFα Shedding | 3 | $G_{\alpha}q$ | $EC_{50}$ | 4.030e-007 (ND) | M |
| | | | | | Top | 0.8005 (ND) | % AP Shedding |
| | | | 3 | $G_{\alpha}14$ | $EC_{50}$ | 7.126e-006 (ND) | M |
| | | | | | Top | 7.408 (ND) | % AP Shedding |
| | | | 3 | $G_{\alpha}16$ | $EC_{50}$ | ~0.007718 (ND) | M |
| | | | | | Top | ~2632 (ND) | % AP Shedding |
| | | | 3 | $G_{\alpha}i1/2$ | $EC_{50}$ | 1.661e-005 (ND) | M |
| | | | | | Top | 8.721 (ND) | % AP Shedding |
| | | | 3 | $G_{\alpha}o$ | $EC_{50}$ | 2.447e-006 (1.043e-006 to 1.129e-005) | M |
| | | | | | Top | 9.421 (7.279 to 16.13) | % AP Shedding |
| | | | 3 | $G_{\alpha}olf$ | $EC_{50}$ | 1.321e-007 (2.634e-008 to 8.655e-007) | M |
| | | | | | Top | 1.244 (0.8548 to 1.730) | % AP Shedding |
| | | | 3 | $G_{\alpha}z$ | $EC_{50}$ | ~0.001822 (ND) | M |
| | | | | | Top | ~714.1 (ND) | % AP Shedding |
| | | | 3 | $G_{\alpha}s$ | $EC_{50}$ | 3.354e-006 (ND) | M |
| | | | | | Top | 0.3948 (ND) | % AP Shedding |
| | | | 3 | $G_{\alpha}12$ | $EC_{50}$ | 3.165e-007 (1.225e-007 to 8.086e-007) | M |
| | | | | | Top | 10.64 (8.278 to 13.27) | % AP Shedding |
| | | | 3 | $G_{\alpha}13$ | $EC_{50}$ | 5.878e-006 (ND) | M |
| | | | | | Top | 12.66 (ND) | % AP Shedding |
| | | | 3 | $G_{\alpha}\Delta C$ | $EC_{50}$ | 2.482e-008 (ND) | M |
| | | | | | Top | 0.209 (ND) | % AP Shedding |
| **1** | **K** | Maximal NTSR1 ligand-induced G protein activation by AP-TGFα shedding assay, two-way ANOVA followed by Bonferroni multiple comparisons test. N = 3<br><br>$F_{G\ protein}(11, 263) = 17.50$ P<0.0001<br>$F_{Ligand}(1, 263) = 435.4$ P<0.0001<br>$F_{Interaction}(11, 263) = 17.57$ P<0.0001<br><br>*p<0.05, as compared to GΔC, unless otherwise indicated | | | | | |

CI, confidence interval; NT, neurotensin; SBI-553, SBI-0654553; $EC_{50}$, half-maximal effective concentration; ND, not defined

### Table S2. Curve parameters and statistical comparisons presented in Figure 2 – Supporting Figure 2.

| Fig | Panel | Description | N | Condition | Parameter | Mean (95% CI) | Units |
|---|---|---|---|---|---|---|---|
| **2** | **A** | Illustration | | | | | |
| **2** | **B** | Illustration | | | | | |
| **2** | **C** | Effect of SR142948A on NT-induced β-arrestin1 recruitment | 3 | 0 nM SR142948A | $EC_{50}$ | 1.521e-008 (1.074e-008 to 2.156e-008) | M |
| | | | | | Top | 0.4559 (0.4396 to 0.4723) | -Δ Net BRET |
| | | | | 10 nM SR142948A | $EC_{50}$ | 3.074e-008 (2.136e-008 to 4.424e-008) | M |
| | | | | | Top | Shared | -Δ Net BRET |
| | | | | 100 nM SR142948A | $EC_{50}$ | 1.281e-007 (9.077e-008 to 1.809e-007) | M |
| | | | | | Top | Shared | -Δ Net BRET |
| | | | | 1 µM SR142948A | $EC_{50}$ | 6.967e-007 (4.925e-007 to 9.855e-007) | M |
| | | | | | Top | Shared | -Δ Net BRET |
| | | | | 10 µM SR142948A | $EC_{50}$ | 4.454e-006 (3.113e-006 to 6.371e-006) | M |
| | | | | | Top | Shared | -Δ Net BRET |
| | | | | 100 µM SR142948A | $EC_{50}$ | 1.974e-005 (1.323e-005 to 2.947e-005) | M |
| | | | | | Top | Shared | -Δ Net BRET |
| | | Effect of SR142948A on NT-induced β-arrestin2 recruitment | 3 | 0 nM SR142948A | $EC_{50}$ | 7.796e-009 (6.234e-009 to 9.750e-009) | M |
| | | | | | Top | 0.5180 (0.5065 to 0.5294) | -Δ Net BRET |
| | | | | 10 nM SR142948A | $EC_{50}$ | 2.015e-008 (1.600e-008 to 2.538e-008) | M |
| | | | | | Top | Shared | -Δ Net BRET |
| | | | | 100 nM SR142948A | $EC_{50}$ | 1.014e-007 (8.124e-008 to 1.266e-007) | M |
| | | | | | Top | Shared | -Δ Net BRET |
| | | | | 1 µM SR142948A | $EC_{50}$ | 4.048e-007 (3.204e-007 to 5.115e-007) | M |
| | | | | | Top | Shared | -Δ Net BRET |
| | | | | 10 µM SR142948A | $EC_{50}$ | 3.147e-006 (2.489e-006 to 3.978e-006) | M |
| | | | | | Top | Shared | -Δ Net BRET |
| | | | | 100 µM SR142948A | $EC_{50}$ | 1.469e-005 (1.156e-005 to 1.867e-005) | M |
| | | | | | Top | Shared | -Δ Net BRET |
| **2** | **D** | Effect of SR142948A on NT-induced activation of $G_{\alpha}q$ | 3 | 0 nM SR142948A | $EC_{50}$ | 3.301e-009 (2.304e-009 to 4.729e-009) | M |
| | | | | | Top | 0.3204 (0.3108 to 0.3300) | -Δ Net BRET |
| | | | | 10 nM SR142948A | $EC_{50}$ | 5.654e-009 (3.988e-009 to 8.015e-009) | M |
| | | | | | Top | Shared | -Δ Net BRET |
| | | | | 100 nM SR142948A | $EC_{50}$ | 3.537e-008 (2.470e-008 to 5.065e-008) | M |
| | | | | | Top | Shared | -Δ Net BRET |
| | | | | 1 µM SR142948A | $EC_{50}$ | 8.479e-008 (6.035e-008 to 1.191e-007) | M |
| | | | | | Top | Shared | -Δ Net BRET |
| | | | | 10 µM SR142948A | $EC_{50}$ | 1.007e-006 (7.179e-007 to 1.412e-006) | M |
| | | | | | Top | Shared | -Δ Net BRET |
| | | | | 100 µM SR142948A | $EC_{50}$ | 6.564e-006 (4.636e-006 to 9.295e-006) | M |

| | | | | | | |
|---|---|---|---|---|---|---|
| | | | | Top | Shared | -Δ Net BRET |
| | Effect of SR142948A on NT-induced activation of $G_{\alpha}11$ | 3 | 0 nM SR142948A | $EC_{50}$ | 2.095e-009 (1.536e-009 to 2.856e-009) | M |
| | | | | Top | 0.2658 (0.2587 to 0.2730) | -Δ Net BRET |
| | | | 10 nM SR142948A | $EC_{50}$ | 8.888e-009 (6.447e-009 to 1.225e-008) | M |
| | | | | Top | Shared | -Δ Net BRET |
| | | | 100 nM SR142948A | $EC_{50}$ | 2.697e-008 (1.969e-008 to 3.694e-008) | M |
| | | | | Top | Shared | -Δ Net BRET |
| | | | 1 µM SR142948A | $EC_{50}$ | 1.071e-007 (7.953e-008 to 1.442e-007) | M |
| | | | | Top | Shared | -Δ Net BRET |
| | | | 10 µM SR142948A | $EC_{50}$ | 7.059e-007 (5.225e-007 to 9.536e-007) | M |
| | | | | Top | Shared | -Δ Net BRET |
| | | | 100 µM SR142948A | $EC_{50}$ | 4.171e-006 (3.050e-006 to 5.704e-006) | M |
| | | | | Top | Shared | -Δ Net BRET |
| | Effect of SR142948A on NT-induced activation of $G_{\alpha}15$ | 3 | 0 nM SR142948A | $EC_{50}$ | 2.530e-009 (1.764e-009 to 3.629e-009) | M |
| | | | | Top | 0.1377 (0.1306 to 0.1447) | -Δ Net BRET |
| | | | 10 nM SR142948A | $EC_{50}$ | 8.222e-009 (5.832e-009 to 1.159e-008) | M |
| | | | | Top | Shared | -Δ Net BRET |
| | | | 100 nM SR142948A | $EC_{50}$ | 3.429e-008 (2.391e-008 to 4.916e-008) | M |
| | | | | Top | Shared | -Δ Net BRET |
| | | | 1 µM SR142948A | $EC_{50}$ | 1.370e-007 (9.696e-008 to 1.936e-007) | M |
| | | | | Top | Shared | -Δ Net BRET |
| | | | 10 µM SR142948A | $EC_{50}$ | 8.528e-007 (5.711e-007 to 1.274e-006) | M |
| | | | | Top | Shared | -Δ Net BRET |
| | | | 100 µM SR142948A | $EC_{50}$ | 6.366e-006 (2.529e-006 to 1.602e-005) | M |
| | | | | Top | Shared | -Δ Net BRET |
| | Effect of SR142948A on NT-induced activation of $G_{\alpha}i1$ | 3 | 0 nM SR142948A | $EC_{50}$ | 1.155e-008 (7.537e-009 to 1.769e-008) | M |
| | | | | Top | 0.2965 (0.2847 to 0.3083) | -Δ Net BRET |
| | | | 10 nM SR142948A | $EC_{50}$ | 1.879e-008 (1.210e-008 to 2.917e-008) | M |
| | | | | Top | Shared | -Δ Net BRET |
| | | | 100 nM SR142948A | $EC_{50}$ | 5.576e-008 (3.595e-008 to 8.649e-008) | M |
| | | | | Top | Shared | -Δ Net BRET |
| | | | 1 µM SR142948A | $EC_{50}$ | 2.804e-007 (1.720e-007 to 4.572e-007) | M |
| | | | | Top | Shared | -Δ Net BRET |
| | | | 10 µM SR142948A | $EC_{50}$ | 1.994e-006 (1.237e-006 to 3.216e-006) | M |
| | | | | Top | Shared | -Δ Net BRET |
| | | | 100 µM SR142948A | $EC_{50}$ | 1.789e-005 (1.102e-005 to 2.904e-005) | M |
| | | | | Top | Shared | -Δ Net BRET |
| | Effect of SR142948A on NT-induced activation of | 3 | 0 nM SR142948A | $EC_{50}$ | 6.891e-009 (4.193e-009 to 1.133e-008) | M |
| | | | | Top | 0.2365 (0.2235 to 0.2495) | -Δ Net BRET |

| | | | | | | |
|---|---|---|---|---|---|---|
| | $G_{\alpha}i2$ | | 10 nM SR142948A | $EC_{50}$ | 1.670e-008 (1.012e-008 to 2.756e-008) | M |
| | | | | Top | Shared | -Δ Net BRET |
| | | | 100 nM SR142948A | $EC_{50}$ | 3.888e-008 (2.317e-008 to 6.523e-008) | M |
| | | | | Top | Shared | -Δ Net BRET |
| | | | 1 µM SR142948A | $EC_{50}$ | 2.413e-007 (1.442e-007 to 4.037e-007) | M |
| | | | | Top | Shared | -Δ Net BRET |
| | | | 10 µM SR142948A | $EC_{50}$ | 2.306e-006 (1.383e-006 to 3.845e-006) | M |
| | | | | Top | Shared | -Δ Net BRET |
| | | | 100 µM SR142948A | $EC_{50}$ | 7.712e-006 (4.683e-006 to 1.270e-005) | M |
| | | | | Top | Shared | -Δ Net BRET |
| | Effect of SR142948A on NT-induced activation of $G_{\alpha}i3$ | 3 | 0 nM SR142948A | $EC_{50}$ | 1.960e-008 (1.512e-008 to 2.541e-008) | M |
| | | | | Top | 0.4220 (0.4104 to 0.4337) | -Δ Net BRET |
| | | | 10 nM SR142948A | $EC_{50}$ | 5.137e-008 (3.960e-008 to 6.663e-008) | M |
| | | | | Top | Shared | -Δ Net BRET |
| | | | 100 nM SR142948A | $EC_{50}$ | 1.690e-007 (1.308e-007 to 2.184e-007) | M |
| | | | | Top | Shared | -Δ Net BRET |
| | | | 1 µM SR142948A | $EC_{50}$ | 8.335e-007 (6.487e-007 to 1.071e-006) | M |
| | | | | Top | Shared | -Δ Net BRET |
| | | | 10 µM SR142948A | $EC_{50}$ | 5.731e-006 (4.433e-006 to 7.410e-006) | M |
| | | | | Top | Shared | -Δ Net BRET |
| | | | 100 µM SR142948A | $EC_{50}$ | 2.075e-005 (1.543e-005 to 2.792e-005) | M |
| | | | | Top | Shared | -Δ Net BRET |
| | Effect of SR142948A on NT-induced activation of $G_{\alpha}oA$ | 3 | 0 nM SR142948A | $EC_{50}$ | 1.188e-008 (8.965e-009 to 1.575e-008) | M |
| | | | | Top | 0.2953 (0.2871 to 0.3036) | -Δ Net BRET |
| | | | 10 nM SR142948A | $EC_{50}$ | 4.005e-008 (2.977e-008 to 5.388e-008) | M |
| | | | | Top | Shared | -Δ Net BRET |
| | | | 100 nM SR142948A | $EC_{50}$ | 9.890e-008 (7.468e-008 to 1.310e-007) | M |
| | | | | Top | Shared | -Δ Net BRET |
| | | | 1 µM SR142948A | $EC_{50}$ | 4.795e-007 (3.578e-007 to 6.426e-007) | M |
| | | | | Top | Shared | -Δ Net BRET |
| | | | 10 µM SR142948A | $EC_{50}$ | 2.446e-006 (1.823e-006 to 3.282e-006) | M |
| | | | | Top | Shared | -Δ Net BRET |
| | | | 100 µM SR142948A | $EC_{50}$ | 1.233e-005 (9.197e-006 to 1.653e-005) | M |
| | | | | Top | Shared | -Δ Net BRET |
| | Effect of SR142948A on NT-induced activation of $G_{\alpha}oB$ | 4 | 0 nM SR142948A | $EC_{50}$ | 2.445e-008 (1.755e-008 to 3.406e-008) | M |
| | | | | Top | 0.2818 (0.2721 to 0.2914) | -Δ Net BRET |
| | | | 10 nM SR142948A | $EC_{50}$ | 5.159e-008 (3.720e-008 to 7.155e-008) | M |
| | | | | Top | Shared | -Δ Net BRET |
| | | | 100 nM SR142948A | $EC_{50}$ | 1.480e-007 (1.076e-007 to 2.037e-007) | M |

| | | | | | | |
|---|---|---|---|---|---|---|
| | | | | Top | Shared | -Δ Net BRET |
| | | | 1 μM SR142948A | $EC_{50}$ | 5.734e-007 (4.149e-007 to 7.923e-007) | M |
| | | | | Top | Shared | -Δ Net BRET |
| | | | 10 μM SR142948A | $EC_{50}$ | 2.767e-006 (1.987e-006 to 3.853e-006) | M |
| | | | | Top | Shared | -Δ Net BRET |
| | | | 100 μM SR142948A | $EC_{50}$ | 1.176e-005 (8.494e-006 to 1.628e-005) | M |
| | | | | Top | Shared | -Δ Net BRET |
| | Effect of SR142948A on NT-induced activation of $G_{\alpha}z$ | 3 | 0 nM SR142948A | $EC_{50}$ | 8.419e-009 (6.582e-009 to 1.077e-008) | M |
| | | | | Top | 0.2930 (0.2857 to 0.3003) | -Δ Net BRET |
| | | | 10 nM SR142948A | $EC_{50}$ | 2.706e-008 (2.088e-008 to 3.507e-008) | M |
| | | | | Top | Shared | -Δ Net BRET |
| | | | 100 nM SR142948A | $EC_{50}$ | 7.732e-008 (6.040e-008 to 9.898e-008) | M |
| | | | | Top | Shared | -Δ Net BRET |
| | | | 1 μM SR142948A | $EC_{50}$ | 4.139e-007 (3.198e-007 to 5.357e-007) | M |
| | | | | Top | Shared | -Δ Net BRET |
| | | | 10 μM SR142948A | $EC_{50}$ | 2.848e-006 (2.200e-006 to 3.688e-006) | M |
| | | | | Top | Shared | -Δ Net BRET |
| | | | 100 μM SR142948A | $EC_{50}$ | 1.500e-005 (1.150e-005 to 1.956e-005) | M |
| | | | | Top | Shared | -Δ Net BRET |
| | Effect of SR142948A on NT-induced activation of $G_{\alpha}$gustducin | 4 | 0 nM SR142948A | $EC_{50}$ | 1.585e-008 (8.262e-009 to 3.042e-008) | M |
| | | | | Top | 0.06952 (0.06502 to 0.07403) | -Δ Net BRET |
| | | | 10 nM SR142948A | $EC_{50}$ | 9.301e-008 (4.900e-008 to 1.765e-007) | M |
| | | | | Top | Shared | -Δ Net BRET |
| | | | 100 nM SR142948A | $EC_{50}$ | 3.400e-007 (1.724e-007 to 6.706e-007) | M |
| | | | | Top | Shared | -Δ Net BRET |
| | | | 1 μM SR142948A | $EC_{50}$ | 9.267e-007 (4.894e-007 to 1.755e-006) | M |
| | | | | Top | Shared | -Δ Net BRET |
| | | | 10 μM SR142948A | $EC_{50}$ | 6.241e-006 (3.252e-006 to 1.198e-005) | M |
| | | | | Top | Shared | -Δ Net BRET |
| | | | 100 μM SR142948A | $EC_{50}$ | 4.828e-005 (1.499e-005 to 0.0001555) | M |
| | | | | Top | Shared | -Δ Net BRET |
| | Effect of SR142948A on NT-induced activation of $G_{\alpha}12$ | 3 | 0 nM SR142948A | $EC_{50}$ | 1.388e-008 (8.070e-009 to 2.389e-008) | M |
| | | | | Top | 0.3578 (0.3370 to 0.3786) | -Δ Net BRET |
| | | | 10 nM SR142948A | $EC_{50}$ | 3.305e-008 (1.868e-008 to 5.845e-008) | M |
| | | | | Top | Shared | -Δ Net BRET |
| | | | 100 nM SR142948A | $EC_{50}$ | 1.145e-007 (6.684e-008 to 1.960e-007) | M |
| | | | | Top | Shared | -Δ Net BRET |
| | | | 1 μM SR142948A | $EC_{50}$ | 5.366e-007 (3.081e-007 to 9.344e-007) | M |
| | | | | Top | Shared | -Δ Net BRET |

| | | | | | | | |
|---|---|---|---|---|---|---|---|
| | | | | 10 µM SR142948A | $EC_{50}$ | 3.542e-006 (2.011e-006 to 6.239e-006) | M |
| | | | | | Top | Shared | -∆ Net BRET |
| | | | | 100 µM SR142948A | $EC_{50}$ | 2.271e-005 (1.172e-005 to 4.400e-005) | M |
| | | | | | Top | Shared | -∆ Net BRET |
| | | Effect of SR142948A on NT-induced activation of $G_{\alpha}13$ | 3 | 0 nM SR142948A | $EC_{50}$ | 9.708e-009 (7.410e-009 to 1.272e-008) | M |
| | | | | | Top | 0.5207 (0.5068 to 0.5346) | -∆ Net BRET |
| | | | | 10 nM SR142948A | $EC_{50}$ | 2.368e-008 (1.783e-008 to 3.144e-008) | M |
| | | | | | Top | Shared | -∆ Net BRET |
| | | | | 100 nM SR142948A | $EC_{50}$ | 6.666e-008 (5.066e-008 to 8.773e-008) | M |
| | | | | | Top | Shared | -∆ Net BRET |
| | | | | 1 µM SR142948A | $EC_{50}$ | 4.201e-007 (3.163e-007 to 5.579e-007) | M |
| | | | | | Top | Shared | -∆ Net BRET |
| | | | | 10 µM SR142948A | $EC_{50}$ | 2.224e-006 (1.679e-006 to 2.946e-006) | M |
| | | | | | Top | Shared | -∆ Net BRET |
| | | | | 100 µM SR142948A | $EC_{50}$ | 8.843e-006 (6.719e-006 to 1.164e-005) | M |
| | | | | | Top | Shared | -∆ Net BRET |
| **2** | **E** | Effect of SR142948A on NT DRC $EC_{50}$ for combined G proteins, one-way ANOVA followed by Bonferroni multiple comparisons test.<br>$F (5, 78) = 6.018$, $P<0.0001$<br>*p<0.05, 100 µM vs. Vehicle | | | | | |
| **2** | **F** | Illustration | | | | | |
| **2** | **G** | Illustration | | | | | |
| **2** | **H** | Effect of SBI-553 on NT-induced β-arrestin1 recruitment | 3 | 0 µM SBI-553 | $EC_{50}$ | 8.366e-009 (5.880e-009 to 1.190e-008) | M |
| | | | | | Top | 0.5022 (0.4762 to 0.5281) | ∆ Net BRET |
| | | | | | Bottom | -0.008639 (-0.03099 to 0.01372) | ∆ Net BRET |
| | | | | 1 µM SBI-553 | $EC_{50}$ | 6.121e-009 (4.318e-009 to 8.679e-009) | M |
| | | | | | Top | 0.5590 (0.5324 to 0.5857) | ∆ Net BRET |
| | | | | | Bottom | 0.004868 (-0.01913 to 0.02886) | ∆ Net BRET |
| | | | | 3 µM SBI-553 | $EC_{50}$ | 6.668e-009 (4.723e-009 to 9.415e-009) | M |
| | | | | | Top | 0.5651 (0.5396 to 0.5906) | ∆ Net BRET |
| | | | | | Bottom | 0.03620 (0.01354 to 0.05886) | ∆ Net BRET |
| | | | | 10 µM SBI-553 | $EC_{50}$ | 7.938e-009 (4.962e-009 to 1.270e-008) | M |
| | | | | | Top | 0.5508 (0.5236 to 0.578) | ∆ Net BRET |
| | | | | | Bottom | 0.1461 (0.1225 to 0.1697) | ∆ Net BRET |
| | | | | 30 µM SBI-553 | $EC_{50}$ | 9.307e-009 (3.649e-009 to 2.374e-008) | M |
| | | | | | Top | 0.5370 (0.5061 to 0.5678) | ∆ Net BRET |
| | | | | | Bottom | 0.3108 (0.2846 to 0.3370) | ∆ Net BRET |
| | | Effect of SBI-553 on NT-induced β-arrestin2 recruitment | 3 | 0 µM SBI-553 | $EC_{50}$ | 1.462e-008 (1.119e-008 to 1.911e-008) | M |
| | | | | | Top | 0.5629 (0.5404 to 0.5853) | ∆ Net BRET |
| | | | | | Bottom | -0.001865 (-0.01979 to 0.01606) | ∆ Net BRET |
| | | | | 1 µM SBI-553 | $EC_{50}$ | 1.381e-008 (1.016e-008 to 1.878e-008) | M |

| | | | | | | |
|---|---|---|---|---|---|---|
| | | | | Top | 0.6212 (0.5937 to 0.6487) | Δ Net BRET |
| | | | | Bottom | 0.01863 (-0.003501 to 0.04076) | Δ Net BRET |
| | | | 3 μM SBI-553 | $EC_{50}$ | 1.486e-008 (1.036e-008 to 2.130e-008) | M |
| | | | | Top | 0.6242 (0.5960 to 0.6523) | Δ Net BRET |
| | | | | Bottom | 0.09722 (0.07477 to 0.1197) | Δ Net BRET |
| | | | 10 μM SBI-553 | $EC_{50}$ | 2.135e-008 (1.248e-008 to 3.651e-008) | M |
| | | | | Top | 0.6198 (0.5939 to 0.6456) | Δ Net BRET |
| | | | | Bottom | 0.2960 (0.2766 to 0.3154) | Δ Net BRET |
| | | | 30 μM SBI-553 | $EC_{50}$ | 2.337e-008 (9.356e-009 to 5.837e-008) | M |
| | | | | Top | 0.6376 (0.6188 to 0.6564) | Δ Net BRET |
| | | | | Bottom | 0.4998 (0.4859 to 0.5137) | Δ Net BRET |
| I | Effect of SBI-553 on NT-induced activation of $G_{\alpha}q$ | 3 | 0 μM SBI-553 | $EC_{50}$ | 2.792e-009 (1.935e-009 to 4.028e-009) | M |
| | | | | Top | 0.3051 (0.2929 to 0.3174) | -Δ Net BRET |
| | | | | Bottom | -0.004371 (-0.01680 to 0.008062) | -Δ Net BRET |
| | | | 1 μM SBI-553 | $EC_{50}$ | 1.784e-008 (7.137e-009 to 4.458e-008) | M |
| | | | | Top | 0.1883 (0.1634 to 0.2132) | -Δ Net BRET |
| | | | | Bottom | 0.004988 (-0.01432 to 0.02429) | -Δ Net BRET |
| | | | 3 μM SBI-553 | $EC_{50}$ | 2.444e-008 (5.071e-009 to 1.178e-007) | M |
| | | | | Top | 0.09894 (-0.001856 to 0.03925) | -Δ Net BRET |
| | | | | Bottom | -0.007306 (-0.02556 to 0.01095) | -Δ Net BRET |
| | | | 10 μM SBI-553 | $EC_{50}$ | 3.279e-008 (2.555e-010 to 4.210e-006) | M |
| | | | | Top | 0.01870 (-0.001856 to 0.03925) | -Δ Net BRET |
| | | | | Bottom | -0.009091 (-0.02332 to 0.005141) | -Δ Net BRET |
| | | | 30 μM SBI-553 | $EC_{50}$ | 5.053e-007 (5.776e-018 to 44201) | M |
| | | | | Top | 0.004370 (-0.01697 to 0.02571) | -Δ Net BRET |
| | | | | Bottom | 0.0004696 (-0.007560 to 0.008499) | -Δ Net BRET |
| | Effect of SBI-553 on NT-induced activation of $G_{\alpha}11$ | 3 | 0 μM SBI-553 | $EC_{50}$ | 3.669e-009 (2.549e-009 to 5.281e-009) | M |
| | | | | Top | 0.2685 (0.2561 to 0.2809) | -Δ Net BRET |
| | | | | Bottom | 0.001133 (-0.01097 to 0.01323) | -Δ Net BRET |
| | | | 1 μM SBI-553 | $EC_{50}$ | 9.765e-009 (5.738e-009 to 1.662e-008) | M |
| | | | | Top | 0.2327 (0.2136 to 0.2517) | -Δ Net BRET |
| | | | | Bottom | -0.01223 (-0.02830 to 0.003844) | -Δ Net BRET |
| | | | 3 μM SBI-553 | $EC_{50}$ | 2.030e-008 (1.156e-008 to 3.566e-008) | M |
| | | | | Top | 0.2067 (0.1886 to 0.2247) | -Δ Net BRET |
| | | | | Bottom | -0.009058 (-0.02274 to 0.004621) | -Δ Net BRET |
| | | | 10 μM SBI-553 | $EC_{50}$ | 2.774e-008 (1.236e-008 to 6.229e-008) | M |
| | | | | Top | 0.1197 (0.1038 to 0.1355) | -Δ Net BRET |
| | | | | Bottom | -0.01084 (-0.02217 to 0.0004920) | -Δ Net BRET |
| | | | 30 μM SBI-553 | $EC_{50}$ | 2.705e-008 (2.445e-009 to 2.991e-007) | M |
| | | | | Top | 0.03650 (0.01804 to 0.05496) | -Δ Net BRET |

| | | | | | | |
|---|---|---|---|---|---|---|
| | | | | Bottom | -0.01476 (-0.02803 to -0.001487) | -Δ Net BRET |
| | Effect of SBI-553 on NT-induced activation of $G_\alpha 15$ | 3 | 0 µM SBI-553 | $EC_{50}$ | 6.933e-009 (3.610e-009 to 1.331e-008) | M |
| | | | | Top | 0.1293 (0.1185 to 0.1400) | -Δ Net BRET |
| | | | | Bottom | -0.002762 (-0.01209 to 0.006568) | -Δ Net BRET |
| | | | 1 µM SBI-553 | $EC_{50}$ | 1.307e-008 (1.554e-009 to 1.100e-007) | M |
| | | | | Top | 0.08373 (0.06876 to 0.09870) | -Δ Net BRET |
| | | | | Bottom | 0.03628 (0.02413 to 0.04844) | -Δ Net BRET |
| | | | 3 µM SBI-553 | $EC_{50}$ | 2.227e-009 (8.828e-011 to 5.616e-008) | M |
| | | | | Top | 0.08597 (0.07606 to 0.09588( | -Δ Net BRET |
| | | | | Bottom | 0.06113 (0.05062 to 0.07165) | -Δ Net BRET |
| | | | 10 µM SBI-553 | $EC_{50}$ | 5.569e-008 (2.899e-009 to 1.070e-006) | M |
| | | | | Top | 0.07667 (0.06718 to 0.08615) | -Δ Net BRET |
| | | | | Bottom | 0.09601 (0.09005 to 0.1020) | -Δ Net BRET |
| | | | 30 µM SBI-553 | $EC_{50}$ | 4.181e-008 (1.701e-008 to 1.028e-007) | M |
| | | | | Top | 0.05093 (0.04292 to 0.05895) | -Δ Net BRET |
| | | | | Bottom | 0.1075 (0.1022 to 0.1128) | -Δ Net BRET |
| | Effect of SBI-553 on NT-induced activation of $G_\alpha i1$ | 3 | 0 µM SBI-553 | $EC_{50}$ | 4.989e-009 (2.837e-009 to 8.772e-009) | M |
| | | | | Top | 0.2681 (0.2486 to 0.2876) | -Δ Net BRET |
| | | | | Bottom | 0.009071 (-0.009004 to 0.02715) | -Δ Net BRET |
| | | | 1 µM SBI-553 | $EC_{50}$ | 6.738e-009 (4.318e-009 to 1.052e-008) | M |
| | | | | Top | 0.2282 (0.2139 to 0.2426) | -Δ Net BRET |
| | | | | Bottom | -0.001507 (-0.01421 to 0.01119) | -Δ Net BRET |
| | | | 3 µM SBI-553 | $EC_{50}$ | 6.691e-009 (4.546e-009 to 9.848e-009) | M |
| | | | | Top | 0.2242 (0.2120 to 0.2364) | -Δ Net BRET |
| | | | | Bottom | -0.0009451 (-0.01175 to 0.009864) | -Δ Net BRET |
| | | | 10 µM SBI-553 | $EC_{50}$ | 5.301e-009 (3.713e-009 to 7.568e-009) | M |
| | | | | Top | 0.2205 (0.2106 to 0.2305) | -Δ Net BRET |
| | | | | Bottom | 0.01236 (0.003189 to 0.02153) | -Δ Net BRET |
| | | | 30 µM SBI-553 | $EC_{50}$ | 7.585e-009 (4.907e-009 to 1.172e-008) | M |
| | | | | Top | 0.2159 (0.2055 to 0.2264) | -Δ Net BRET |
| | | | | Bottom | 0.04686 (0.03771 to 0.05601) | -Δ Net BRET |
| | Effect of SBI-553 on NT-induced activation of $G_\alpha i2$ | 3 | 0 µM SBI-553 | $EC_{50}$ | 8.190e-009 (4.390e-009 to 1.528e-008) | M |
| | | | | Top | 0.2543 (0.2317 to 0.2769) | -Δ Net BRET |
| | | | | Bottom | 0.002192 (-0.01732 to 0.02145) | -Δ Net BRET |
| | | | 1 µM SBI-553 | $EC_{50}$ | 1.932e-008 (9.379e-009 to 3.982e-008) | M |
| | | | | Top | 0.2191 (0.1952 to 0.2429) | -Δ Net BRET |
| | | | | Bottom | -0.002859 (-0.02090 to 0.01497) | -Δ Net BRET |
| | | | 3 µM SBI-553 | $EC_{50}$ | 1.958e-008 (9.322e-009 to 4.113e-008) | M |
| | | | | Top | 0.2284 (0.2018 to 0.2550) | -Δ Net BRET |

| | | | | | | |
|---|---|---|---|---|---|---|
| | | | | Bottom | -0.01298 (-0.03320 to 0.006989) | -Δ Net BRET |
| | | | 10 μM SBI-553 | $EC_{50}$ | 1.756e-008 (8.656e-009 to 3.563e-008) | M |
| | | | | Top | 0.2167 (0.1928 to 0.2406) | -Δ Net BRET |
| | | | | Bottom | -0.01105 (-0.02953 to 0.007208) | -Δ Net BRET |
| | | | 30 μM SBI-553 | $EC_{50}$ | 1.437e-008 (6.956e-009 to 3.018e-008) | M |
| | | | | Top | 0.1946 (0.1770 to 0.2129) | -Δ Net BRET |
| | | | | Bottom | 0.03008 (0.01562 to 0.04435) | -Δ Net BRET |
| | Effect of SBI-553 on NT-induced activation of $G_{\alpha}i3$ | 3 | 0 μM SBI-553 | $EC_{50}$ | 2.789e-008 (2.029e-008 to 3.836e-008) | M |
| | | | | Top | 0.4199 (0.3996 to 0.4402) | -Δ Net BRET |
| | | | | Bottom | -0.004657 (-0.01917 to 0.009854) | -Δ Net BRET |
| | | | 1 μM SBI-553 | $EC_{50}$ | 2.659e-008 (1.699e-008 to 4.159e-008) | M |
| | | | | Top | 0.2034 (0.1890 to 0.2177) | -Δ Net BRET |
| | | | | Bottom | -0.01096 (-0.02132 to -0.0006084) | -Δ Net BRET |
| | | | 3 μM SBI-553 | $EC_{50}$ | 2.785e-008 (1.605e-008 to 4.832e-008) | M |
| | | | | Top | 0.1906 (0.1742 to 0.2070) | -Δ Net BRET |
| | | | | Bottom | -0.008003 (-0.01975 to 0.003740) | -Δ Net BRET |
| | | | 10 μM SBI-553 | $EC_{50}$ | 2.098e-008 (1.148e-008 to 3.832e-008) | M |
| | | | | Top | 0.1827 (0.1664 to 0.1991) | -Δ Net BRET |
| | | | | Bottom | 6.552e-005 (-0.01226 to 0.01239) | -Δ Net BRET |
| | | | 30 μM SBI-553 | $EC_{50}$ | 2.105e-008 (9.406e-009 to 4.712e-008) | M |
| | | | | Top | 0.1966 (0.1776 to 0.2156) | -Δ Net BRET |
| | | | | Bottom | 0.03771 (0.02339 to 0.05203) | -Δ Net BRET |
| | Effect of SBI-553 on NT-induced activation of $G_{\alpha}oA$ | 3 | 0 μM SBI-553 | $EC_{50}$ | 1.315e-008 (9.775e-009 to 1.769e-008) | M |
| | | | | Top | 0.2966 (0.2833 to 0.3099) | -Δ Net BRET |
| | | | | Bottom | -0.006282 (-0.01708 to 0.004518) | -Δ Net BRET |
| | | | 1 μM SBI-553 | $EC_{50}$ | 1.450e-008 (1.091e-008 to 1.927e-008) | M |
| | | | | Top | 0.2735 (0.2622 to 0.2849) | -Δ Net BRET |
| | | | | Bottom | 0.004023 (-0.005073 to 0.01312) | -Δ Net BRET |
| | | | 3 μM SBI-553 | $EC_{50}$ | 1.252e-008 (9.185e-009 to 1.706e-008) | M |
| | | | | Top | 0.2826 (0.2704 to 0.2947) | -Δ Net BRET |
| | | | | Bottom | 0.01642 (0.006461 to 0.02638) | -Δ Net BRET |
| | | | 10 μM SBI-553 | $EC_{50}$ | 1.468e-008 (9.237e-009 to 2.332e-008) | M |
| | | | | Top | 0.2868 (0.2716 to 0.3021) | -Δ Net BRET |
| | | | | Bottom | 0.06481 (0.05263 to 0.07700) | -Δ Net BRET |
| | | | 30 μM SBI-553 | $EC_{50}$ | 1.034e-008 (6.084e-009 to 1.758e-008) | M |
| | | | | Top | 0.2786 (0.2669 to 0.2904) | -Δ Net BRET |
| | | | | Bottom | 0.1174 (0.1074 to 0.1273) | -Δ Net BRET |
| | Effect of SBI-553 on NT- | 3 | 0 μM SBI-553 | $EC_{50}$ | 3.583e-008 (2.257e-008 to 5.687e-008) | M |
| | | | | Top | 0.2827 (0.2629 to 0.3025) | -Δ Net BRET |

| | | | | | | |
|---|---|---|---|---|---|---|
| | induced activation of $G_{\alpha}oB$ | | | Bottom | 0.004527 (-0.008947 to 0.01800) | -Δ Net BRET |
| | | | 1 μM SBI-553 | $EC_{50}$ | 3.914e-008 (2.389e-008 to 6.412e-008) | M |
| | | | | Top | 0.2768 (0.2560 to 0.2976) | -Δ Net BRET |
| | | | | Bottom | 0.006857 (-0.007049 to 0.02076) | -Δ Net BRET |
| | | | 3 μM SBI-553 | $EC_{50}$ | 2.777e-008 (1.894e-008 to 4.072e-008) | M |
| | | | | Top | 0.2794 (0.2642 to 0.2947) | -Δ Net BRET |
| | | | | Bottom | 0.01430 (0.003404 to 0.02520) | -Δ Net BRET |
| | | | 10 μM SBI-553 | $EC_{50}$ | 2.555e-008 (1.598e-008 to 4.084e-008) | M |
| | | | | Top | 0.2828 (0.2658 to 0.2998) | -Δ Net BRET |
| | | | | Bottom | 0.04076 (0.02844 to 0.05309) | -Δ Net BRET |
| | | | 30 μM SBI-553 | $EC_{50}$ | 1.785e-008 (9.400e-009 to 3.389e-008) | M |
| | | | | Top | 0.2718 (0.2564 to 0.2872) | -Δ Net BRET |
| | | | | Bottom | 0.1009 (0.08906 to 0.1128) | -Δ Net BRET |
| | Effect of SBI-553 on NT-induced activation of $G_{\alpha}z$ | 3 | 0 μM SBI-553 | $EC_{50}$ | 6.727e-009 (3.885e-009 to 1.165e-008) | M |
| | | | | Top | 0.2871 (0.2658 to 0.3085) | -Δ Net BRET |
| | | | | Bottom | 0.009686 (-0.009238 to 0.02861) | -Δ Net BRET |
| | | | 1 μM SBI-553 | $EC_{50}$ | 1.467e-008 (7.816e-009 to 2.752e-008) | M |
| | | | | Top | 0.1972 (0.1798 to 0.2147) | -Δ Net BRET |
| | | | | Bottom | 0.01057 (-0.003348 to 0.02450) | -Δ Net BRET |
| | | | 3 μM SBI-553 | $EC_{50}$ | 1.795e-008 (7.919e-009 to 4.068e-008) | M |
| | | | | Top | 0.1527 (0.1358 to 0.1696) | -Δ Net BRET |
| | | | | Bottom | 0.01348 (0.0004051 to 0.02656) | -Δ Net BRET |
| | | | 10 μM SBI-553 | $EC_{50}$ | 1.341e-008 (5.128e-009 to 3.508e-008) | M |
| | | | | Top | 0.1145 (0.09960 to 0.1295) | -Δ Net BRET |
| | | | | Bottom | 0.009683 (-0.002410 to 0.02177) | -Δ Net BRET |
| | | | 30 μM SBI-553 | $EC_{50}$ | 1.117e-008 (3.360e-009 to 3.716e-008) | M |
| | | | | Top | 0.08938 (0.07731 to 0.1014) | -Δ Net BRET |
| | | | | Bottom | 0.02120 (0.01119 to 0.03122) | -Δ Net BRET |
| | Effect of SBI-553 on NT-induced activation of $G_{\alpha}$gustducin | 4 | 0 μM SBI-553 | $EC_{50}$ | 1.265e-008 (4.984e-009 to 3.212e-008) | M |
| | | | | Top | 0.07106 (0.06256 to 0.07956) | -Δ Net BRET |
| | | | | Bottom | 0.009434 (0.002500 to 0.01637) | -Δ Net BRET |
| | | | 1 μM SBI-553 | $EC_{50}$ | 4.670e-008 (5.463e-009 to 3.992e-007) | M |
| | | | | Top | 0.05056 (0.03826 to 0.06286) | -Δ Net BRET |
| | | | | Bottom | 0.01486 (0.006894 to 0.02283) | -Δ Net BRET |
| | | | 3 μM SBI-553 | $EC_{50}$ | 2.364e-008 (4.773e-009 to 1.171e-007) | M |
| | | | | Top | 0.05660 (0.04559 to 0.06761) | -Δ Net BRET |
| | | | | Bottom | 0.01039 (0.002273 to 0.01851) | -Δ Net BRET |
| | | | 10 μM SBI-553 | $EC_{50}$ | 4.065e-008 (7.725e-009 to 2.139e-007) | M |
| | | | | Top | 0.06659 (0.05620 to 0.07699) | -Δ Net BRET |

| | | | | | | | |
|---|---|---|---|---|---|---|---|
| | | | | | Bottom | 0.02669 (0.01978 to 0.03359) | -Δ Net BRET |
| | | | | 30 μM SBI-553 | $EC_{50}$ | 1.090e-008 (2.185e-009 to 5.434e-008) | M |
| | | | | | Top | 0.07036 (0.06389 to 0.07684) | -Δ Net BRET |
| | | | | | Bottom | 0.04299 (0.03760 to 0.04838) | -Δ Net BRET |
| | | Effect of SBI-553 on NT-induced activation of $G_{\alpha}12$ | 3 | 0 μM SBI-553 | $EC_{50}$ | 1.885e-008 (1.036e-008 to 3.429e-008) | M |
| | | | | | Top | 0.3125 (0.2855 to 0.3395) | -Δ Net BRET |
| | | | | | Bottom | 0.008787 (-0.01191 to 0.02949) | -Δ Net BRET |
| | | | | 1 μM SBI-553 | $EC_{50}$ | 7.868e-009 (4.457e-009 to 1.389e-008) | M |
| | | | | | Top | 0.3103 (0.2864 to 0.3341) | -Δ Net BRET |
| | | | | | Bottom | 0.01697 (-0.003742 to 0.03768) | -Δ Net BRET |
| | | | | 3 μM SBI-553 | $EC_{50}$ | 5.083e-009 (2.636e-009 to 9.805e-009) | M |
| | | | | | Top | 0.3101 (0.2869 to 0.3334) | -Δ Net BRET |
| | | | | | Bottom | 0.04556 (0.02407 to 0.06706) | -Δ Net BRET |
| | | | | 10 μM SBI-553 | $EC_{50}$ | 7.104e-009 (2.718e-009 to 1.856e-008) | M |
| | | | | | Top | 0.3113 (0.2817 to 0.3409) | -Δ Net BRET |
| | | | | | Bottom | 0.09278 (0.06669 to 0.1189) | -Δ Net BRET |
| | | | | 30 μM SBI-553 | $EC_{50}$ | 7.633e-009 (1.801e-009 to 3.236e-008) | M |
| | | | | | Top | 0.3011 (0.2741 to 0.3280) | -Δ Net BRET |
| | | | | | Bottom | 0.1700 (0.1465 to 0.1935) | -Δ Net BRET |
| | | Effect of SBI-553 on NT-induced activation of $G_{\alpha}13$ | 3 | 0 μM SBI-553 | $EC_{50}$ | 1.295e-008 (8.876e-009 to 1.888e-008) | M |
| | | | | | Top | 0.5175 (0.4884 to 0.5467) | -Δ Net BRET |
| | | | | | Bottom | -0.004046 (-0.02776 to 0.01966) | -Δ Net BRET |
| | | | | 1 μM SBI-553 | $EC_{50}$ | 1.009e-008 (6.561e-009 to 1.552e-008) | M |
| | | | | | Top | 0.4853 (0.4554 to 0.5152) | -Δ Net BRET |
| | | | | | Bottom | 0.01054 (-0.01463 to 0.03571) | -Δ Net BRET |
| | | | | 3 μM SBI-553 | $EC_{50}$ | 8.475e-009 (5.894e-009 to 1.218e-008) | M |
| | | | | | Top | 0.4924 (0.4687 to 0.5162) | -Δ Net BRET |
| | | | | | Bottom | 0.03886 (0.01842 to 0.05930) | -Δ Net BRET |
| | | | | 10 μM SBI-553 | $EC_{50}$ | 6.319e-009 (4.291e-009 to 9.304e-009) | M |
| | | | | | Top | 0.4896 (0.4674 to 0.5118) | -Δ Net BRET |
| | | | | | Bottom | 0.07522 (0.05532 to 0.09512) | -Δ Net BRET |
| | | | | 30 μM SBI-553 | $EC_{50}$ | 3.436e-009 (1.730e-009 to 6.825e-009) | M |
| | | | | | Top | 0.4703 (0.4448 to 0.4958) | -Δ Net BRET |
| | | | | | Bottom | 0.1763 (0.1512 to 0.2015) | -Δ Net BRET |
| **2** | **J** | Effect of SBI-553 on NT DRC $EC_{50}$ for combined G proteins, one-way ANOVA followed by Bonferroni multiple comparisons test.<br><br>F (2.357, 22.39) = 0.9654, P = 0.4088<br><br>p<0.5 (n.s.), 30 μM vs. Vehicle | | | | | |
| **2** | **K** | Effect of SBI-553 on NT potency. *indicates G proteins for which the NT $EC_{50}$ vs. NT + SBI-553 $EC_{50}$ have non-overlapping 95% confidence intervals (CI). SBI-553 dose selected was the highest for which a sigmoidal curve could be fit. | | | | | |
| | | | | NT | | NT + SBI-553 | |

| | Mean (M) | 95% CI (M) | Mean (M) | 95% CI (M) | Fold-change |
|---|---|---|---|---|---|
| G13 | 1.3E-08 | 8.876e-09 to 1.888e-08 | 3.436E-09 | 1.730e-09 to 6.825e-09 | 3.768917346 |
| G12 | 1.89E-08 | 1.036e-08 to 3.429e-08 | 7.63E-09 | 1.801e-09 to 3.236e-08 | 2.469540155 |
| Gz | 6.73E-09 | 3.885e-09 to 1.165e-08 | 1.12E-08 | 3.360e-09 to 3.716e-08 | 0.602238138 |
| Gg | 1.27E-08 | 4.984e-09 to 3.212e-08 | 1.09E-08 | 2.185e-09 to 5.434e-08 | 1.160550459 |
| GoB | 3.58E-08 | 2.257e-08 to 5.687e-08 | 2.371E-08 | 1.255e-08 to 4.479e-08 | 1.511176719 |
| GoA | 1.32E-08 | 9.775e-09 to 1.769e-08 | 1.385E-08 | 8.069e-09 to 2.377e-08 | 0.949458484 |
| Gi3 | 2.79E-08 | 2.029e-08 to 3.836e-08 | 2.105E-08 | 9.405e-09 to 4.711e-08 | 1.324940618 |
| Gi2 | 8.19E-09 | 4.459e-09 to 1.480e-08 | 1.437E-08 | 6.956e-09 to 3.018e-08 | 0.56993737 |
| Gi1 | 4.99E-09 | 2.837e-09 to 8.772e-09 | 7.585E-09 | 4.907e-09 to 1.172e-08 | 0.65774555 |
| G15 | 6.93E-09 | 3.610e-09 to 1.331e-08 | 4.181E-08 | 1.701e-08 to 1.028e-07 | 0.165821574 |
| G11 | 3.67E-09 | 2.549e-09 to 5.281e-09 | 2.705E-08 | 1.236e-08 to 6.229e-08 | 0.135637708 |
| Gq | 2.79E-09 | 1.935e-09 to 4.028e-09 | 3.279E-08 | 2.555e-08 to 4.210e-06 | 0.085147911 |

| | | | | | | | |
|---|---|---|---|---|---|---|---|
| **2** | **L** | Radar plots summarize the extent of activation of each transducer in the presence NT, SBI-553, and both NT and SBI-553 | | | | | |
| **2** | **M** | Effect of SR142948A vs. SBI-553 on NT-induced G protein activation by TGFα Shedding - $G_{\alpha}q$ | 4 | SR142948A | $IC_{50}$ | 1.757e-007 (1.478e-007 to 2.084e-007) | M |
| | | | | | Bottom | -0.3333 (-1.539 to 0.8547) | % AP Shedding |
| | | | | SBI-553 | $IC_{50}$ | 4.135e-007 (ND) | M |
| | | | | | Bottom | 2.081 (0.1138 to 3.679) | % AP Shedding |
| | | Effect of SR142948A vs. SBI-553 on NT-induced G protein activation by TGFα Shedding - $G_{\alpha}i1/2$ | 3 | SR142948A | $IC_{50}$ | 1.843e-008 (1.244e-008 to 2.712e-008) | M |
| | | | | | Bottom | 0.2383 (-0.7989 to 1.103) | % AP Shedding |
| | | | | SBI-553 | $IC_{50}$ | 4.411e-008 (5.210e-009 to 1.033e-006) | M |
| | | | | | Bottom | 14.53 (12.66 to 16.75) | % AP Shedding |
| | | Effect of SR142948A vs. SBI-553 on NT-induced G protein activation by TGFα Shedding - $G_{\alpha}o$ | 3 | SR142948A | $IC_{50}$ | 1.708e-007 (1.465e-007 to 1.981e-007) | M |
| | | | | | Bottom | -1.409 (-2.142 to -0.6838) | % AP Shedding |
| | | | | SBI-553 | $IC_{50}$ | 1.426e-007 (8.519e-008 to 2.893e-007) | M |
| | | | | | Bottom | 10.95 (9.954 to 11.79) | % AP Shedding |
| | | Effect of SR142948A vs. SBI-553 on NT-induced G protein activation by TGFα Shedding - $G_{\alpha}12$ | 3 | SR142948A | $IC_{50}$ | 1.175e-008 (7.754e-009 to 1.737e-008) | M |
| | | | | | Bottom | 1.058 (0.2823 to 1.806) | % AP Shedding |
| | | | | SBI-553 | $IC_{50}$ | 3.773e-009 (1.102e-012 to 1.292e-005) | M |
| | | | | | Bottom | 13.80 (8.486 to 19.10) | % AP Shedding |
| **2** | **N** | Illustration | | | | | |

CI, confidence interval; NT, neurotensin; SBI-553, SBI-0654553; $EC_{50}$, half-maximal effective concentration; ND, not defined

**Table S3. Curve parameters and statistical comparisons presented in Figure 3 – Supporting Figure 3.**

| Fig | Panel | Description | N | Condition | Parameter | Mean (95% CI) | Units |
|---|---|---|---|---|---|---|---|
| **3** | **A** | Illustration | | | | | |
| **3** | **B** | Effect of SR142948A vs. SBI-553 on NT-induced recruitment of mini $G_{\alpha}q$ | 3 | SBI-553 | $IC_{50}$ | 1.944e-008 (1.689e-008 to 2.237e-008) | M |
| | | | | | Top | 0.01585 (0.01360 to 0.01808) | Δ Net BRET |
| | | | | | Bottom | 0.1404 (0.1368 to 0.1440) | Δ Net BRET |
| | | | | SR142948A | $IC_{50}$ | 3.786e-008 (2.734e-008 to 5.199e-008) | M |
| | | | | | Top | 0.005873 (0.0007201 to 0.01097) | Δ Net BRET |
| | | | | | Bottom | 0.1410 (0.1361 to 0.1460) | Δ Net BRET |
| | | Effect of SR142948A vs. SBI-553 on NT-induced recruitment of mini $G_{\alpha}i1$ | 3 | SBI-553 | $IC_{50}$ | 6.170e-009 (4.935e-009 to 7.711e-009) | M |
| | | | | | Top | 0.04704 (0.04563 to 0.04844) | Δ Net BRET |
| | | | | | Bottom | 0.1037 (0.1009 to 0.1066) | Δ Net BRET |
| | | | | SR142948A | $IC_{50}$ | 1.492e-008 (1.116e-008 to 2.012e-008) | M |
| | | | | | Top | 0.003487 (0.0002021 to 0.006742) | Δ Net BRET |
| | | | | | Bottom | 0.09830 (0.09463 to 0.1020) | Δ Net BRET |
| | | Effect of SR142948A vs. SBI-553 on NT-induced recruitment of mini $G_{\alpha}s$ | 3 | SBI-553 | $IC_{50}$ | 5.494e-009 (2.383e-009 to 1.368e-008) | M |
| | | | | | Top | 0.01117 (0.009764 to 0.01250) | Δ Net BRET |
| | | | | | Bottom | 0.02551 (0.02261 to 0.02849) | Δ Net BRET |
| | | | | SR142948A | $IC_{50}$ | 1.055e-008 (3.626e-009 to 3.216e-008) | M |
| | | | | | Top | 0.004397 (0.001924 to 0.006809) | Δ Net BRET |
| | | | | | Bottom | 0.02416 (0.02132 to 0.02710) | Δ Net BRET |
| | | Effect of SR142948A vs. SBI-553 on NT-induced recruitment of mini $G_{\alpha}o$ | 3 | SBI-553 | $IC_{50}$ | 1.019e-008 (3.892e-009 to 2.522e-008) | M |
| | | | | | Top | 0.02176 (0.01974 to 0.02345) | Δ Net BRET |
| | | | | | Bottom | 0.02583 (0.02090 to 0.02676) | Δ Net BRET |
| | | | | SR142948A | $IC_{50}$ | 3.694e-009 (ND) | M |
| | | | | | Top | 0.003910 (0.001731 to 0.006042) | Δ Net BRET |
| | | | | | Bottom | 0.02507 (0.02299 to 0.02719) | Δ Net BRET |
| | | Effect of SR142948A vs. SBI-553 on NT-induced recruitment of mini $G_{\alpha}12$ | 3 | SBI-553 | $IC_{50}$ | 2.265e-009 (5.540e-010 to 6.552e-009) | M |
| | | | | | Top | 0.02865 (0.02686 to 0.03050) | Δ Net BRET |
| | | | | | Bottom | 0.01426 (0.01052 to 0.01796) | Δ Net BRET |
| | | | | SR142948A | $IC_{50}$ | 1.996e-008 (2.533e-009 to 1.869e-007) | M |
| | | | | | Top | 0.005963 (0.003774 to 0.007751) | Δ Net BRET |
| | | | | | Bottom | 0.01335 (0.01171 to 0.01508) | Δ Net BRET |
| **3** | **C** | Effect of SBI-553 on NT-induced recruitment of mini $G_{\alpha}q$ in β-arrestin1/2-null | 3 | Parental HEK293 cells | $IC_{50}$ | 2.286e-008 (1.815e-008 to 2.904e-008) | M |
| | | | | | Top | 0.09135 (0.08708 to 0.09572) | Δ Net BRET |
| | | | | | Bottom | -0.02950 (-0.03419 to –0.02557) | Δ Net BRET |

| | | | | | |
|---|---|---|---|---|---|
| vs. Parental HEK293 cells | 3 | β-arrestin1/2-null HEK293 cells | $IC_{50}$ | 1.022e-008 (7.769e-009 to 1.351e-008) | M |
| | | | Top | 0.1194 (0.1128 to 0.1261) | ∆ Net BRET |
| | | | Bottom | -0.02246 (-0.02846 to –0.01751) | ∆ Net BRET |

CI, confidence interval; NT, neurotensin; SBI-553, SBI-0654553; $EC_{50}$, half-maximal effective concentration; ND, not defined

**Table S4. Curve parameters and statistical comparisons presented in Figure 4 – Supporting Figure 4.**

| Fig | Panel | Description | N | Condition | Parameter | Mean (95% CI) | Units |
|---|---|---|---|---|---|---|---|
| 4 | A | Illustration | | | | | |
| 4 | B | Effect of SBI-553 on NT-induced activation of $G_\alpha oA^{G\alpha q\ C\text{-}term}$ | 3 | Vehicle | $EC_{50}$ | 2.001e-009 (1.278e-009 to 3.133e-009) | M |
| | | | | | Top | 0.3083 (0.2909 to 0.3256) | -Δ Net BRET |
| | | | | | Bottom | -0.004874 (-0.02360 to 0.01385) | -Δ Net BRET |
| | | | | 1 µM SBI-553 | $EC_{50}$ | 9.427e-009 (6.001e-009 to 1.481e-008) | M |
| | | | | | Top | 0.2294 (0.2132 to 0.2456) | -Δ Net BRET |
| | | | | | Bottom | -0.01694 (-0.03069 to -0.003181) | -Δ Net BRET |
| | | | | 3 µM SBI-553 | $EC_{50}$ | 1.328e-008 (9.228e-009 to 1.911e-008) | M |
| | | | | | Top | 0.1828 (0.1719 to 0.1937) | -Δ Net BRET |
| | | | | | Bottom | -0.01915 (-0.02798 to -0.01033) | -Δ Net BRET |
| | | | | 10 µM SBI-553 | $EC_{50}$ | 1.611e-008 (8.681e-009 to 2.990e-008) | M |
| | | | | | Top | 0.1466 (0.1316 to 0.1617) | -Δ Net BRET |
| | | | | | Bottom | -0.01726 (-0.02910 to -0.005414) | -Δ Net BRET |
| | | | | 30 µM SBI-553 | $EC_{50}$ | 2.783e-008 (1.528e-008 to 5.070e-008) | M |
| | | | | | Top | 0.1194 (0.1086 to 0.1301) | -Δ Net BRET |
| | | | | | Bottom | 0.0001591 (-0.007512 to 0.007831) | -Δ Net BRET |
| 4 | C | Effect of SBI-553 on NT-induced activation of $G_\alpha q^{G\alpha oA\ C\text{-}term}$ | 3 | Vehicle | $EC_{50}$ | 8.359e-9 (5.446e-9 to 1.283e-8) | M |
| | | | | | Top | 0.2349 (0.2198 to 0.2501) | -Δ Net BRET |
| | | | | | Bottom | -0.01111 (-0.02419 to 0.001977) | -Δ Net BRET |
| | | | | 1 µM SBI-553 | $EC_{50}$ | 5.962e-9 (3.824e-9 to 9.296e-9) | M |
| | | | | | Top | 0.1630 (0.1522 to 0.1739) | -Δ Net BRET |
| | | | | | Bottom | -0.01512 (-0.02493 to -0.005305) | -Δ Net BRET |
| | | | | 3 µM SBI-553 | $EC_{50}$ | 4.269e-9 (2.941e-9 to 6.198e-9) | M |
| | | | | | Top | 0.1635 (0.1551 to 0.1719) | -Δ Net BRET |
| | | | | | Bottom | -0.009743 (-0.01773 to -0.001759) | -Δ Net BRET |
| | | | | 10 µM SBI-553 | $EC_{50}$ | 4.339e-9 (2.651e-9 to 7.103e-9) | M |
| | | | | | Top | 0.1533 (0.1436 to 0.1631) | -Δ Net BRET |
| | | | | | Bottom | 0.001367 (-0.007892 to 0.01063) | -Δ Net BRET |
| | | | | 30 µM SBI-553 | $EC_{50}$ | 7.361e-9 (4.406e-9 to 1.230e-8) | M |
| | | | | | Top | 0.1268 (0.1180 to 0.1355) | -Δ Net BRET |
| | | | | | Bottom | 0.007651 (0.001782 to 0.01352) | -Δ Net BRET |
| 4 | D | Illustration | | | | | |

| | | | | | | | |
|---|---|---|---|---|---|---|---|
| **4** | **E** | Effect of SBI-553 on NT-induced activation of $G_{\alpha}q^{\Delta 234\text{-}246\ G\alpha oA}$ | 9 | Vehicle | $EC_{50}$ | 1.779e-008 (4.386e-009 to 7.215e-008) | M |
| | | | | | Top | 0.05623 (0.04437 to 0.06809) | -Δ Net BRET |
| | | | | | Bottom | -0.0008487 (-0.01004 to 0.008345) | -Δ Net BRET |
| | | | | 1 µM SBI-553 | $EC_{50}$ | 5.821e-009 (3.054e-009 to 1.109e-008) | M |
| | | | | | Top | 0.06294 (0.05567 to 0.07021) | -Δ Net BRET |
| | | | | | Bottom | -0.01949 (-0.02607 to -0.01290) | -Δ Net BRET |
| | | | | 3 µM SBI-553 | $EC_{50}$ | 5.542e-009 (2.872e-009 to 1.070e-008) | M |
| | | | | | Top | 0.06090 (0.05348 to 0.06832) | -Δ Net BRET |
| | | | | | Bottom | -0.02225 (-0.02902 to -0.01548) | -Δ Net BRET |
| | | | | 10 µM SBI-553 | $EC_{50}$ | 4.049e-009 (1.760e-009 to 9.313e-009) | M |
| | | | | | Top | 0.04991 (0.04303 to 0.05678) | -Δ Net BRET |
| | | | | | Bottom | -0.01402 (-0.02061 to -0.007426) | -Δ Net BRET |
| | | | | 30 µM SBI-553 | $EC_{50}$ | 5.451e-009 (1.028e-009 to 2.891e-008) | M |
| | | | | | Top | 0.02168 (0.01402 to 0.02934) | -Δ Net BRET |
| | | | | | Bottom | -0.01224 (-0.01925 to -0.005238) | -Δ Net BRET |
| **4** | **F** | G protein C-terminus amino acid alignment | | | | | |
| **4** | **G** | Comparison of amino acid properties at positions that differ between Gq and other G proteins | | | | | |
| **4** | **H** | NT-induced activation of Gq point mutants | 3-4 | $G_{\alpha}q^{WT}$ | $EC_{50}$ | 4.078e-009 (2.678e-009 to 6.210e-009) | M |
| | | | | | Top | 0.2992 (0.2823 to 0.3161) | -Δ Net BRET |
| | | | | | Bottom | -0.01188 (-0.02807 to 0.004317) | -Δ Net BRET |
| | | | | $G_{\alpha}q^{V246Y}$ | $EC_{50}$ | 1.937e-008 (1.251e-008 to 3.000e-008) | M |
| | | | | | Top | 0.2717 (0.2536 to 0.2899) | -Δ Net BRET |
| | | | | | Bottom | -0.007551 (-0.02141 to 0.006312) | -Δ Net BRET |
| | | | | $G_{\alpha}q^{N244G}$ | $EC_{50}$ | 5.260e-009 (3.847e-009 to 7.192e-009) | M |
| | | | | | Top | 0.2809 (0.2691 to 0.2927) | -Δ Net BRET |
| | | | | | Bottom | 0.001289 (-0.009536 to 0.01211) | -Δ Net BRET |
| | | | | $G_{\alpha}q^{Y243C}$ | $EC_{50}$ | 5.641e-009 (3.759e-009 to 8.466e-009) | M |
| | | | | | Top | 0.2971 (0.2807 to 0.3135) | -Δ Net BRET |
| | | | | | Bottom | 0.0001203 (-0.01482 to 0.01506) | -Δ Net BRET |
| | | | | $G_{\alpha}q^{L238N}$ | $EC_{50}$ | 6.996e-009 (2.083e-009 to 2.350e-008) | M |
| | | | | | Top | 0.06447 (0.05225 to 0.07670) | -Δ Net BRET |
| | | | | | Bottom | -0.007251 (-0.01805 to 0.003547) | -Δ Net BRET |
| **4** | **I** | Effect of SBI-553 on NT-induced | 3-7 | $G_{\alpha}q^{WT}$ | $IC_{50}$ | 1.603e-006 (1.153e-006 to 2.229e-006) | M |
| | | | | | Top | 102.1 (97.48 to 106.7) | % NT Activation |

| | | | | | | |
|---|---|---|---|---|---|---|
| | | activation of Gq point mutants | | Bottom | 3.250 (-3.808 to 10.31) | % NT Activation |
| | | | $G_{\alpha}q^{V246Y}$ | $IC_{50}$ | 6.488e-007 (4.873e-007 to 8.638e-007) | M |
| | | | | Top | 103.9 (89.37 to 118.5) | % NT Activation |
| | | | | Bottom | 4.024 (-0.8469 to 8.895) | % NT Activation |
| | | | $G_{\alpha}q^{N244G}$ | $IC_{50}$ | 6.488e-7 (4.873e-7 to 8.638e-7) | M |
| | | | | Top | 105.7 (101.4 to 110.0) | % NT Activation |
| | | | | Bottom | 14.26 (9.590 to 18.94) | % NT Activation |
| | | | $G_{\alpha}q^{Y243C}$ | $IC_{50}$ | 1.637e-007 (1.176e-007 to 2.279e-007) | M |
| | | | | Top | 102.7 (95.22 to 110.3) | % NT Activation |
| | | | | Bottom | 2.096 (-2.288 to 6.479) | % NT Activation |
| | | | $G_{\alpha}q^{L238N}$ | $IC_{50}$ | 6.797e-008 (4.490e-008 to 1.029e-007) | M |
| | | | | Top | 210.1 (13.41 to 406.8) | % NT Activation |
| | | | | Bottom | 102.4 (93.36 to 111.5) | % NT Activation |
| **4** | **J** | Structures of NTSR1-GoA (left), NTSR1-SBI-553-GoA (middle), and NTSR1-Gq (right) | | | | |
| **4** | **K** | Illustration | | | | |
| **4** | **L** | Illustration | | | | |
| **4** | **M** | Illustration | | | | |
| **4** | **N** | NTSR1-SBI-553-G protein 'open' position free energy of dissociation categorized by extent of SBI-553 inhibition | | | | |
| **4** | **O** | Correlation of NTSR1-SBI-553-G protein 'open' position free energy of dissociation vs. SBI-553-induced change in NT potency correlation | | | | |
| | | Null hypothesis | | | Slope = 0 | |
| | | Alternative hypothesis | | | Slope unconstrained | |
| | | P value | | | <0.0001 | |
| | | Conclusion (alpha = 0.05) | | | Reject null hypothesis | |
| | | Preferred model | | | Slope unconstrained | |
| | | F (DFn, DFd) | | | 164.4 (1, 7) | |
| | | Slope unconstrained | | | | |
| | | Best-fit values | | | | |
| | | Y-Intercept | | | 0.4434 | |
| | | Slope | | | 0.3976 | |
| | | 95% CI (profile likelihood) | | | | |
| | | Y-Intercept | | | 0.3187 to 0.5681 | |
| | | Slope | | | 0.3242 to 0.4709 | |
| | | Goodness of Fit | | | | |
| | | Degrees of Freedom | | | 7 | |
| | | R squared | | | 0.9591 | |
| **4** | **P** | Correlation of NTSR1-SBI-553-G protein 'open' position free energy of dissociation vs. max SBI-553-induced delta BRET in TRUPATH assay of G protein activation | | | | |
| | | Comparison of Fits | | | | |
| | | Null hypothesis | | | Slope = 0 | |
| | | Alternative hypothesis | | | Slope unconstrained | |
| | | P value | | | 0.0005 | |
| | | Conclusion (alpha = 0.05) | | | Reject null hypothesis | |
| | | Preferred model | | | Slope unconstrained | |
| | | F (DFn, DFd) | | | 37.99 (1, 7) | |

| | | | |
|---|---|---|---|
| | | Slope unconstrained | |
| | | Best-fit values | |
| | | Y-Intercept | 0.4587 |
| | | Slope | 5.387 |
| | | 95% CI (profile likelihood) | |
| | | Y-Intercept | 0.2107 to 0.7067 |
| | | Slope | 3.320 to 7.453 |
| | | Goodness of Fit | |
| | | Degrees of Freedom | 7 |
| | | R squared | 0.8444 |
| **4** | **Q** | Illustration | |

CI, confidence interval; NT, neurotensin; SBI-553, SBI-0654553; $EC_{50}$, half-maximal effective concentration; ND, not defined

**Table S5. Curve parameters and statistical comparisons presented in Figure 5 – Supporting Figure 5.**

| Fig | Panel | Description | N | Condition | Parameter | Mean (95% CI) | Units |
|---|---|---|---|---|---|---|---|
| **5** | **A** | Structure Image | | | | | |
| **5** | **B** | SAR summary | | | | | |
| **5** | **C** | Comparison of SBI-553 and SBI-342 structures | | | | | |
| **5** | **D** | Effect of SBI-553 vs. SBI-342 on NT (100 nM)-induced activation of $G_{\alpha}q$ | 3 | SBI-553 | $IC_{50}$ | 1.976e-6 (1.413e-6 to 2.764e-6) | M |
| | | | | | Top | 0.2546 (0.2431 to 0.2661) | -Δ Net BRET |
| | | | | | Bottom | 0.006087 (-0.01280 to 0.02497) | -Δ Net BRET |
| | | | | SBI-342 | $IC_{50}$ | 6.659e-6 (4.403e-6 to 1.007e-5) | M |
| | | | | | Top | 0.2594 (0.2526 to 0.2661) | -Δ Net BRET |
| | | | | | Bottom | 0.06608 (0.04050 to 0.09166) | -Δ Net BRET |
| | | Effect of SBI-553 vs. SBI-342 on NT (100 nM)-induced activation of $G_{\alpha}i1$ | 3 | SBI-553 | $IC_{50}$ | 2.261e-7 (7.741e-008 to 6.604e-7) | M |
| | | | | | Top | 0.2530 (0.2339 to 0.2720) | -Δ Net BRET |
| | | | | | Bottom | 0.1778 (0.1672 to 0.1885) | -Δ Net BRET |
| | | | | SBI-342 | $IC_{50}$ | 1.481e-6 (7.568e-7 to 2.898e-6) | M |
| | | | | | Top | 0.2650 (0.2556 to 0.2743) | -Δ Net BRET |
| | | | | | Bottom | 0.1646 (0.1501 to 0.1791) | -Δ Net BRET |
| | | Effect of SBI-553 vs. SBI-342 on NT (100 nM)-induced activation of $G_{\alpha}oA$ | 3 | SBI-553 | $IC_{50}$ | 8.548e-006 (4.621e-007 to 0.0001581) | M |
| | | | | | Top | 0.1877 (0.1717 to 0.2037) | -Δ Net BRET |
| | | | | | Bottom | 0.2533 (0.1880 to 0.3186) | -Δ Net BRET |
| | | | | SBI-342 | $IC_{50}$ | 5.841e-6 (2.127e-6 to 1.604e-5) | M |
| | | | | | Top | 0.2098 (0.2010 to 0.2186) | -Δ Net BRET |
| | | | | | Bottom | 0.1127 (0.08263 to 0.1428) | -Δ Net BRET |
| | | Effect of SBI-553 vs. SBI-342 on NT (100 nM)-induced activation of $G_{\alpha}12$ | 3 | SBI-553 | $IC_{50}$ | 2.400e-008 (1.183e-009 to 4.870e-007) | M |
| | | | | | Top | 0.1544 (0.08356 to 0.2253) | -Δ Net BRET |
| | | | | | Bottom | 0.2183 (0.2080 to 0.2285) | -Δ Net BRET |
| | | | | SBI-342 | $IC_{50}$ | 1.036e-005 (1.238e-009 to 0.08676) | M |
| | | | | | Top | 0.1782 (0.1605 to 0.1959) | -Δ Net BRET |
| | | | | | Bottom | 0.2074 (0.1091 to 0.3057) | -Δ Net BRET |
| **5** | **E** | SBI-553 and SBI-342-induced recruitment of β-arrestin2 | 3 | SBI-553 | $EC_{50}$ | 5.991e-6 (5.018e-6 to 7.153e-6) | M |
| | | | | | Top | 0.6339 (0.5995 to 0.6683) | Δ Net BRET |
| | | | | | Bottom | 0.003077 (-0.007140 to 0.01329) | Δ Net BRET |
| | | | | SBI-342 | $EC_{50}$ | 7.563e-6 (6.020e-6 to 9.500e-6) | M |
| | | | | | Top | 0.6202 (0.5728 to 0.6675) | Δ Net BRET |
| | | | | | Bottom | -0.001912 (-0.01318 to 0.009359) | Δ Net BRET |
| **5** | **F** | Effect of SBI-553 vs. SBI- | 6 | NT + 30 μM SBI-553 | $EC_{50}$ | 1.385e-008 (8.070e-9 to 2.377e-8) | M |

| | | | | | | | |
|---|---|---|---|---|---|---|---|
| | | 342 on NT-induced activation of $G_{\alpha}oA$ (NT dose-response) | | | Top | 0.1710 (0.1631 to 0.1790) | -Δ Net BRET |
| | | | | | Bottom | 0.07148 (0.06506 to 0.07791) | -Δ Net BRET |
| | | | | NT + 30 µM SBI-342 | $EC_{50}$ | 2.175e-8 (8.085e-9 to 5.850e-8) | M |
| | | | | | Top | 0.08451 (0.06807 to 0.1009) | -Δ Net BRET |
| | | | | | Bottom | -0.02726 (-0.03957 to -0.01495) | -Δ Net BRET |
| | | | | NT + Vehicle | $EC_{50}$ | 8.244e-9 (5.406e-9 to 1.257e-8) | M |
| | | | | | Top | 0.1764 (0.1646 to 0.1882) | -Δ Net BRET |
| | | | | | Bottom | -0.01807 (-0.02826 to -0.007876) | -Δ Net BRET |
| **5** | **G** | Effect of SBI-342 on $G_{\alpha}oA$ activation in non-NTSR1-expressing HEK293T cells | 3 | | Slope | 0.004923 (0.0003748 to 0.009472) | -Δ Net BRET |
| | | | | | Y-intercept | 0.04403 (0.01508 to 0.07298) | -Δ Net BRET |
| **5** | **H** | Effect of SBI-342 on dopamine-induced activation of $G_{\alpha}oA$ in D2-expressing HEK293T cells | 3 | Dopamine + Vehicle | $EC_{50}$ | 6.066e-8 (4.393e-8 to 8.496e-8) | M |
| | | | | | Top | 0.2371 (0.2266 to 0.2492) | -Δ Net BRET |
| | | | | | Bottom | -0.0007679 (-0.01027 to 0.007998) | -Δ Net BRET |
| | | | | Dopamine + SBI-342 | $EC_{50}$ | 6.663e-9 (4.133e-9 to 1.051e-8) | M |
| | | | | | Top | 0.2871 (0.2736 to 0.3024) | -Δ Net BRET |
| | | | | | Bottom | -0.006840 (-0.03060 to 0.01191) | -Δ Net BRET |
| **5** | **I** | Effect of SBI-553 vs. SBI-342 on NT (100 nM)-induced activation of $G_{\alpha}o$ Shedding | 4 | SBI-553 | $IC_{50}$ | 1.426e-7 (8.519e-8 to 2.893e-7) | M |
| | | | | | Top | 14.60 (13.94 to 15.26) | % AP Shedding |
| | | | | | Bottom | 10.95 (9.954 to 11.79) | % AP Shedding |
| | | | | SBI-342 | $IC_{50}$ | 5.636e-7 (4.096e-007 to 8.037e-7) | M |
| | | | | | Top | 13.72 (12.95 to 14.56) | % AP Shedding |
| | | | | | Bottom | 0.5828 (-1.160 to 1.859) | % AP Shedding |
| **5** | **J** | Effect of SBI-553 vs. SBI-342 on NT (100 nM)-induced mini $G_{\alpha}o$ Recruitment | 3 | SBI-553 | $IC_{50}$ | 1.214e-7 (ND) | M |
| | | | | | Top | 0.02417 (0.02143 to NA) | Δ Net BRET |
| | | | | | Bottom | 0.01855 (0.01650 to 0.02025) | Δ Net BRET |
| | | | | SBI-342 | $IC_{50}$ | 9.144e-7 (2.465e-007 to 3.196e-6) | M |
| | | | | | Top | 0.02573 (0.02185 to 0.03053) | Δ Net BRET |
| | | | | | Bottom | 0.004451 (-0.0009533 to 0.008759) | Δ Net BRET |
| **5** | **K** | Structure Image | | | | | |
| **5** | **L** | Comparison of SBI-553 and SBI-593 structures | | | | | |
| **5** | **M** | Effect of SBI-553 vs. SBI-593 on NT (100 nM)-induced activation of $G_{\alpha}q$ | 3 | SBI-553 | $IC_{50}$ | 1.976e-6 (1.413e-6 to 2.764e-6) | M |
| | | | | | Top | 0.2546 (0.2431 to 0.2661) | -Δ Net BRET |
| | | | | | Bottom | 0.006087 (-0.01280 to 0.02497) | -Δ Net BRET |
| | | | | SBI-593 | $IC_{50}$ | 3.227e-006 (1.533e-006 to 6.791e-006) | M |
| | | | | | Top | 0.2533 (0.2456 to 0.2610) | -Δ Net BRET |

| | | | | | | | |
|---|---|---|---|---|---|---|---|
| | | | | | Bottom | 0.1612 (0.1434 to 0.1790) | -Δ Net BRET |
| | | Effect of SBI-553 vs. SBI-593 on NT (100nM)-induced activation of $G_{\alpha}i1$ | 3 | SBI-553 | $IC_{50}$ | 2.261e-7 (7.741e-008 to 6.604e-7) | M |
| | | | | | Top | 0.2530 (0.2339 to 0.2720) | -Δ Net BRET |
| | | | | | Bottom | 0.1778 (0.1672 to 0.1885) | -Δ Net BRET |
| | | | | SBI-593 | $IC_{50}$ | 1.048e-006 (4.066e-007 to 2.703e-006) | M |
| | | | | | Top | 0.2617 (0.2521 to 0.2713) | -Δ Net BRET |
| | | | | | Bottom | 0.1940 (0.1813 to 0.2068) | -Δ Net BRET |
| | | Effect of SBI-553 vs. SBI-593 on NT (100 nM)-induced activation of $G_{\alpha}oA$ | 3 | SBI-553 | $IC_{50}$ | 8.548e-006 (4.621e-007 to 0.0001581) | M |
| | | | | | Top | 0.1877 (0.1717 to 0.2037) | -Δ Net BRET |
| | | | | | Bottom | 0.2533 (0.1880 to 0.3186) | -Δ Net BRET |
| | | | | SBI-593 | $IC_{50}$ | 2.029e-005 (7.550e-034 to 5.455e+023) | M |
| | | | | | Top | 0.2044 (0.1950 to 0.2138) | -Δ Net BRET |
| | | | | | Bottom | 0.1323 (-3.984 to 4.249) | -Δ Net BRET |
| | | Effect of SBI-553 vs. SBI-593 on NT (100 nM)-induced activation of $G_{\alpha}12$ | 3 | SBI-553 | $IC_{50}$ | 2.400e-008 (1.183e-009 to 4.870e-007) | M |
| | | | | | Top | 0.1544 (0.08356 to 0.2253) | -Δ Net BRET |
| | | | | | Bottom | 0.2183 (0.2080 to 0.2285) | -Δ Net BRET |
| | | | | SBI-593 | $IC_{50}$ | 1.634e-007 (2.287e-009 to 1.168e-005) | M |
| | | | | | Top | 0.1715 (0.1246 to 0.2183) | -Δ Net BRET |
| | | | | | Bottom | 0.2200 (0.1927 to 0.2473) | -Δ Net BRET |
| **5** | **N** | SBI-553 and SBI-593-induced recruitment of β-arrestin2 | 3 | SBI-553 | $EC_{50}$ | 5.991e-6 (5.018e-6 to 7.153e-6) | M |
| | | | | | Top | 0.6339 (0.5995 to 0.6683) | Δ Net BRET |
| | | | | | Bottom | 0.003077 (-0.007140 to 0.01329) | Δ Net BRET |
| | | | | SBI-593 | $EC_{50}$ | 4.750e-6 (3.924e-6 to 5.750e-6) | M |
| | | | | | Top | 0.4837 (0.4565 to 0.5110) | Δ Net BRET |
| | | | | | Bottom | -0.01077 (-0.01999 to -0.001550) | Δ Net BRET |
| **5** | **O** | Effect of SBI-553 vs. SBI-593 on NT-induced activation of $G_{\alpha}q$ (NT dose-response) | 3 | NT + 30 μM SBI-553 | $EC_{50}$ | 5.053e-007 (5.776e-018 to 44201) | M |
| | | | | | Top | 0.004370 (-0.01697 to 0.02571) | -Δ Net BRET |
| | | | | | Bottom | 0.0004696 (-0.007560 to 0.008499) | -Δ Net BRET |
| | | | | NT + 30 μM SBI-593 | $EC_{50}$ | 3.371e-8 (2.240e-008 to 5.074e-008) | M |
| | | | | | Top | 0.1924 (0.1799 to 0.2049) | -Δ Net BRET |
| | | | | | Bottom | -0.007901 (-0.01652 to 0.0007175) | -Δ Net BRET |
| | | | | NT + Vehicle | $EC_{50}$ | 1.169e-8 (7.280e-9 to 1.878e-8) | M |
| | | | | | Top | 0.2674 (0.2480 to 0.2869) | -Δ Net BRET |
| | | | | | Bottom | -0.01092 (-0.02697 to 0.005141) | -Δ Net BRET |
| **5** | **P** | Effect of SBI-593 on $G_{\alpha}q$ activation in non-NTSR1- | 3 | SBI-593 | Slope | -0.002987 (-0.008509 to 0.002536) | -Δ Net BRET/M |
| | | | | | Y-intercept | -0.01328 (-0.04842 to 0.02187) | -Δ Net BRET |

| | | | | | | | |
|---|---|---|---|---|---|---|---|
| | | expressing HEK293T cells | | | | | |
| **5** | **Q** | Effect of SBI-553 vs. SBI-593 on NT (100 nM)-induced activation of $G_{\alpha}q$ Shedding | 3 | SBI-553 | $EC_{50}$ | 6.767e-7 (4.664e-7 to 1.002e-6) | M |
| | | | | | Top | 19.59 (18.78 to 20.41) | % AP Shedding |
| | | | | | Bottom | -2.984 (-6.236 to -0.3645) | % AP Shedding |
| | | | | SBI-593 | $EC_{50}$ | 4.725e-7 (2.997e-7 to 7.434e-7) | M |
| | | | | | Top | 19.79 (18.83 to 20.78) | % AP Shedding |
| | | | | | Bottom | 4.654 (3.070 to 6.095) | % AP Shedding |
| **5** | **R** | Effect of SBI-553 vs. SBI-593 on NT (100 nM)-induced mini $G_{\alpha}q$ Recruitment | 3 | SBI-553 | $EC_{50}$ | 4.767e-008 (3.671e-008 to 6.138e-008) | M |
| | | | | | Top | 0.1764 (0.1663 to 0.1884) | ∆ Net BRET |
| | | | | | Bottom | 0.03563 (0.03282 to 0.03839) | ∆ Net BRET |
| | | | | SBI-593 | $EC_{50}$ | 3.502e-007 (2.801e-007 to 4.371e-007) | M |
| | | | | | Top | 0.1432 (0.1397 to 0.1468) | ∆ Net BRET |
| | | | | | Bottom | 0.05822 (0.05527 to 0.06111) | ∆ Net BRET |
| **5** | **S** | Structure image | | | | | |
| **5** | **T** | Structure image | | | | | |
| **5** | **U** | Structure image | | | | | |
| **5** | **V** | Structure image | | | | | |
| **5** | **W** | NT + SBI-553 $G_{\alpha}q$ | 3 | Vehicle | NT $EC_{50}$ | 5.755e-9 (3.560e-9 to 9.303e-9) | M |
| | | | | | NT Top | 0.3072 (0.2868 to 0.3275) | -∆ Net BRET |
| | | | | | Bottom | -0.003109 (-0.02157 to 0.01535) | -∆ Net BRET |
| | | | | 1 µM SBI-553 | NT $EC_{50}$ | 1.784e-8 (7.137e-9 to 4.458e-8) | M |
| | | | | | NT Top | 0.1883 (0.1634 to 0.2132) | -∆ Net BRET |
| | | | | | Bottom | 0.004988 (-0.01432 to 0.02429) | -∆ Net BRET |
| | | | | 3 µM SBI-553 | NT $EC_{50}$ | 2.444e-8 (5.071e-9 to 1.178e-7) | M |
| | | | | | NT Top | 0.09894 (0.07403 to 0.1239) | -∆ Net BRET |
| | | | | | Bottom | -0.007306 (-0.02556 to 0.01095) | -∆ Net BRET |
| | | | | 10 µM SBI-553 | NT $EC_{50}$ | 3.279e-8 (2.555e-10 to 4.210e-6) | M |
| | | | | | NT Top | 0.01870 (-0.001856 to 0.03925) | -∆ Net BRET |
| | | | | | Bottom | -0.009091 (-0.02332 to 0.005141) | -∆ Net BRET |
| | | | | 30 µM SBI-553 | NT $EC_{50}$ | 5.053e-7 (5.776e-18 to 44201) | M |
| | | | | | NT Top | 0.004370 (-0.01697 to 0.02571) | -∆ Net BRET |
| | | | | | Bottom | 0.0004696 (-0.007560 to 0.008499) | -∆ Net BRET |
| | | NT + SBI-553 $G_{\alpha}i1$ | 3 | Vehicle | NT $EC_{50}$ | 4.989e-009 (2.837e-009 to 8.772e-009) | M |
| | | | | | NT Top | 0.2681 (0.2486 to 0.2876) | -∆ Net BRET |
| | | | | | Bottom | 0.009071 (-0.009004 to 0.02715) | -∆ Net BRET |

| | | | | | | |
|---|---|---|---|---|---|---|
| | | | 1 μM SBI-553 | NT $EC_{50}$ | 6.738e-009 (4.318e-009 to 1.052e-008) | M |
| | | | | NT Top | 0.2282 (0.2139 to 0.2426) | -Δ Net BRET |
| | | | | Bottom | -0.001507 (-0.01421 to 0.01119) | -Δ Net BRET |
| | | | 3 μM SBI-553 | NT $EC_{50}$ | 6.691e-009 (4.546e-009 to 9.848e-009) | M |
| | | | | NT Top | 0.2242 (0.2120 to 0.2364) | -Δ Net BRET |
| | | | | Bottom | -0.0009451 (-0.01175 to 0.009864) | -Δ Net BRET |
| | | | 10 μM SBI-553 | NT $EC_{50}$ | 5.301e-009 (3.713e-009 to 7.568e-009) | M |
| | | | | NT Top | 0.2205 (0.2106 to 0.2305) | -Δ Net BRET |
| | | | | Bottom | 0.01236 (0.003189 to 0.02153) | -Δ Net BRET |
| | | | 30 μM SBI-553 | NT $EC_{50}$ | 7.585e-009 (4.907e-009 to 1.172e-008) | M |
| | | | | NT Top | 0.2159 (0.2055 to 0.2264) | -Δ Net BRET |
| | | | | Bottom | 0.04686 (0.03771 to 0.05601) | -Δ Net BRET |
| | NT + SBI-553 $G_{\alpha}oA$ | 3 | Vehicle | $EC_{50}$ | 1.315e-008 (9.775e-009 to 1.769e-008) | M |
| | | | | Top | 0.2966 (0.2833 to 0.3099) | -Δ Net BRET |
| | | | | Bottom | -0.006282 (-0.01708 to 0.004518) | -Δ Net BRET |
| | | | 1 μM SBI-0654553 | $EC_{50}$ | 1.450e-008 (1.091e-008 to 1.927e-008) | M |
| | | | | Top | 0.2735 (0.2622 to 0.2849) | -Δ Net BRET |
| | | | | Bottom | 0.004023 (-0.005073 to 0.01312) | -Δ Net BRET |
| | | | 3 μM SBI-0654553 | $EC_{50}$ | 1.252e-008 (9.185e-009 to 1.706e-008) | M |
| | | | | Top | 0.2826 (0.2704 to 0.2947) | -Δ Net BRET |
| | | | | Bottom | 0.01642 (0.006461 to 0.02638) | -Δ Net BRET |
| | | | 10 μM SBI-0654553 | $EC_{50}$ | 1.468e-008 (9.237e-009 to 2.332e-008) | M |
| | | | | Top | 0.2868 (0.2716 to 0.3021) | -Δ Net BRET |
| | | | | Bottom | 0.06481 (0.05263 to 0.07700) | -Δ Net BRET |
| | | | 30 μM SBI-0654553 | $EC_{50}$ | 1.385e-008 (8.069e-009 to 2.377e-008) | M |
| | | | | Top | 0.2802 (0.2671 to 0.2932) | -Δ Net BRET |
| | | | | Bottom | 0.1171 (0.1066 to 0.1276) | -Δ Net BRET |
| | NT + SBI-553 $G_{\alpha}12$ | 3 | Vehicle | $EC_{50}$ | 1.877e-008 (1.093e-008 to 3.222e-008) | M |
| | | | | Top | 0.3035 (0.2796 to 0.3274) | -Δ Net BRET |
| | | | | Bottom | 0.005665 (-0.01269 to 0.02402) | -Δ Net BRET |
| | | | 1 μM SBI-0654553 | $EC_{50}$ | 8.984e-009 (5.167e-009 to 1.562e-008) | M |
| | | | | Top | 0.2976 (0.2746 to 0.3206) | -Δ Net BRET |
| | | | | Bottom | 0.01100 (-0.008638 to 0.03064) | -Δ Net BRET |
| | | | 3 μM SBI-0654553 | $EC_{50}$ | 6.201e-009 (3.266e-009 to 1.177e-008) | M |
| | | | | Top | 0.2988 (0.2755 to 0.3222) | -Δ Net BRET |
| | | | | Bottom | 0.03520 (0.01424 to 0.05617) | -Δ Net BRET |

| | | | | | | |
|---|---|---|---|---|---|---|
| | | | 10 μM SBI-0654553 | $EC_{50}$ | 7.388e-009 (2.891e-009 to 1.888e-008) | M |
| | | | | Top | 0.2954 (0.2653 to 0.3255) | -Δ Net BRET |
| | | | | Bottom | 0.06917 (0.04278 to 0.09555) | -Δ Net BRET |
| | | | 30 μM SBI-0654553 | $EC_{50}$ | 9.441e-009 (2.080e-009 to 4.285e-008) | M |
| | | | | Top | 0.2875 (0.2532 to 0.3218) | -Δ Net BRET |
| | | | | Bottom | 0.1320 (0.1029 to 0.1611) | -Δ Net BRET |
| | NT + SBI-553 β-arrestin2 | 3 | Vehicle | $EC_{50}$ | 1.462e-008 (1.119e-008 to 1.911e-008) | M |
| | | | | Top | 0.5629 (0.5404 to 0.5853) | Δ Net BRET |
| | | | | Bottom | -0.001865 (-0.01979 to 0.01606) | Δ Net BRET |
| | | | 1 μM SBI-0647342 | $EC_{50}$ | 1.381e-008 (1.016e-008 to 1.878e-008) | M |
| | | | | Top | 0.6212 (0.5937 to 0.6487) | Δ Net BRET |
| | | | | Bottom | 0.01863 (-0.003501 to 0.04076) | Δ Net BRET |
| | | | 3 μM SBI-0647342 | $EC_{50}$ | 1.486e-008 (1.036e-008 to 2.130e-008) | M |
| | | | | Top | 0.6242 (0.5960 to 0.6523) | Δ Net BRET |
| | | | | Bottom | 0.09722 (0.07477 to 0.1197) | Δ Net BRET |
| | | | 10 μM SBI-0647342 | $EC_{50}$ | 2.135e-008 (1.248e-008 to 3.651e-008) | M |
| | | | | Top | 0.6198 (0.5939 to 0.6456) | Δ Net BRET |
| | | | | Bottom | 0.2960 (0.2766 to 0.3154) | Δ Net BRET |
| | | | 30 μM SBI-0647342 | $EC_{50}$ | 2.337e-008 (9.356e-009 to 5.837e-008) | M |
| | | | | Top | 0.6376 (0.6188 to 0.6564) | Δ Net BRET |
| | | | | Bottom | 0.4998 (0.4859 to 0.5137) | Δ Net BRET |
| | NT + SBI-342 $G_{\alpha}q$ | 3 | Vehicle | $EC_{50}$ | 4.336e-009 (2.782e-009 to 6.759e-009) | M |
| | | | | Top | 0.2824 (0.2652 to 0.2997) | -Δ Net BRET |
| | | | | Bottom | -0.02626 (-0.05017 to -0.002348) | -Δ Net BRET |
| | | | 1 μM SBI-0647342 | $EC_{50}$ | 8.881e-009 (5.039e-009 to 1.565e-008) | M |
| | | | | Top | 0.2798 (0.2548 to 0.3047) | -Δ Net BRET |
| | | | | Bottom | -0.02419 (-0.04554 to -0.002847) | -Δ Net BRET |
| | | | 3 μM SBI-0647342 | $EC_{50}$ | 1.233e-008 (7.876e-009 to 1.931e-008) | M |
| | | | | Top | 0.2555 (0.2367 to 0.2743) | -Δ Net BRET |
| | | | | Bottom | -0.02851 (-0.04393 to -0.01309) | -Δ Net BRET |
| | | | 10 μM SBI-0647342 | $EC_{50}$ | 2.304e-008 (1.294e-008 to 4.102e-008) | M |
| | | | | Top | 0.1846 (0.1665 to 0.2028) | -Δ Net BRET |
| | | | | Bottom | -0.02695 (-0.04041 to -0.01350) | -Δ Net BRET |
| | | | 30 μM SBI-0647342 | $EC_{50}$ | 2.897e-008 (1.148e-008 to 7.310e-008) | M |
| | | | | Top | 0.08891 (0.07448 to 0.1033) | -Δ Net BRET |
| | | | | Bottom | -0.01462 (-0.02485 to -0.004380) | -Δ Net BRET |

| | | | | | | |
|---|---|---|---|---|---|---|
| | NT + SBI-342 $G_{\alpha}i1$ | 3 | 0 µM SBI-0647342 | $EC_{50}$ | 2.202e-009 (1.282e-009 to 3.780e-009) | M |
| | | | | Top | 0.2639 (0.2458 to 0.2821) | -Δ Net BRET |
| | | | | Bottom | -0.007692 (-0.02698 to 0.01159) | -Δ Net BRET |
| | | | 1 µM SBI-0647342 | $EC_{50}$ | 5.887e-009 (4.540e-009 to 7.634e-009) | M |
| | | | | Top | 0.2245 (0.2165 to 0.2325) | -Δ Net BRET |
| | | | | Bottom | 0.0009526 (-0.006248 to 0.008153) | -Δ Net BRET |
| | | | 3 µM SBI-0647342 | $EC_{50}$ | 8.005e-009 (6.081e-009 to 1.054e-008) | M |
| | | | | Top | 0.2115 (0.2027 to 0.2203) | -Δ Net BRET |
| | | | | Bottom | -0.01253 (-0.02017 to -0.004879) | -Δ Net BRET |
| | | | 10 µM SBI-0647342 | $EC_{50}$ | 1.154e-008 (8.985e-009 to 1.483e-008) | M |
| | | | | Top | 0.1870 (0.1798 to 0.1942) | -Δ Net BRET |
| | | | | Bottom | -0.008602 (-0.01458 to -0.002628) | -Δ Net BRET |
| | | | 30 µM SBI-0647342 | $EC_{50}$ | 8.967e-009 (5.453e-009 to 1.474e-008) | M |
| | | | | Top | 0.1774 (0.1651 to 0.1897) | -Δ Net BRET |
| | | | | Bottom | 0.006582 (-0.003943 to 0.01711) | -Δ Net BRET |
| | NT + SBI-342 $G_{\alpha}oA$ | 3 | 0 µM SBI-0647342 | $EC_{50}$ | 8.244e-009 (5.406e-009 to 1.257e-008) | M |
| | | | | Top | 0.2044 (0.1926 to 0.2162) | -Δ Net BRET |
| | | | | Bottom | 0.009921 (-0.0002703 to 0.02011) | -Δ Net BRET |
| | | | 1 µM SBI-0647342 | $EC_{50}$ | 2.045e-008 (1.183e-008 to 3.534e-008) | M |
| | | | | Top | 0.1749 (0.1608 to 0.1890) | -Δ Net BRET |
| | | | | Bottom | 0.001402 (-0.009271 to 0.01208) | -Δ Net BRET |
| | | | 3 µM SBI-0647342 | $EC_{50}$ | 1.886e-008 (1.050e-008 to 3.388e-008) | M |
| | | | | Top | 0.1517 (0.1379 to 0.1655) | -Δ Net BRET |
| | | | | Bottom | -0.006902 (-0.01749 to 0.003683) | -Δ Net BRET |
| | | | 10 µM SBI-0647342 | $EC_{50}$ | 1.870e-008 (1.039e-008 to 3.365e-008) | M |
| | | | | Top | 0.1234 (0.1123 to 0.1345) | -Δ Net BRET |
| | | | | Bottom | -0.003635 (-0.01215 to 0.004879) | -Δ Net BRET |
| | | | 30 µM SBI-0647342 | $EC_{50}$ | 2.175e-008 (8.085e-009 to 5.850e-008) | M |
| | | | | Top | 0.1125 (0.09606 to 0.1289) | -Δ Net BRET |
| | | | | Bottom | 0.0007274 (-0.01158 to 0.01303) | -Δ Net BRET |
| | NT + SBI-342 $G_{\alpha}12$ | 3 | 0 µM SBI-0647342 | $EC_{50}$ | 6.205e-009 (2.552e-009 to 1.436e-008) | M |
| | | | | Top | 0.2614 (0.2310 to 0.2968) | -Δ Net BRET |
| | | | | Bottom | -0.006051 (-0.03728 to 0.02223) | -Δ Net BRET |
| | | | 1 µM SBI-0647342 | $EC_{50}$ | 9.411e-009 (3.416e-009 to 2.429e-008) | M |
| | | | | Top | 0.2509 (0.2162 to 0.2932) | -Δ Net BRET |
| | | | | Bottom | 0.01205 (-0.02269 to 0.04264) | -Δ Net BRET |

| | | | | | | |
|---|---|---|---|---|---|---|
| | | | 3 µM SBI-0647342 | $EC_{50}$ | 1.116e-008 (3.928e-009 to 3.402e-008) | M |
| | | | | Top | 0.2545 (0.2162 to 0.3092) | -Δ Net BRET |
| | | | | Bottom | -0.004815 (-0.04246 to 0.02680) | -Δ Net BRET |
| | | | 10 µM SBI-0647342 | $EC_{50}$ | 1.123e-008 (3.652e-009 to 3.583e-008) | M |
| | | | | Top | 0.2343 (0.1971 to 0.2875) | -Δ Net BRET |
| | | | | Bottom | -0.01150 (-0.04825 to 0.01963) | -Δ Net BRET |
| | | | 30 µM SBI-0647342 | $EC_{50}$ | 1.012e-008 (4.937e-009 to 2.142e-008) | M |
| | | | | Top | 0.2368 (0.2131 to 0.2675) | -Δ Net BRET |
| | | | | Bottom | -0.001556 (-0.02374 to 0.01811) | -Δ Net BRET |
| | NT + SBI-342<br>β-arrestin2 | 3 | 0 µM SBI-0647342 | $EC_{50}$ | 8.855e-009 (6.716e-009 to 1.167e-008) | M |
| | | | | Top | 0.6037 (0.5794 to 0.6280) | Δ Net BRET |
| | | | | Bottom | -0.003028 (-0.02381 to 0.01776) | Δ Net BRET |
| | | | 1 µM SBI-0647342 | $EC_{50}$ | 8.770e-009 (6.397e-009 to 1.202e-008) | M |
| | | | | Top | 0.6080 (0.5834 to 0.6327) | Δ Net BRET |
| | | | | Bottom | 0.06818 (0.04707 to 0.08930) | Δ Net BRET |
| | | | 3 µM SBI-0647342 | $EC_{50}$ | 5.589e-009 (3.863e-009 to 8.086e-009) | M |
| | | | | Top | 0.6735 (0.6497 to 0.6973) | Δ Net BRET |
| | | | | Bottom | 0.1995 (0.1778 to 0.2212) | Δ Net BRET |
| | | | 10 µM SBI-0647342 | $EC_{50}$ | 5.141e-009 (2.896e-009 to 9.126e-009) | M |
| | | | | Top | 0.7051 (0.6845 to 0.7257) | Δ Net BRET |
| | | | | Bottom | 0.4376 (0.4186 to 0.4566) | Δ Net BRET |
| | | | 30 µM SBI-0647342 | $EC_{50}$ | 9.188e-009 (1.757e-009 to 4.806e-008) | M |
| | | | | Top | 0.7913 (0.7652 to 0.8175 | Δ Net BRET |
| | | | | Bottom | 0.6825 (0.6602 to 0.7048) | Δ Net BRET |
| | NT + SBI-593<br>$G_{\alpha}q$ | 3 | 0 µM SBI-0646593 | $EC_{50}$ | 1.169e-008 (7.280e-009 to 1.878e-008) | M |
| | | | | Top | 0.2674 (0.2480 to 0.2869) | -Δ Net BRET |
| | | | | Bottom | -0.01092 (-0.02697 to 0.005141) | -Δ Net BRET |
| | | | 1 µM SBI-0646593 | $EC_{50}$ | 1.800e-008 (1.164e-008 to 2.784e-008) | M |
| | | | | Top | 0.2692 (0.2508 to 0.2876) | -Δ Net BRET |
| | | | | Bottom | -0.01557 (-0.02982 to -0.001323) | -Δ Net BRET |
| | | | 3 µM SBI-0646593 | $EC_{50}$ | 2.675e-008 (1.781e-008 to 4.017e-008) | M |
| | | | | Top | 0.2469 (0.2306 to 0.2633) | -Δ Net BRET |
| | | | | Bottom | -0.02159 (-0.03337 to -0.009811) | -Δ Net BRET |
| | | | 10 µM SBI-0646593 | $EC_{50}$ | 3.121e-008 (2.179e-008 to 4.468e-008) | M |
| | | | | Top | 0.2039 (0.1919 to 0.2158) | -Δ Net BRET |
| | | | | Bottom | -0.01599 (-0.02435 to -0.007625) | -Δ Net BRET |

| | | | | | | |
|---|---|---|---|---|---|---|
| | | | 30 μM SBI-0646593 | $EC_{50}$ | 3.371e-008 (2.240e-008 to 5.074e-008) | M |
| | | | | Top | 0.1924 (0.1799 to 0.2049) | -Δ Net BRET |
| | | | | Bottom | -0.007901 (-0.01652 to 0.0007175) | -Δ Net BRET |
| | NT + SBI-593 $G_\alpha$i1 | 3 | 0 μM SBI-0646593 | $EC_{50}$ | 1.063e-008 (7.131e-009 to 1.583e-008) | M |
| | | | | Top | 0.2613 (0.2457 to 0.2770) | -Δ Net BRET |
| | | | | Bottom | -0.006159 (-0.01926 to 0.006939) | -Δ Net BRET |
| | | | 1 μM SBI-0646593 | $EC_{50}$ | 1.147e-008 (8.229e-009 to 1.598e-008) | M |
| | | | | Top | 0.2252 (0.2131 to 0.2374) | -Δ Net BRET |
| | | | | Bottom | -0.02262 (-0.03266 to -0.01259) | -Δ Net BRET |
| | | | 3 μM SBI-0646593 | $EC_{50}$ | 1.386e-008 (9.491e-009 to 2.025e-008) | M |
| | | | | Top | 0.2042 (0.1911 to 0.2172) | -Δ Net BRET |
| | | | | Bottom | -0.02781 (-0.03831 to -0.01732) | -Δ Net BRET |
| | | | 10 μM SBI-0646593 | $EC_{50}$ | 1.561e-008 (1.086e-008 to 2.244e-008) | M |
| | | | | Top | 0.1951 (0.1839 to 0.2064) | -Δ Net BRET |
| | | | | Bottom | -0.01407 (-0.02298 to -0.005152) | -Δ Net BRET |
| | | | 30 μM SBI-0646593 | $EC_{50}$ | 1.466e-008 (9.419e-009 to 2.281e-008) | M |
| | | | | Top | 0.1933 (0.1806 to 0.2059) | -Δ Net BRET |
| | | | | Bottom | 0.0003866 (-0.009722 to 0.01050) | -Δ Net BRET |
| | NT + SBI-593 $G_\alpha$oA | 3 | 0 μM SBI-0646593 | $EC_{50}$ | 2.397e-008 (1.583e-008 to 3.630e-008) | M |
| | | | | Top | 0.2139 (0.2006 to 0.2271) | -Δ Net BRET |
| | | | | Bottom | -6.091e-008 (-0.009725 to 0.009724) | -Δ Net BRET |
| | | | 1 μM SBI-0646593 | $EC_{50}$ | 3.669e-008 (2.512e-008 to 5.358e-008) | M |
| | | | | Top | 0.2029 (0.1910 to 0.2147) | -Δ Net BRET |
| | | | | Bottom | 3.090e-006 (-0.008040 to 0.008047) | -Δ Net BRET |
| | | | 3 μM SBI-0646593 | $EC_{50}$ | 3.131e-008 (2.015e-008 to 4.865e-008) | M |
| | | | | Top | 0.2122 (0.1980 to 0.2263) | -Δ Net BRET |
| | | | | Bottom | 2.343e-007 (-0.009906 to 0.009906) | -Δ Net BRET |
| | | | 10 μM SBI-0646593 | $EC_{50}$ | 2.928e-008 (1.961e-008 to 4.373e-008) | M |
| | | | | Top | 0.2109 (0.1982 to 0.2237) | -Δ Net BRET |
| | | | | Bottom | 2.035e-006 (-0.009024 to 0.009028) | -Δ Net BRET |
| | | | 30 μM SBI-0646593 | $EC_{50}$ | 1.967e-008 (9.587e-009 to 4.036e-008) | M |
| | | | | Top | 0.1915 (0.1711 to 0.2120) | -Δ Net BRET |
| | | | | Bottom | 2.619e-006 (-0.01557 to 0.01558) | -Δ Net BRET |
| | NT + SBI-593 $G_\alpha$12 | 3 | 0 μM SBI-0646593 | $EC_{50}$ | 1.847e-008 (1.248e-008 to 2.736e-008) | M |
| | | | | Top | 0.1974 (0.1856 to 0.2091) | -Δ Net BRET |
| | | | | Bottom | -0.004860 (-0.01394 to 0.004216) | -Δ Net BRET |

| | | | | | | | |
|---|---|---|---|---|---|---|---|
| | | | | 1 μM SBI-0646593 | $EC_{50}$ | 2.191e-008 (1.512e-008 to 3.176e-008) | M |
| | | | | | Top | 0.1967 (0.1846 to 0.2088) | -Δ Net BRET |
| | | | | | Bottom | -0.02225 (-0.03129 to -0.01322) | -Δ Net BRET |
| | | | | 3 μM SBI-0646593 | $EC_{50}$ | 1.670e-008 (1.126e-008 to 2.476e-008) | M |
| | | | | | Top | 0.1927 (0.1797 to 0.2058) | -Δ Net BRET |
| | | | | | Bottom | -0.03068 (-0.04091 to -0.02045) | -Δ Net BRET |
| | | | | 10 μM SBI-0646593 | $EC_{50}$ | 1.797e-008 (1.254e-008 to 2.576e-008) | M |
| | | | | | Top | 0.1970 (0.1852 to 0.2087) | -Δ Net BRET |
| | | | | | Bottom | -0.02309 (-0.03219 to -0.01399) | -Δ Net BRET |
| | | | | 30 μM SBI-0646593 | $EC_{50}$ | 1.741e-008 (1.079e-008 to 2.809e-008) | M |
| | | | | | Top | 0.2002 (0.1863 to 0.2142) | -Δ Net BRET |
| | | | | | Bottom | 0.004165 (-0.006658 to 0.01499) | -Δ Net BRET |
| | | NT + SBI-593 | 3 | 0 μM SBI-0646593 | $EC_{50}$ | 1.906e-008 (1.138e-008 to 3.192e-008) | M |
| | | β-arrestin2 | | | Top | 0.4671 (0.4319 to 0.5022) | Δ Net BRET |
| | | | | | Bottom | 0.008224 (-0.01870 to 0.03515) | Δ Net BRET |
| | | | | 1 μM SBI-0646593 | $EC_{50}$ | 1.869e-008 (1.043e-008 to 3.349e-008) | M |
| | | | | | Top | 0.4973 (0.4596 to 0.5349) | Δ Net BRET |
| | | | | | Bottom | 0.06263 (0.03371 to 0.09156) | Δ Net BRET |
| | | | | 3 μM SBI-0646593 | $EC_{50}$ | 1.995e-008 (8.343e-009 to 4.772e-008) | M |
| | | | | | Top | 0.5570 (0.5059 to 0.6081) | Δ Net BRET |
| | | | | | Bottom | 0.1623 (0.1234 to 0.2011) | Δ Net BRET |
| | | | | 10 μM SBI-0646593 | $EC_{50}$ | 1.838e-008 (5.997e-009 to 5.636e-008) | M |
| | | | | | Top | 0.6422 (0.5945 to 0.6898) | Δ Net BRET |
| | | | | | Bottom | 0.3548 (0.3173 to 0.3922) | Δ Net BRET |
| | | | | 30 μM SBI-0646593 | $EC_{50}$ | 1.413e-008 (3.649e-009 to 5.471e-008) | M |
| | | | | | Top | 0.6318 (0.5825 to 0.6810) | Δ Net BRET |
| | | | | | Bottom | 0.3858 (0.3455 to 0.4262) | Δ Net BRET |
| **5** | **X** | Radar plot summary of G protein activation by SBI-553, SBI-342 and SBI-593 | | | | | |
| **5** | **Y** | Cartoon of G protein activation by SBI-553, SBI-342 and SBI-593 | | | | | |

CI, confidence interval; NT, neurotensin; SBI-553, SBI-0654553; M, molar; $EC_{50}$, half-maximal effective concentration; $IC_{50}$, half-maximal inhibitory concentration; NA, not applicable, ND, not defined

**Table S6. Curve parameters and statistical comparisons presented in Figure 6 – Supporting Figure 6.**

| Fig | Panel | Description | Statistical Test | F Statistics | Multiple Comparisons | N | Condition | Parameter | Mean (95% CI) | Units |
|---|---|---|---|---|---|---|---|---|---|---|
| **6** | **A** | Experimental timeline of PD149163 (i.p.)-induced hypothermia studies | | | | | | | | |
| **6** | **B** | PD149163-induced Hypothermia in WT mice – Time Course of Body Temperature | Two-way repeated measures ANOVA followed by Dunnett's multiple comparisons test | $F_{Time}$ (3.015, 132.0) = 64.22, p<0.0001<br>$F_{Treatment}$ (3, 46) = 67.62, p<0.0001<br>$F_{Interaction}$ (30, 438) = 23.03, p<0.0001 | *p<0.05, **p<0.01, ***p <0.001, PD149163 vs Vehicle | 9-17 | | | | |
| | | PD149163-induced Hypothermia in NTSR1$^{-/-}$ mice – Time Course of Body Temperature | Two-way repeated measures ANOVA | $F_{Time}$ (3, 26) = 2.728<br>$F_{Treatment}$ (4.669, 121.4) = 3.961<br>$F_{Interaction}$ (24, 208) = 2.689 | N/A | 6-8 | | | | |
| **6** | **C** | PD149163-induced Human NTSR1 $G_{\alpha}q$ Activation | | | | 3 | SBI-553 | $IC_{50}$ | 2.072e-015 (ND) | M |
| | | | | | | | | Top | 12.90 (ND) | ∆ Net BRET |
| | | | | | | | | Bottom | 0.02533 (0.007351 to 0.04034) | ∆ Net BRET |
| | | | | | | 3 | SBI-593 | $IC_{50}$ | 2.161e-007 (ND) | M |
| | | | | | | | | Top | 0.1474 (ND) | ∆ Net BRET |
| | | | | | | | | Bottom | 0.06322 (ND) | ∆ Net BRET |
| **6** | **D** | PD149163-induced Mouse NTSR1 $G_{\alpha}q$ Activation | | | | 3 | SBI-553 | $IC_{50}$ | 4.070e-014 (ND) | M |
| | | | | | | | | Top | 0.6686 (ND) | ∆ Net BRET |
| | | | | | | | | Bottom | 0.002194 (-0.006034 to 0.008608) | ∆ Net BRET |
| | | | | | | 3 | SBI-593 | $IC_{50}$ | 5.072e-007 (2.616e-007 to 8.596e-007) | M |
| | | | | | | | | Top | 0.1616 (0.1458 to 0.2396) | ∆ Net BRET |

| | | | | | | | | | |
|---|---|---|---|---|---|---|---|---|---|
| | | | | | | | Bottom | 0.05582 (0.01734 to 0.06830) | Δ Net BRET |
| **6** | **E** | Experimental timeline of studies evaluating the effect of SBI compounds (i.p.) on PD149163 (i.p.)-induced hypothermia | | | | | | | |
| **6** | **F** | SBI-553 Attenuation of PD149163-induced Hypothermia – Time Course of Body Temperature | Two-way repeated measures ANOVA, Time 30-300 min post PD149163 treatment | $F_{Time}$ (1.156, 16.19) = 28.50, $p<0.0001$<br><br>$F_{Treatment}$ (1, 14) = 4.671, $p=0.0485$<br><br>$F_{Interaction}$ (9, 126) = 2.163, $p= 0.0289$ | N/A | 8 | | | |
| | **G** | SBI-593 Attenuation of PD149163-induced Hypothermia – Time Course of Body Temperature | Two-way repeated measures ANOVA, Time 30-300 min post PD149163 treatment | $F_{Time}$ (2.445, 34.23) = 129.3, $p<0.0001$<br><br>$F_{Treatment}$ (1, 14) = 1.817, $p=0.1990$<br><br>$F_{Interaction}$ (14, 196) = 0.9650, $p=0.4907$ | N/A | 8 | | | |
| | **H** | Illustration showing location of intra-nucleus accumbens cannula placement | | | | | | | |
| | **I** | Experimental timeline of PD149163 (intra-nucleus accumbens)-induced hypothermia studies | | | | | | | |
| | **J** | Illustration of ink-verified intra-nucleus accumbens cannula placements for mice in experimental panel K | | | | | | | |
| | **K** | PD149163-induced Hypothermia in WT mice – Body Temperature Pre and Post Intra-NAc Treatment | Two-way repeated measures ANOVA | $F_{Time}$ (1, 24) = 227.1, $p<0.0001$<br><br>$F_{Treatment}$ (3, 24) = 64.23, $p<0.0001$<br><br>$F_{Interaction}$ (3, 24) = 62.95, $p<0.0001$ | ***p<0.001, WT Vehicle vs. WT PD149163 | 8 | | | |
| | | PD149163-induced Hypothermia in NTSR1$^{-/-}$ mice – Body Temperature Pre and Post Intra-NAc Treatment | Two-way repeated measures ANOVA | $F_{Time}$ (1, 10) = 2.455<br><br>$F_{Treatment}$ (1, 10) = 1.609<br><br>$F_{Interaction}$ (1, 10) = 4.277 | N/A | 6 | | | |
| | **L** | Experimental timeline of studies evaluating the effect of SBI compounds (intra-nucleus accumbens) on PD149163 (intra-nucleus accumbens)-induced hypothermia | | | | | | | |
| | **M** | Illustration of ink-verified intra-nucleus accumbens cannula placements for mice in experimental panel N | | | | | | | |
| | **N** | SBI-593 Attenuation of Intra-NAc PD149163-induced Hypothermia – Time Course of Body Temperature | Two-way repeated measures ANOVA, Time 30-480 min post PD149163 treatment | $F_{Time}$ (1.979, 35.62) = 15.88, $p<0.0001$<br><br>$F_{Treatment}$ (2, 19) = 6.298, $p=0.0080$<br><br>$F_{Interaction}$ (16, 144) = 3.909, $p<0.0001$ | *p<0.01, **p<0.0001, SBI-553 vs SBI-593 vs Vehicle | 6-9 | | | |

| | | | | | |
|---|---|---|---|---|---|
| **O** | Area Under the Curve | One-way nonparametric ANOVA followed by Dunn's multiple comparisons and Kruskal-Wallin tests | Kruskal-Wallis statistic= 6.304, p = 0.0363 | *p = 0.0341, Vehicle vs SBI-553<br><br>p = 0.1897, Vehicle vs SBI-593 | 6-9 |
| **P** | Complete vs Incomplete Blockade of PD149163-induced Hypothermia | Chi-Square, Two-sided | Chi-square, df = 100.8, 1<br><br>Z = 10.04 | *p<0.0001, SBI-553 vs SBI-593 | 6-9 |

One-way ANOVA; Dunnett's multiple comparison test; Kruskal-Wallis test; CI, confidence interval; NT, neurotensin; SBI-553, SBI-0654553; $IC_{50}$, half-maximal inhibitory concentration ND, not defined

**Table S7. Compound identification and curve parameters presented in Supplemental Figure 4.**

| Fig | Compound Number | Compound Name | Structure | | Parameter | Mean (95% CI) | Units |
|---|---|---|---|---|---|---|---|
| **S4** | 1 | SBI-0654498 | 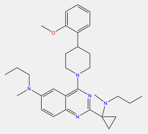 | Gαq | $IC_{50}$ | ~ 0.08349 (ND) | M |
| | | | | | Bottom | 0.000 | % Inhibition |
| | | | | $G_{\alpha}i1$ | $IC_{50}$ | ~ 0.2478 (ND) | M |
| | | | | | Bottom | 0.000 | % Inhibition |
| | | | | GαoA | $IC_{50}$ | 0.0001010 | M |
| | | | | | Bottom | 0.000 | % Inhibition |
| | | | | Gα12 | $IC_{50}$ | ND | M |
| | | | | | Bottom | 0.000 | % Inhibition |
| | | | | β-arrestin 2 | $EC_{50}$ | ~ 0.3865 (ND) | M |
| | | | | | Top | 107.938 | % Agonism |
| | 2 | SBI-0653810 | 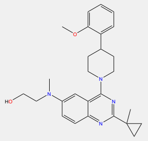 | Gαq | $IC_{50}$ | 2.552e-006 | M |
| | | | | | Bottom | 69.362 | % Inhibition |
| | | | | $G_{\alpha}i1$ | $IC_{50}$ | 1.231e-006 | M |
| | | | | | Bottom | 31.794 | % Inhibition |
| | | | | GαoA | $IC_{50}$ | ND | M |
| | | | | | Bottom | 0.000 | % Inhibition |
| | | | | Gα12 | $IC_{50}$ | ND | M |
| | | | | | Bottom | 0.000 | % Inhibition |
| | | | | β-arrestin 2 | $EC_{50}$ | 6.596e-006 | M |
| | | | | | Top | 105.957 | % Agonism |
| | 3 | SBI-0655884 | 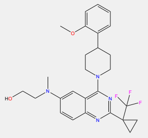 | Gαq | $IC_{50}$ | 6.277e-007 | M |
| | | | | | Bottom | 78.481 | % Inhibition |
| | | | | $G_{\alpha}i1$ | $IC_{50}$ | 1.852e-007 | M |
| | | | | | Bottom | 29.862 | % Inhibition |
| | | | | GαoA | $IC_{50}$ | 2.188e-006 | M |
| | | | | | Bottom | -21.482 | % Inhibition |
| | | | | Gα12 | $IC_{50}$ | ND | M |
| | | | | | Bottom | 0.000 | % Inhibition |
| | | | | β-arrestin 2 | $EC_{50}$ | 3.371e-006 | M |
| | | | | | Top | 112.778 | % Agonism |
| | 4 | SBI-0647348 | 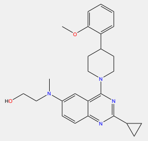 | Gαq | $IC_{50}$ | 7.041e-006 | M |
| | | | | | Bottom | 94.046 | % Inhibition |
| | | | | $G_{\alpha}i1$ | $IC_{50}$ | 1.143e-006 | M |
| | | | | | Bottom | 43.763 | % Inhibition |
| | | | | GαoA | $IC_{50}$ | 9.264e-007 | M |
| | | | | | Bottom | 0.000 | % Inhibition |
| | | | | Gα12 | $IC_{50}$ | ~ 1.353e-013 (ND) | M |
| | | | | | Bottom | 0.000 | % Inhibition |
| | | | | β-arrestin 2 | $EC_{50}$ | 6.763e-006 | M |
| | | | | | Top | 113.437 | % Agonism |
| | 5 | SBI-0654503 | 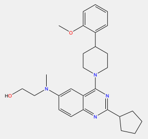 | Gαq | $IC_{50}$ | 1.021e-005 | M |
| | | | | | Bottom | 116.431 | % Inhibition |
| | | | | $G_{\alpha}i1$ | $IC_{50}$ | 3.52e-005 | M |
| | | | | | Bottom | 52.913 | % Inhibition |
| | | | | GαoA | $IC_{50}$ | ~ 0.03047 (ND) | M |
| | | | | | Bottom | 0.000 | % Inhibition |
| | | | | Gα12 | $IC_{50}$ | 8.411e-007 | M |
| | | | | | Bottom | 0.000 | % Inhibition |
| | | | | β-arrestin 2 | $EC_{50}$ | 1.510e-005 | M |

| | | | | | | |
|---|---|---|---|---|---|---|
| | | | | Top | 115.302 | % Agonism |
| 6 | SBI-0656152 | 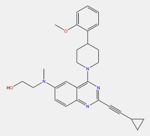 | Gαq | $IC_{50}$ | 5.683e-006 | M |
| | | | | Bottom | 83.075 | % Inhibition |
| | | | $G_{\alpha}i1$ | $IC_{50}$ | 2.09e-006 | M |
| | | | | Bottom | 34.615 | % Inhibition |
| | | | GαoA | $IC_{50}$ | 2.560e-008 | M |
| | | | | Bottom | 0.000 | % Inhibition |
| | | | Gα12 | $IC_{50}$ | ND | M |
| | | | | Bottom | 0.000 | % Inhibition |
| | | | β-arrestin 2 | $EC_{50}$ | 1.305e-005 | M |
| | | | | Top | 106.223 | % Agonism |
| 7 | SBI-0656154 | 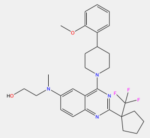 | Gαq | $IC_{50}$ | 8.821e-007 | M |
| | | | | Bottom | 63.674 | % Inhibition |
| | | | $G_{\alpha}i1$ | $IC_{50}$ | 6.51e-008 | M |
| | | | | Bottom | 2.900 | % Inhibition |
| | | | GαoA | $IC_{50}$ | 2.956e-006 | M |
| | | | | Bottom | 0.000 | % Inhibition |
| | | | Gα12 | $IC_{50}$ | 2.460e-008 | M |
| | | | | Bottom | 0.000 | % Inhibition |
| | | | β-arrestin 2 | $EC_{50}$ | 1.796e-005 | M |
| | | | | Top | 91.881 | % Agonism |
| 8 | SBI-0656032 | 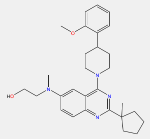 | Gαq | $IC_{50}$ | 1.17e-005 | M |
| | | | | Bottom | 96.304 | % Inhibition |
| | | | $G_{\alpha}i1$ | $IC_{50}$ | 9.70e-006 | M |
| | | | | Bottom | 41.492 | % Inhibition |
| | | | GαoA | $IC_{50}$ | 7.474e-009 | M |
| | | | | Bottom | 0.000 | % Inhibition |
| | | | Gα12 | $IC_{50}$ | ND | M |
| | | | | Bottom | 0.000 | % Inhibition |
| | | | β-arrestin 2 | $EC_{50}$ | 4.022e-005 | M |
| | | | | Top | 83.462 | % Agonism |
| 9 | SBI-0657449 | 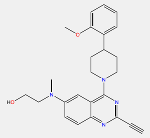 | Gαq | $IC_{50}$ | ~ 0.02862 (ND) | M |
| | | | | Bottom | 33.028 | % Inhibition |
| | | | $G_{\alpha}i1$ | $IC_{50}$ | ~ 0.01649 (ND) | M |
| | | | | Bottom | 0.000 | % Inhibition |
| | | | GαoA | $IC_{50}$ | 8.780e-007 | M |
| | | | | Bottom | 0.000 | % Inhibition |
| | | | Gα12 | $IC_{50}$ | ND | M |
| | | | | Bottom | 0.000 | % Inhibition |
| | | | β-arrestin 2 | $EC_{50}$ | 0.0001746 | M |
| | | | | Top | 37.132 | % Agonism |
| 10 | SBI-0656588 | 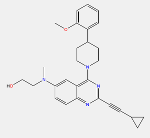 | Gαq | $IC_{50}$ | ~ 0.03411 (ND) | M |
| | | | | Bottom | 32.591 | % Inhibition |
| | | | $G_{\alpha}i1$ | $IC_{50}$ | ~ 0.02671 (ND) | M |
| | | | | Bottom | 0.000 | % Inhibition |
| | | | GαoA | $IC_{50}$ | ~ 0.03883 (ND) | M |
| | | | | Bottom | 0.000 | % Inhibition |
| | | | Gα12 | $IC_{50}$ | ND | M |
| | | | | Bottom | 0.000 | % Inhibition |
| | | | β-arrestin 2 | $EC_{50}$ | ~ 0.01559 (ND) | M |
| | | | | Top | 36.578 | % Agonism |
| 11 | SBI-0647347 | | Gαq | $IC_{50}$ | 1.222e-006 | M |
| | | | | Bottom | 103.917 | % Inhibition |
| | | | $G_{\alpha}i1$ | $IC_{50}$ | 5.54e-007 | M |

| | | | | | | |
|---|---|---|---|---|---|---|
| | | 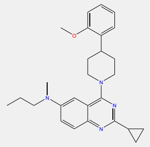 | | Bottom | 40.965 | % Inhibition |
| | | | GαoA | $IC_{50}$ | ~ 0.05441 (ND) | M |
| | | | | Bottom | 0.000 | % Inhibition |
| | | | Gα12 | $IC_{50}$ | 2.071e-008 | M |
| | | | | Bottom | 0.000 | % Inhibition |
| | | | β-arrestin 2 | $EC_{50}$ | 2.166e-006 | M |
| | | | | Top | 137.067 | % Agonism |
| 12 | SBI-0654552 | 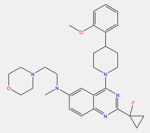 | Gαq | $IC_{50}$ | 3.718e-006 | M |
| | | | | Bottom | 78.189 | % Inhibition |
| | | | $G_{\alpha}$i1 | $IC_{50}$ | 8.747e-008 | M |
| | | | | Bottom | 15.150 | % Inhibition |
| | | | GαoA | $IC_{50}$ | 1.331e-005 | M |
| | | | | Bottom | 0.000 | % Inhibition |
| | | | Gα12 | $IC_{50}$ | 3.477e-008 | M |
| | | | | Bottom | 0.000 | % Inhibition |
| | | | β-arrestin 2 | $EC_{50}$ | 7.457e-006 | M |
| | | | | Top | 83.938 | % Agonism |
| 13 | SBI-0656026 | 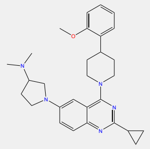 | Gαq | $IC_{50}$ | 4.98e-005 | M |
| | | | | Bottom | 37.333 | % Inhibition |
| | | | $G_{\alpha}$i1 | $IC_{50}$ | 1.35e-005 | M |
| | | | | Bottom | 33.350 | % Inhibition |
| | | | GαoA | $IC_{50}$ | 0.7238 (ND) | M |
| | | | | Bottom | 0.000 | % Inhibition |
| | | | Gα12 | $IC_{50}$ | 2.099e-007 | M |
| | | | | Bottom | 0.000 | % Inhibition |
| | | | β-arrestin 2 | $EC_{50}$ | ~ 0.06570 (ND) | M |
| | | | | Top | 40.578 | % Agonism |
| 14 | SBI-0656912 | 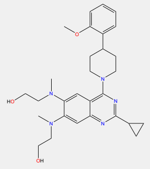 | Gαq | $IC_{50}$ | 1.454e-005 | M |
| | | | | Bottom | 0.000 | % Inhibition |
| | | | $G_{\alpha}$i1 | $IC_{50}$ | ~ 0.01075 (ND) | M |
| | | | | Bottom | 0.000 | % Inhibition |
| | | | GαoA | $IC_{50}$ | 3.941e-009 | M |
| | | | | Bottom | 0.000 | % Inhibition |
| | | | Gα12 | $IC_{50}$ | ~ 2.532e-013 (ND) | M |
| | | | | Bottom | 0.000 | % Inhibition |
| | | | β-arrestin 2 | $EC_{50}$ | 8.765e-005 | M |
| | | | | Top | 0.000 | % Agonism |
| 15 | SBI-0655874 | 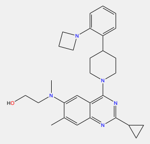 | Gαq | $IC_{50}$ | ~ 0.01032 (ND) | M |
| | | | | Bottom | 32.300 | % Inhibition |
| | | | $G_{\alpha}$i1 | $IC_{50}$ | 5.98e-006 | M |
| | | | | Bottom | 23.463 | % Inhibition |
| | | | GαoA | $IC_{50}$ | ND | M |
| | | | | Bottom | 0.000 | % Inhibition |
| | | | Gα12 | $IC_{50}$ | 1.505e-007 | M |
| | | | | Bottom | 0.000 | % Inhibition |
| | | | β-arrestin 2 | $EC_{50}$ | 9.726e-005 | M |
| | | | | Top | 65.583 | % Agonism |
| 16 | SBI-0657755 | 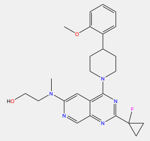 | Gαq | $IC_{50}$ | 1.464e-005 | M |
| | | | | Bottom | 22.398 | % Inhibition |
| | | | $G_{\alpha}$i1 | $IC_{50}$ | 2.58e-005 | M |
| | | | | Bottom | 24.291 | % Inhibition |
| | | | GαoA | $IC_{50}$ | 8.321e-005 | M |
| | | | | Bottom | 0.000 | % Inhibition |
| | | | Gα12 | $IC_{50}$ | 0.8450 (ND) | M |

| | | | | | | |
|---|---|---|---|---|---|---|
| | | | | Bottom | 0.000 | % Inhibition |
| | | | β-arrestin 2 | EC$_{50}$ | 1.300e-005 | M |
| | | | | Top | 95.382 | % Agonism |
| 17 | SBI-0657962 | 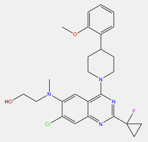 | Gαq | IC$_{50}$ | 5.558e-006 | M |
| | | | | Bottom | 37.252 | % Inhibition |
| | | | G$_{\alpha}$i1 | IC$_{50}$ | 1.28e-006 | M |
| | | | | Bottom | 20.465 | % Inhibition |
| | | | GαoA | IC$_{50}$ | 1.865e-007 | M |
| | | | | Bottom | 0.000 | % Inhibition |
| | | | Gα12 | IC$_{50}$ | 6.220e-009 | M |
| | | | | Bottom | 0.000 | % Inhibition |
| | | | β-arrestin 2 | EC$_{50}$ | 1.754e-005 | M |
| | | | | Top | 83.744 | % Agonism |
| 18 | SBI-0646912 | 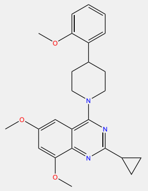 | Gαq | IC$_{50}$ | 6.949e-005 | M |
| | | | | Bottom | 43.057 | % Inhibition |
| | | | G$_{\alpha}$i1 | IC$_{50}$ | 1.448e-005 | M |
| | | | | Bottom | 42.900 | % Inhibition |
| | | | GαoA | IC$_{50}$ | ND | M |
| | | | | Bottom | 0.000 | % Inhibition |
| | | | Gα12 | IC$_{50}$ | 2.807e-009 | M |
| | | | | Bottom | 0.000 | % Inhibition |
| | | | β-arrestin 2 | EC$_{50}$ | ~ 4.830e-014 (ND) | M |
| | | | | Top | 59.258 | % Agonism |
| 19 | SBI-0656149 | 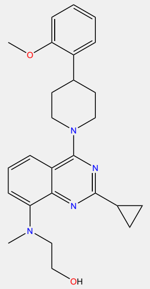 | Gαq | IC$_{50}$ | 3.665e-014 | M |
| | | | | Bottom | 0.000 | % Inhibition |
| | | | G$_{\alpha}$i1 | IC$_{50}$ | 4.89e-004 | M |
| | | | | Bottom | 11.762 | % Inhibition |
| | | | GαoA | IC$_{50}$ | 2.752e-008 | M |
| | | | | Bottom | 0.000 | % Inhibition |
| | | | Gα12 | IC$_{50}$ | ND | M |
| | | | | Bottom | 0.000 | % Inhibition |
| | | | β-arrestin 2 | EC$_{50}$ | ~ 0.05446 (ND) | M |
| | | | | Top | 0.000 | % Agonism |
| 20 | SBI-0656571 | 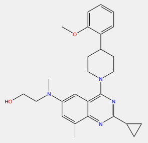 | Gαq | IC$_{50}$ | 6.489e-006 | M |
| | | | | Bottom | 92.464 | % Inhibition |
| | | | G$_{\alpha}$i1 | IC$_{50}$ | 2.57e-006 | M |
| | | | | Bottom | 74.566 | % Inhibition |
| | | | GαoA | IC$_{50}$ | ~ 0.01347 (ND) | M |
| | | | | Bottom | 0.000 | % Inhibition |
| | | | Gα12 | IC$_{50}$ | ND | M |
| | | | | Bottom | 0.000 | % Inhibition |
| | | | β-arrestin 2 | EC$_{50}$ | 9.598e-006 | M |
| | | | | Top | 89.642 | % Agonism |
| 21 | SBI-0657754 | 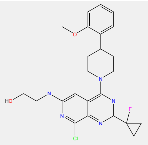 | Gαq | IC$_{50}$ | 1.865e-006 | M |
| | | | | Bottom | 99.355 | % Inhibition |
| | | | G$_{\alpha}$i1 | IC$_{50}$ | 7.65e-007 | M |
| | | | | Bottom | 45.646 | % Inhibition |
| | | | GαoA | IC$_{50}$ | 1.046e-007 | M |
| | | | | Bottom | 0.000 | % Inhibition |
| | | | Gα12 | IC$_{50}$ | ND | M |
| | | | | Bottom | 0.000 | % Inhibition |
| | | | β-arrestin 2 | EC$_{50}$ | 3.643e-006 | M |
| | | | | Top | 93.819 | % Agonism |
| 22 | SBI-0657750 | | Gαq | IC$_{50}$ | 1.908e-006 | M |

| | | | | | | |
|---|---|---|---|---|---|---|
| | | 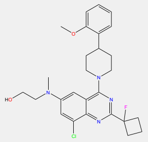 | | Bottom | 107.544 | % Inhibition |
| | | | $G_{\alpha}i1$ | $IC_{50}$ | 3.963e-007 | M |
| | | | | Bottom | 33.778 | % Inhibition |
| | | | GαoA | $IC_{50}$ | ND | M |
| | | | | Bottom | 0.000 | % Inhibition |
| | | | Gα12 | $IC_{50}$ | 6.747e-007 | M |
| | | | | Bottom | 0.000 | % Inhibition |
| | | | β-arrestin 2 | $EC_{50}$ | 3.465e-006 | M |
| | | | | Top | 121.572 | % Agonism |
| 23 | MLS-0463341 | 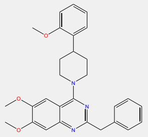 | Gαq | $IC_{50}$ | ~ 0.03469 (ND) | M |
| | | | | Bottom | 0.000 | % Inhibition |
| | | | $G_{\alpha}i1$ | $IC_{50}$ | 7.906e-005 | M |
| | | | | Bottom | 10.907 | % Inhibition |
| | | | GαoA | $IC_{50}$ | 2.182e-007 | M |
| | | | | Bottom | 0.000 | % Inhibition |
| | | | Gα12 | $IC_{50}$ | 1.042e-005 | M |
| | | | | Bottom | 0.000 | % Inhibition |
| | | | β-arrestin 2 | $EC_{50}$ | 0.0001577 | M |
| | | | | Top | 47.155 | % Agonism |
| 24 | MLS-0469654 | 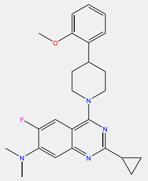 | Gαq | $IC_{50}$ | ND | M |
| | | | | Bottom | 0.000 | % Inhibition |
| | | | $G_{\alpha}i1$ | $IC_{50}$ | ND | M |
| | | | | Bottom | 5.044 | % Inhibition |
| | | | GαoA | $IC_{50}$ | 1.884e-007 | M |
| | | | | Bottom | 0.000 | % Inhibition |
| | | | Gα12 | $IC_{50}$ | 1.677e-005 | M |
| | | | | Bottom | 0.000 | % Inhibition |
| | | | β-arrestin 2 | $EC_{50}$ | ~ 0.06534 (ND) | M |
| | | | | Top | 22.681 | % Agonism |
| 25 | SBI-0646238 | 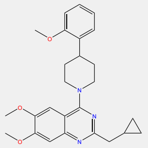 | Gαq | $IC_{50}$ | ND | M |
| | | | | Bottom | 0.000 | % Inhibition |
| | | | $G_{\alpha}i1$ | $IC_{50}$ | 1.58e-005 | M |
| | | | | Bottom | 6.175 | % Inhibition |
| | | | GαoA | $IC_{50}$ | 1.909e-007 | M |
| | | | | Bottom | 0.000 | % Inhibition |
| | | | Gα12 | $IC_{50}$ | 3.923e-006 | M |
| | | | | Bottom | 0.000 | % Inhibition |
| | | | β-arrestin 2 | $EC_{50}$ | ~ 0.2146 (ND) | M |
| | | | | Top | 29.945 | % Agonism |
| 26 | SBI-0646593 | 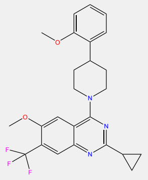 | Gαq | $IC_{50}$ | 3.905e-006 | M |
| | | | | Bottom | 32.449 | % Inhibition |
| | | | $G_{\alpha}i1$ | $IC_{50}$ | 1.05e-006 | M |
| | | | | Bottom | 23.793 | % Inhibition |
| | | | GαoA | $IC_{50}$ | 4.370e-006 | M |
| | | | | Bottom | 0.000 | % Inhibition |
| | | | Gα12 | $IC_{50}$ | 1.634e-007 | M |
| | | | | Bottom | 0.000 | % Inhibition |
| | | | β-arrestin 2 | $EC_{50}$ | 4.750e-006 | M |
| | | | | Top | 75.422 | % Agonism |
| 27 | SBI-0646782 | | Gαq | $IC_{50}$ | ND | M |
| | | | | Bottom | 0.000 | % Inhibition |
| | | | $G_{\alpha}i1$ | $IC_{50}$ | 2.03e-006 | M |
| | | | | Bottom | 1.059 | % Inhibition |
| | | | GαoA | $IC_{50}$ | 5.025e-008 | M |

| | | | | | | |
|---|---|---|---|---|---|---|
| | | 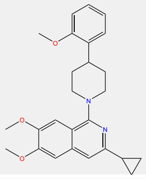 | | Bottom | 0.000 | % Inhibition |
| | | | Gα12 | $IC_{50}$ | 7.059e-007 | M |
| | | | | Bottom | 0.000 | % Inhibition |
| | | | β-arrestin 2 | $EC_{50}$ | 1.076e-005 | M |
| | | | | Top | 9.918 | % Agonism |
| 28 | SBI-0647342 | 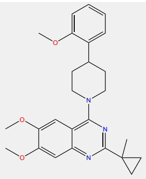 | Gαq | $IC_{50}$ | 6.659e-006 | M |
| | | | | Bottom | 60.927 | % Inhibition |
| | | | $G_{\alpha}$i1 | $IC_{50}$ | 1.48e-006 | M |
| | | | | Bottom | 33.716 | % Inhibition |
| | | | GαoA | $IC_{50}$ | 5.841e-006 | M |
| | | | | Bottom | 41.906 | % Inhibition |
| | | | Gα12 | $IC_{50}$ | 1.036e-005 | M |
| | | | | Bottom | 0.000 | % Inhibition |
| | | | β-arrestin 2 | $EC_{50}$ | 7.563e-006 | M |
| | | | | Top | 94.130 | % Agonism |
| 29 | SBI-0646503 | 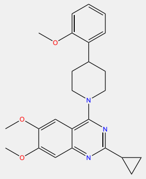 | Gαq | $IC_{50}$ | 1.021e-005 | M |
| | | | | Bottom | 58.928 | % Inhibition |
| | | | $G_{\alpha}$i1 | $IC_{50}$ | 5.58e-006 | M |
| | | | | Bottom | 55.933 | % Inhibition |
| | | | GαoA | $IC_{50}$ | ~ 0.03047 (ND) | M |
| | | | | Bottom | 24.532 | % Inhibition |
| | | | Gα12 | $IC_{50}$ | ND | M |
| | | | | Bottom | 0.000 | % Inhibition |
| | | | β-arrestin 2 | $EC_{50}$ | 1.298e-005 | M |
| | | | | Top | 70.781 | % Agonism |

CI, confidence interval; $EC_{50}$, half-maximal effective concentration; $IC_{50}$, half-maximal inhibitory concentration; ND, not defined
